# Gateway analysis reveals transient molecular programs at cell-fate transitions

**DOI:** 10.64898/2026.03.12.711328

**Authors:** Hu Cang, Sha Sun

## Abstract

Single-cell atlases have transformed our view of cell identity, but they still struggle to resolve the transient states that accompany cell-fate change. These rare interface cells are outnumbered by the stable populations on either side. As a result, the molecular programs that accompany lineage transitions are hard to detect. We developed gateway analysis, a framework that identifies cells at fate boundaries and pinpoints bell and valley genes that peak or dip there. It defines cell neighborhoods from binary mutual information (BMI) in gene on/off patterns and preserves that structure in a regularized latent model. Across four single-cell atlases spanning reprogramming, gastrulation, pancreatic endocrinogenesis, and kidney injury (3,696–126,578 cells), gateway analysis recovered rare boundary populations and the transient gene programs that distinguish them from their flanking states. These signals were missed by standard comparisons of stable endpoint states. They included epithelial remodeling at the late MET-to-iPSC interface, a gate-versus-basin partition of gastrulation regulators, a transient Cck-enriched peak at endocrine hub entry together with a candidate BH4-associated signal at a shared endocrine hilltop, and a proteostasis program at the kidney injury boundary. Orthogonal support from optimal-transport fate probabilities, known markers, and published perturbation phenotypes indicate that gateway cells mark bona fide biological transition intervals. Gateway analysis therefore provides a practical framework for detecting rare transition-state cells and the bell/valley genes that define them in single-cell atlases.

## 1 Introduction

Single-cell atlases have transformed the study of development, regeneration, and disease by resolving cellular states at unprecedented scale. Yet they are much better at cataloging stable endpoints than at capturing the brief states through which cells change fate. Those fleeting states are often mechanistically important. They mark where one regulatory program is dismantled and another is installed^1–2^. Defining the molecular logic of those boundary states remains a central goal of single-cell analysis.

Fate decisions arise in at least two recurrent landscape geometries. One is barrier crossing between attractor basins, as in somatic reprogramming^3^. The other is ridge bifurcation, in which a progenitor population resolves into alternative lineages, as in pancreatic endocrinogenesis^4^. In both settings, the cells closest to the decision boundary are expected to be rare, short-lived, and molecularly mixed. That makes them difficult to isolate even in large atlases, despite their likely importance for explaining how fate changes unfold.

A major reason such cells are missed is *abundance bias*. Most representation-learning and neighborhood-based methods define similarity from expression magnitude and then optimize global reconstruction or manifold structure across all cells^5–6^. In practice, large mature populations dominate the learned geometry. Rare interface cells are then absorbed into nearby endpoints. Trajectory methods can order cells along broad continua^7–9^, but they do not by themselves ensure that the small boundary population and its boundary-specific programs survive the embedding.

We reasoned that rare decision-state structure might be better captured by focusing on shared regulatory programs rather than absolute expression levels. We therefore define cell–cell similarity using binary mutual information (BMI), which measures whether two cells share coordinated gene-detection patterns beyond chance. Because BMI emphasizes on/off program structure while discarding expression magnitude, it can preserve local relationships for rare cells without weighting their contribution by population size. The resulting BMI graph is therefore more robust to abundance-correlated depth variation than conventional expression-based similarity.

We use this graph in two linked components. First, we introduce scAttnVI (single-cell Attention-weighted Variational Inference), a BMI-regularized variational autoencoder that preserves BMI-defined neighborhoods during latent learning through BMI-derived neighbor weights. Here, the “attention” weights are fixed BMI-derived coefficients rather than learned transformer-style attention.

Second, we perform gateway analysis on the resulting geometry to identify cells positioned between two flanking fates and to recover genes whose expression peaks specifically at that interface.

In principle, the workflow in Fig. 2 begins by binarizing gene detection and measuring how strongly pairs of cells share joint ON/OFF programs across genes. Those pairwise BMI values define a cell–cell graph, which is filtered against a null and converted into static attention weights. In the latent space, those weights act like local springs: they keep rare interface cells tethered to neighbors with shared programs rather than letting them be absorbed into abundant flanking populations (Fig. 2f).

**Fig. 1.**
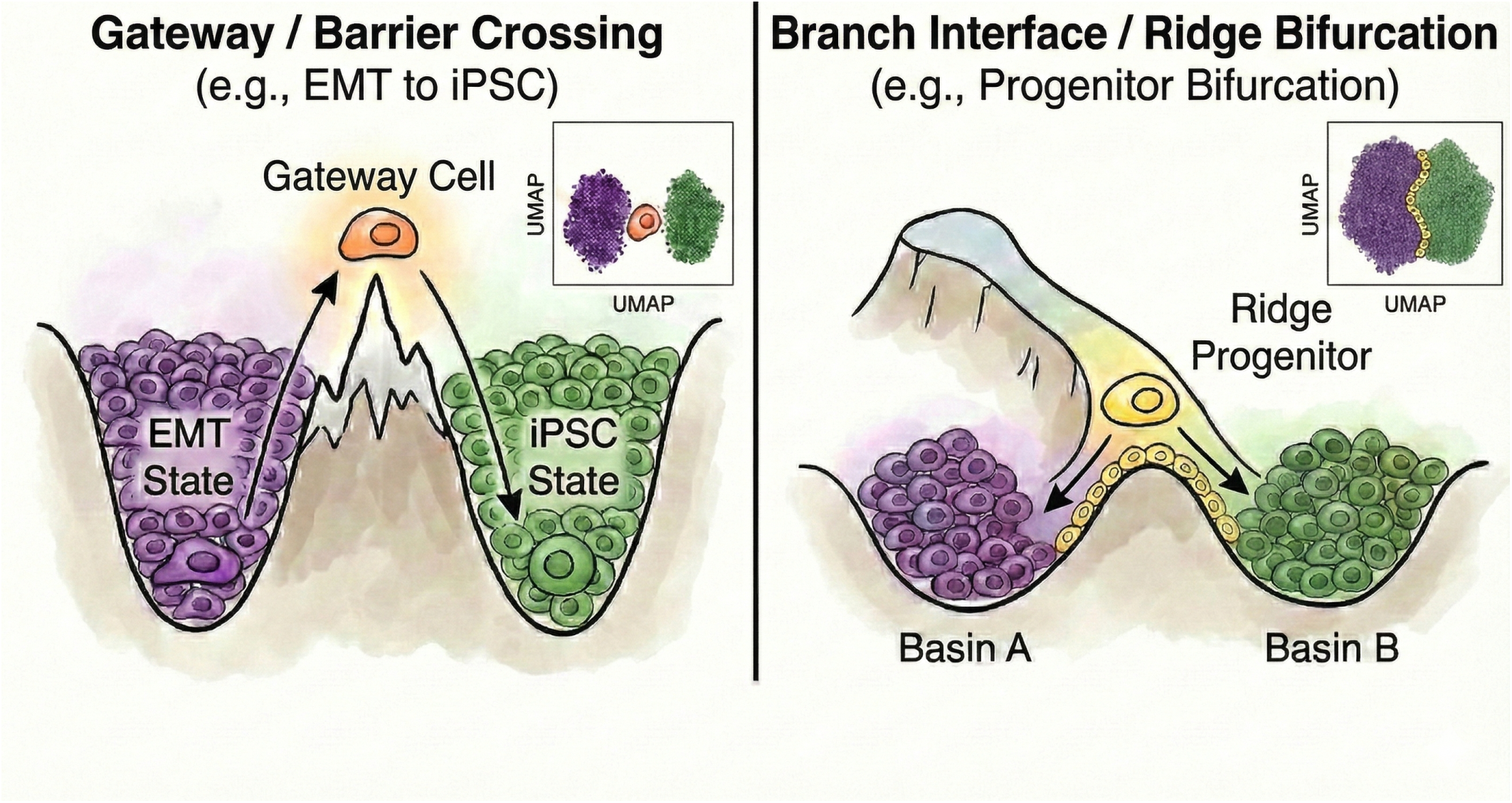
Two landscape topologies for fate-decision states. **Left**: Gateway / barrier crossing (e.g., MET to iPSC)—gateway cells (orange) sit at the saddle point between two stable basins. **Right**: Branch interface / ridge bifurcation (e.g., pancreatic endocrine bifurcation)—cells flow through a shared hilltop pre-commitment state from which two lineages diverge. Gateway scoring on the BMI graph detects transition-state cells in both geometries; each boundary carries its own molecular signature (bell and valley genes), distinct from either flanking fate.

**Fig. 2.**
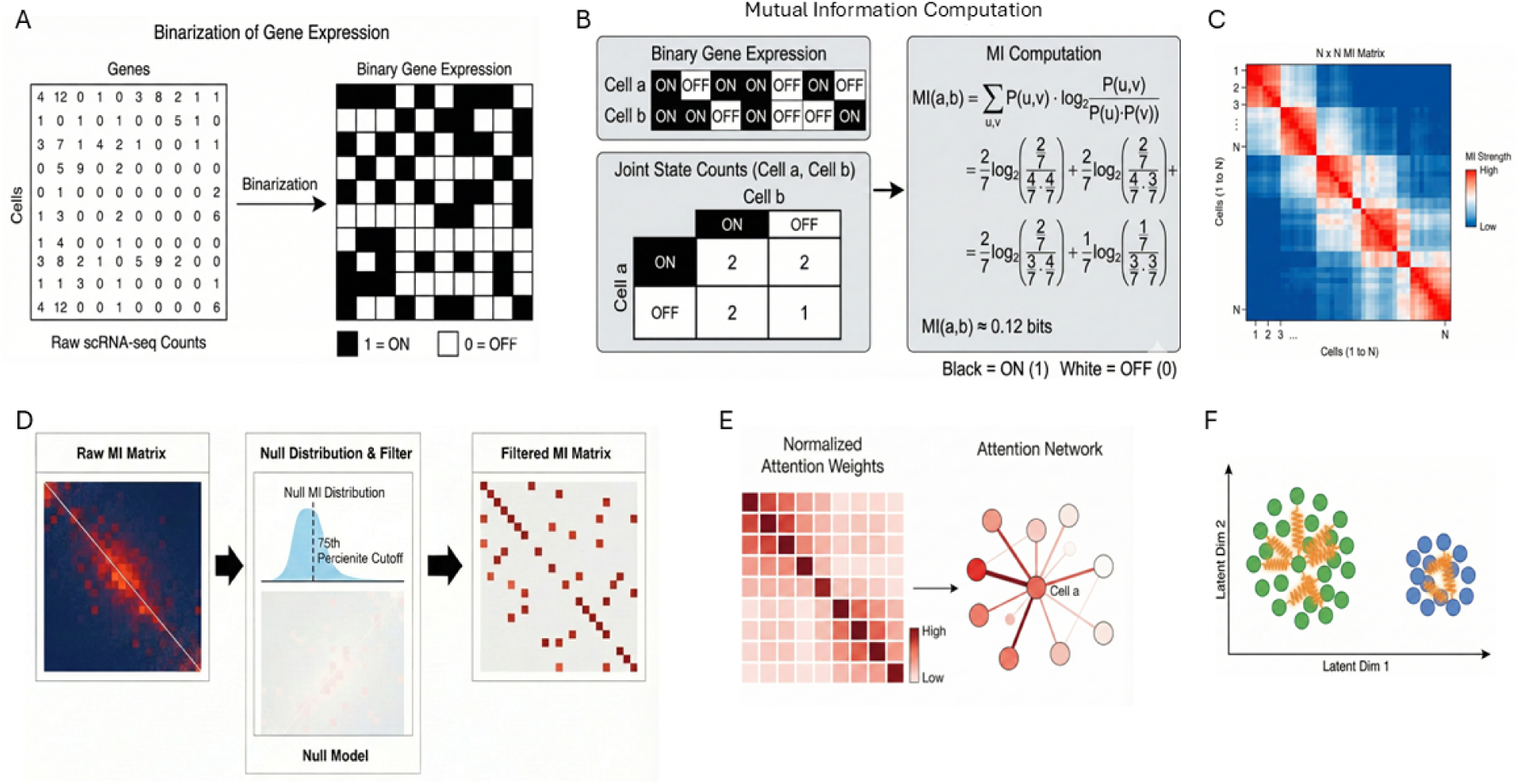
The BMI computational pipeline. **(a)** Gene expression is binarized (ON/OFF at count threshold *τ* = 1). **(b)** Binary mutual information is calculated for each cell pair from the joint detected/not-detected patterns across genes; illustrated with a minimal 2-cell, 2-gene example. **(c)** The full *N* × *N* MI matrix captures pairwise cell–cell similarity. **(d)** A null model (column-permuted MI) defines a significance threshold; sub-threshold entries are zeroed to produce a sparse, filtered MI graph. **(e)** Normalized static attention weights derived from the filtered MI graph define each cell’s neighborhood; masked softmax ensures disconnected neighbors receive exactly zero weight. **(f)** The BMI-regularized latent space: attention-derived springs (orange) anchor rare populations (blue) to their coexpression neighbors, preventing the reconstruction loss from collapsing them into the majority cluster.

The aim is not to replace clustering or trajectory inference. It is to sharpen the local transition interval between biologically credible flanking populations. In this study, we do not infer those flanking annotations de novo. Instead, we start from the published cell-state annotations provided by each atlas and use gateway analysis to refine the interval between them.

We apply gateway analysis across four atlases spanning distinct biological settings and landscape geometries: mouse reprogramming (Waddington-OT^3^), mouse gastrulation^10^, endocrine pancreas development^4^, and acute kidney injury^11^. Across these systems, gateway cells reveal coherent bell/valley gene programs that endpoint comparisons do not recover. Their biological relevance is supported by independent evidence including fate probabilities, known regulators, and published perturbation phenotypes. Most notably, in pancreas the method resolves a shared endocrine pre-commitment interval within a multi-exit fate hub and separates shared hilltop genes from lineage-bias gradients. Together, these results position gateway analysis as a practical framework for detecting real transition populations and the boundary-specific programs that distinguish them in single-cell atlases.

## 2 Results

### 2.1 Binary mutual information defines abundance-robust cell neighborhoods

Reconstruction-based single-cell embeddings are optimized over all cells, so abundant populations dominate the learned geometry. For models such as scVI^6^, the objective is

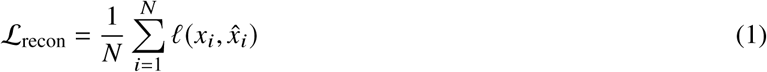

where *x*_*i*_ ∈ ℝ^*G*^ is the gene-count vector of cell *i*, *x̂*_*i*_ is its reconstruction from the latent embedding, and *N* is the total number of cells. If the dataset contains *K* populations of sizes *N*_1_, …, *N*_*K*_, the gradient decomposes as

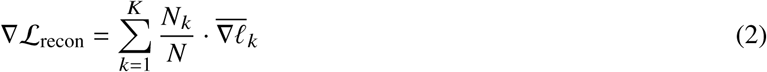

where 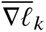 is the mean per-cell gradient from population *k*. Thus, large populations contribute proportionally larger updates, while rare interface populations are structurally underweighted. This abundance bias improves global fit but erodes the rare local neighborhoods needed to resolve transition states.

We address this by redefining cell–cell similarity in binary rather than quantitative terms. For each pair of cells, genes are classified as detected or not detected, and binary mutual information is computed across genes:

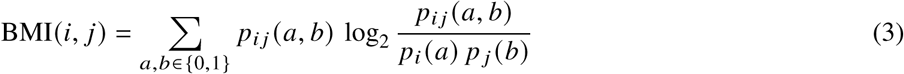

where high BMI indicates coordinated co-detection and co-silencing beyond chance. Because BMI captures shared regulatory-program structure rather than expression magnitude, a rare cell can remain close to its true biological neighbors even when it is surrounded by much larger populations.

We incorporate this structure into latent learning by adding a BMI-weighted neighborhood penalty to the generative objective:

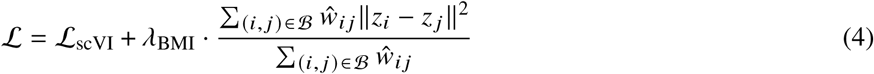

where *z*_*i*_ is the latent representation of cell *i*, *ŵ* _*i* *j*_ are static BMI-derived attention weights, and ℬ denotes within-batch cell pairs. We call the resulting model scAttnVI. The key effect is that cells are pulled toward partners defined by shared programs rather than by the global abundance of their cluster.

### 2.2 BMI preserves rare-population geometry and improves imputation

To test whether BMI preserves rare populations under controlled abundance stress, we used peripheral blood mononuclear cells (PBMCs)^12^, where cell identities are well established and one population can be selectively depleted while the rest of the atlas is held fixed.

We constructed a synthetic thinning experiment (Fig. 3). Starting from ∼12,000 cells containing 1,448 CD8 T cells, we reduced CD8 abundance to 50%, 20%, 10%, and 5% of its original count while keeping all other populations unchanged. PCA, scVI (reconstruction only), and scAttnVI (reconstruction + BMI prior; Supplementary Note 1; Extended Data Fig. 1) were retrained independently on each thinned dataset. All UMAP plots use the same reference PCA-based coordinates and matched clustering settings, so abundance is the only manipulated variable.

**Fig. 3.**
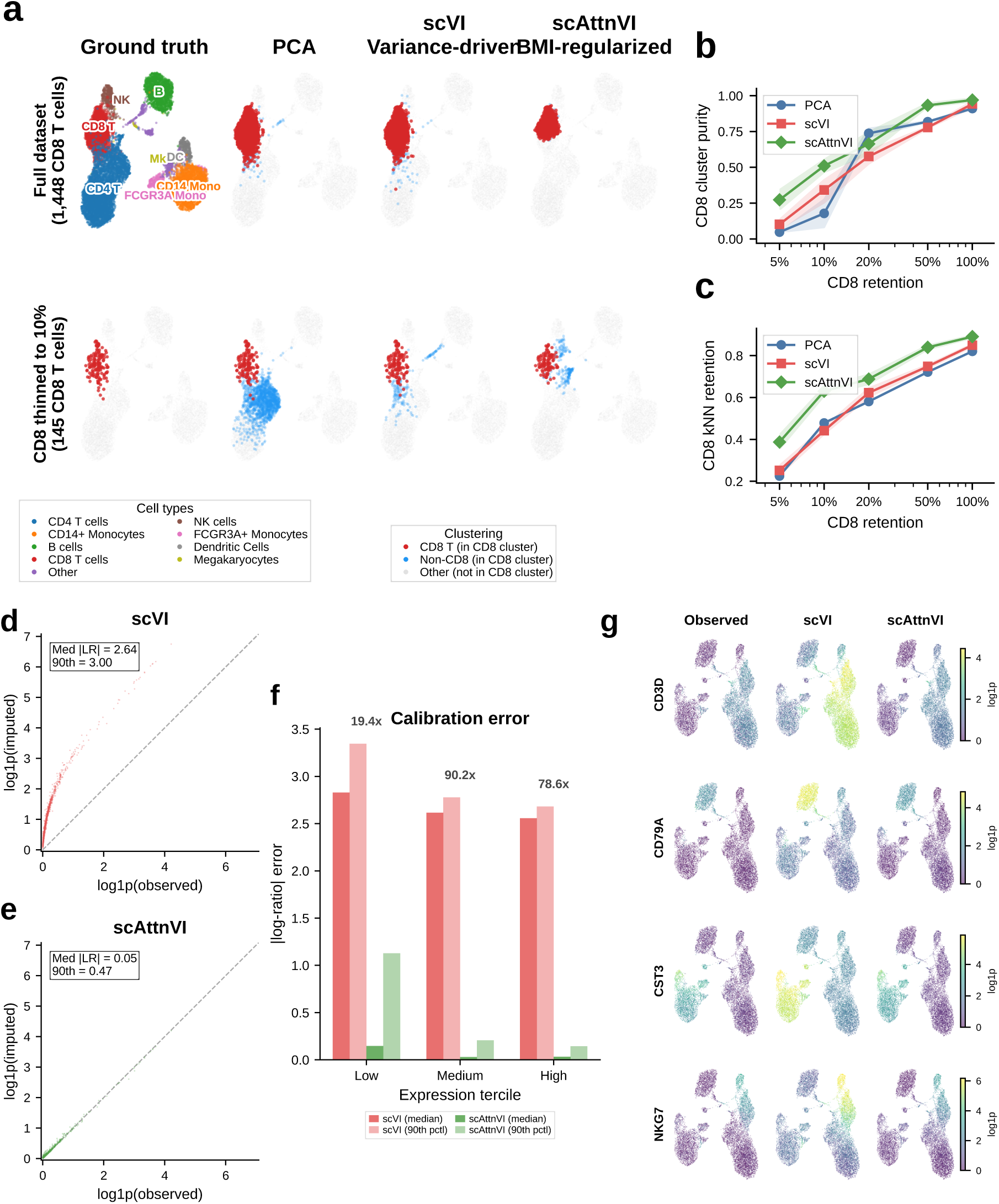
BMI regularization preserves rare-population geometry and improves imputation. **(a)** Top row: full dataset (1,448 CD8 T cells, 12.1%); bottom row: CD8 T cells thinned to 10% retention (145 cells) while all other populations are held fixed. Columns show the reference labels, PCA, scVI, and scAttnVI. At 10% retention, PCA and scVI merge CD8 cells into neighboring clusters, whereas scAttnVI maintains a distinct CD8 neighborhood. **(b)** CD8 cluster purity as a function of retention fraction (*n* = 5 independent runs with different random seeds; error bars: ±1 s.d.). At 10% retention, scAttnVI advantage is significant (Cohen’s *d*_*z*_ = 2.47, *P* = 0.005, paired *t*-test). **(c)** CD8 *k*NN retention: fraction of each CD8 cell’s *k* = 30 nearest neighbors that are true CD8 cells, capturing local geometric preservation. **(d)** scVI calibration: mean imputed versus mean observed expression (log1p scale) across all genes (median log ratio = 2.64). **(e)** scAttnVI calibration: near-perfect alignment along the identity line (median log ratio = 0.05). **(f)** Calibration error stratified by expression level; scAttnVI reduces error by 19–90-fold. **(g)** Marker gene expression on UMAP: observed, scVI-imputed, and scAttnVI-imputed expression for CD3D, CD79A, CST3, and NKG7.

At full retention, all three methods separated CD8 cells similarly (cluster purity 0.91–0.97; Fig. 3a). As CD8 cells became rare, PCA and scVI progressively merged them into adjacent populations, whereas scAttnVI preserved a distinct CD8 neighborhood. At 5% retention (72 CD8 cells among ∼10,600 total), scAttnVI maintained a purity of 0.273, compared with 0.102 for scVI and 0.047 for PCA (Fig. 3b). This behavior was stable across seeds and across a broad *λ*_BMI_ range (Extended Data Fig. 1).

This advantage was also evident at the local-neighborhood level. For each CD8 cell, we measured the fraction of its *k*-nearest neighbors (*k*NN) in latent space that were true CD8 cells (Fig. 3c). scAttnVI maintained higher *k*NN retention across all thinning levels, showing that BMI preserves the local geometry that reconstruction-only objectives sacrifice as rare populations shrink. In the context of gateway analysis, this is the critical property that minimizes the risk of rare boundary cells being absorbed into abundant mature states.

Improved geometry also improved denoising. When cell types are mixed in latent space, imputation spreads marker expression across populations and systematically inflates counts. scVI overestimated mean imputed expression by ∼14-fold (Fig. 3d), whereas scAttnVI restored near-linear agreement with observed expression (Fig. 3e,f) and recovered marker patterns with appropriate spatial restriction on the embedding (Fig. 3g). Thus, the same BMI prior that preserves rare-state neighborhoods also reduces the propagation of abundance bias into downstream imputation and transport-based analyses.

### 2.3 Gateway analysis captures barrier-crossing cells in mouse reprogramming

Schiebinger et al.^3^ introduced Waddington-OT (WOT), a probabilistic optimal-transport framework that reconstructs mouse embryonic fibroblast (MEF)-to-induced pluripotent stem cell (iPSC) reprogramming from more than 330,000 cells sampled at half-day intervals across days 0–18. WOT is an unusually stringent benchmark for gateway analysis because it provides not only a trajectory but also a computationally distinct fate estimate on the same atlas, the backward transport probability *P*(iPSC). Reprogramming does not follow a single path. Cells can progress through a mesenchymal-to-epithelial transition (MET) route toward iPSC or diverge into off-target stromal, trophoblast, and neural branches, with only a minority ultimately reaching pluripotency. The study profiled two conditions: serum (∼166,000 cells) and 2i/LIF (∼169,000 cells), where 2i/LIF combines MEK and GSK3*β* inhibitors with leukemia inhibitory factor^13^. In both settings, MET is the dominant route to iPSC, whereas 2i/LIF suppresses off-target fates and improves reprogramming efficiency.

WOT further showed that successful iPSC ancestors pass through a tight bottleneck: the fraction of cells on the iPSC trajectory contracts from roughly 40% at day 8.5 to only ∼1% in serum and ∼10% in 2i by days 10–11^3^. In landscape terms, this is a barrier-crossing event. Cells must leave a stable mesenchymal basin, traverse an unstable interface, and then enter the pluripotent state (Fig. 1).

A central output of WOT is the backward transport probability *P*(iPSC), which assigns each cell a quantitative estimate of how likely it is to ultimately become an iPSC. Cells undergoing MET show markedly higher *P*(iPSC), reinforcing the idea that MET is the main route to successful reprogramming, whereas stromal, trophoblast, and neural branches remain near zero. This makes WOT an unusually strong benchmark. It tells us where successful reprogramming is happening, and lets us ask whether gateway analysis can isolate the short-lived cells at that bottleneck.

We trained the BMI-regularized model (scAttnVI; Methods) separately on the serum and 2i datasets using the annotations from Schiebinger et al.^3^. The resulting latent spaces recovered the major WOT branches, with clear separation of iPSC, MET, stromal, trophoblast, and neural populations and smooth progression of day labels along the trajectory axis (Fig. 4a–d). When projected onto these embeddings, WOT backward transport probabilities showed a highly localized pattern (Fig. 4e,g). Cells with elevated *P*(iPSC) outside the iPSC cluster accumulated next to the MET–iPSC corridor, whereas off-target lineages remained largely devoid of signal. These non-iPSC cells with high *P*(iPSC) define the candidate gateway population.

**Fig. 4.**
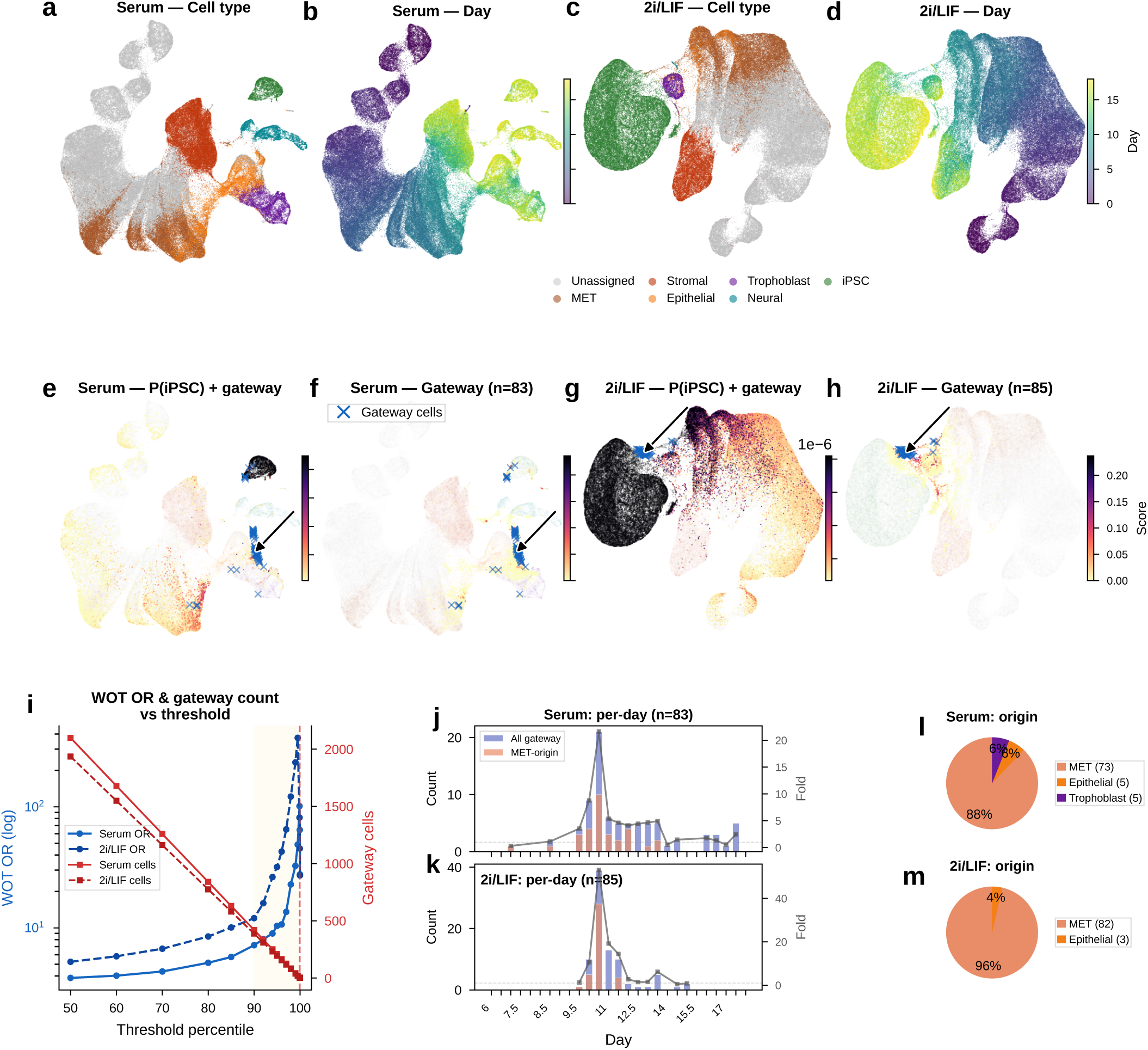
BMI gateway analysis captures gateway cells in MEF-to-iPSC reprogramming. **(a,c)** UMAP colored by WOT-annotated cell type (labels from Schiebinger et al.^3^). **(b,d)** UMAP colored by experimental day (0–18, viridis). **(e,g)** *P*(iPSC) heatmaps (magma_r); blue arrows indicate gateway cells. **(f,h)** Gateway-score overlays (magma_r); blue crosses mark gateway cells (serum *n*=83, 2i *n*=85). **(i)** WOT odds ratio (left axis) and gateway cell count (right axis) as a function of gateway score threshold percentile. Both conditions show sharply increasing enrichment above the 99th percentile; dashed line marks the 99.95th percentile threshold used throughout. **(j,k)** Per-day gateway cell count (bars) with MET-origin overlay (orange) and enrichment fold (right axis, circles). **(l,m)** Gateway cell origin composition. Pie charts show the highest-scoring non-iPSC source for each gateway cell.

The practical question is how to isolate the bottleneck cells themselves rather than all cells with high iPSC fate probability. One option is to threshold *P*(iPSC) directly. However, that strategy has two important limitations. First, *P*(iPSC) depends on WOT’s optimal-transport framework and therefore on densely sampled time-course data, which most single-cell studies do not have. A definition of gateway cells tied to *P*(iPSC) would therefore not generalize beyond this benchmark system. Second, *P*(iPSC) saturates within the iPSC cluster itself: newly arrived boundary cells and deeply committed iPSCs receive similarly high scores. That makes it poorly suited for distinguishing transitional cells on the iPSC side of the boundary. We therefore defined a gateway score that operates on *k*-nearest-neighbor (kNN) neighborhoods in the BMI-regularized latent space and requires only cell-type annotations.

#### 2.3.1 An active transition state at the reprogramming bottleneck

The bottleneck described by Schiebinger et al.^3^ implies that individual cells must cross a barrier between the dominant non-iPSC lineage and the iPSC state. Because this crossing is transient, barrier-crossing cells should remain connected to both fates while not yet being fully committed to either identity. To identify these cells, we computed a *gateway score* from kNN neighborhoods in the BMI-regularized latent space. For each cell *i*, we define a *fate pull* toward iPSC as the fraction of its *k* nearest neighbors annotated as iPSC:

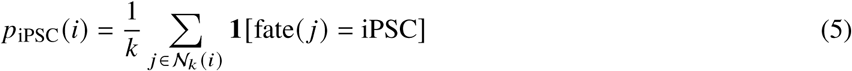

where *N*_*k*_ (*i*) is the *k*-nearest-neighbor set of cell *i* in latent space. We analogously compute the strongest pull toward any non-iPSC lineage, *p*_non-iPSC_(*i*) = max_*B*≠iPSC_ *p*_*B*_(*i*).

A cell deep inside the iPSC cluster has *p*_iPSC_ ≈ 1. A cell far from iPSCs has *p*_iPSC_ ≈ 0. A cell at the boundary has an intermediate value.

Cells at the interface should be pulled toward both fates. We therefore define the gateway score as:

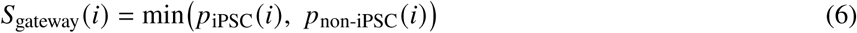

A cell embedded within one lineage scores near zero because one of the two pulls vanishes; only a cell connected to both fates scores high. Because the score is computed in the BMI-regularized latent space, it depends on cell-type annotations and on a geometry shaped by shared gene programs rather than by population size. The method therefore refines the interval between plausible annotated states rather than substituting for expert annotation itself.

The raw gateway score can include isolated high values caused by dropout noise or stochastic gene detection. We therefore applied personalized PageRank diffusion^14–15^ on a latent-space neighbor graph:

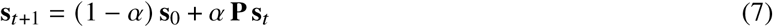

where **s**_0_ is the initial gateway score vector, **P** is the row-stochastic transition matrix. At each step, a cell’s score is updated as a weighted average of its latent-space neighbors while retaining a fraction (1 − *⍺*) of the original signal. This smooths the score landscape along the learned manifold without erasing spatial specificity. The raw score already captures the main biological signal; diffusion is an optional denoising step that modestly sharpens the spatial pattern while leaving most top-scoring cells shared between the raw and diffused analyses (Extended Data Fig. 2a–e).

Projected onto the UMAP, the diffused score formed a narrow ridge at the MET–iPSC interface in both conditions (Fig. 4f,h). The score distribution was highly skewed: most cells scored near zero, with a thin high-value tail (Fig. 4i, left). Increasing score stringency reduced the number of selected cells but increased their median WOT backward *P*(iPSC) (Fig. 4i, right), indicating that higher gateway scores mark cells with greater independent iPSC fate potential. Using a conservative operating cutoff, we identified a compact gateway core of about 80 cells per condition, or roughly 0.05% of the atlas. Below, we show that the same biology persists across a broader range of cutoffs.

These cells were sharply concentrated in the late reprogramming window, with both conditions peaking at day 10.5 (Fig. 4j,k). This timing matches the MET window (days 8–12) reported by Schiebinger et al.^3^, when iPSC ancestors pass through the bottleneck between mesenchymal and pluripotent states. Gateway cells also arose predominantly from the MET lineage (Fig. 4l,m): 86% in serum (71 of 83) and 100% in 2i (85 of 85), with small serum contributions from trophoblast (10%, *n* = 8) and epithelial (5%, *n* = 4) populations. Thus, gateway scoring isolates a rare boundary population at the precise temporal and geometric location expected for the reprogramming bottleneck.

#### 2.3.2 A transient gene program defines the gateway state

Could gateway cells simply be arithmetic mixtures of their flanking populations? Under a continuous-drift model, cells would show blended expression, with mesenchymal genes gradually decreasing and pluripotency genes gradually increasing. Under a barrier model, by contrast, crossing from one basin to another should require an additional active program that peaks specifically at the interface. That distinction matters biologically: if gateway cells are only interpolations, then the boundary has no molecular identity of its own; if they carry bell- or valley-shaped genes, then the crossing state is a distinct and potentially mechanistic part of reprogramming.

To distinguish these possibilities, we focused on MET-origin gateway cells and compared them with their two flanking populations, the upstream MET lineage and iPSCs. We classified each gene by its log-fold change (LFC) versus MET and versus iPSC into four categories: **ascending** (MET < gateway < iPSC), **descending** (MET > gateway > iPSC), **bell** (gateway > both), and **valley** (gateway < both). Bell and valley genes mark expression programs that are specific to the boundary state rather than to either endpoint.

Ascending and descending genes provided an internal check that the selected cells lie on the reprogramming trajectory. In both conditions, every gene in these two classes followed strict monotonic ordering (Fig. 5a). The ascending class contained 28 genes in serum and 30 in 2i. In serum, it included the core pluripotency factors *Nanog*, *Sox2*, *Esrrb*, *Klf2*, and *Zfp42* (Rex1), each showing intermediate expression between the low MET baseline and the high iPSC state. In 2i, several of these genes were not significant, consistent with MEK/GSK3*β* inhibition maintaining basal pluripotency-gene expression across the culture^13^. Genes ascending in both conditions included *Tdgf1* (Cripto), *Dppa2*, and *Dppa4*. The descending class (18 genes in serum, 15 in 2i) captured the complementary dismantling of fibroblast identity and extracellular matrix programs. Together, these classes show that gateway cells occupy the expected intermediate position between MET and iPSC states.

**Fig. 5.**
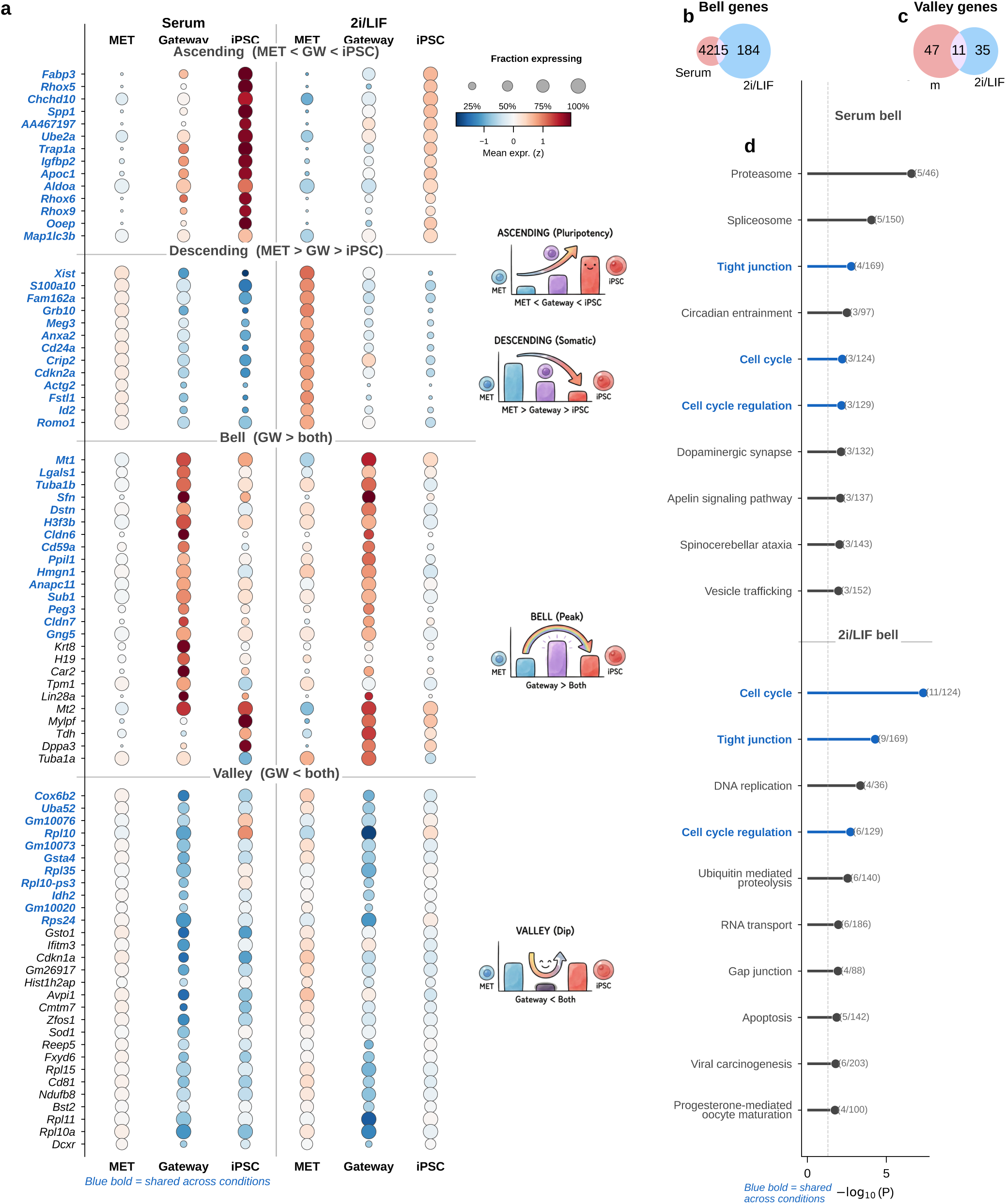
Bell and valley genes define the gateway program across reprogramming conditions. **(a)** Dot plot of top genes in four classes: ascending (MET < GW < iPSC), descending (MET > GW > iPSC), bell (GW > both), and valley (GW < both). Columns show MET, gateway, and iPSC populations for serum (left) and 2i/LIF (right). Dot size encodes the fraction of cells expressing each gene; color encodes mean z-scored log1p expression. Gene names in **blue bold** are shared across conditions. Inset cartoons show the expected expression profile for each class. Serum: 28 ascending, 18 descending, 60 bell, 58 valley. 2i/LIF: 30 ascending, 15 descending, 199 bell, 46 valley. **(b)** Venn diagram of bell genes identified in serum vs. 2i/LIF (overlap of 15 genes; OR = 15.4, *P* < 10^−10^). **(c)** Venn diagram of valley genes (overlap of 11 genes; OR = 57.9, *P* < 10^−12^). **(d)** KEGG pathway enrichment for bell genes in serum (top) and 2i/LIF (bottom), showing the top 10 non-disease terms per condition. Pathways in **blue bold** are shared across both conditions; pathways in black are condition-specific. Overlap gene counts shown in parentheses; dashed line: *P* = 0.05.

The ascending and descending classes mainly serve as descriptive checks that gateway cells sit in the expected intermediate position between MET and iPSC. The stronger test is whether the boundary carries its own transient program. Among genes significant against both reference populations (Methods), we identified 60 bell and 58 valley genes in serum, and 199 bell and 46 valley genes in 2i (Fig. 5a; Extended Data Fig. 3). Of the 60 serum bell genes, 15 were also bell genes in 2i (OR = 15.4, *P* < 10^−10^; Fig. 5b). Valley genes also showed substantial cross-condition conservation (11 shared of 58 in serum and 46 in 2i; OR = 57.9, *P* < 10^−12^; Fig. 5c), with shared genes enriched for ribosomal proteins (*Rpl10*, *Rpl35*, *Rps24*), the ubiquitin–ribosomal fusion gene *Uba52*, and metabolic genes including *Idh2* and the tissue-restricted cytochrome c oxidase subunit *Cox6b2*, consistent with earlier observations that regulated protein synthesis is a recurrent feature of cell-state transitions^16^. These valley patterns are compatible with transient translational reconfiguration or metabolic switching during successful crossing, although some contribution from stress-linked remodeling cannot be excluded from expression data alone. The WOT enrichment, matching analyses, and synthetic-mixture controls argue against interpreting them as simple dropout artifacts. Because the dual-comparison framework has lower statistical power for valley genes than for bell genes in zero-inflated scRNA-seq data, we focused the remaining analysis on bell genes.

The shared bell genes converged on epithelial remodeling. Tight-junction assembly was the top-ranked KEGG pathway in both conditions (Fig. 5d), driven by the claudins *Cldn6* and *Cldn7*, together with the epithelial identity regulator *Sfn* (stratifin/14-3-3*σ*) and the cytoskeletal gene *Tuba1b*, a microtubule component involved in cytoskeletal reorganization during cell-state transitions^17^. This result fits prior work showing that claudins are induced early during reprogramming, before *Nanog*, and that MET is a critical early step in canonical factor-mediated reprogramming, with its inhibition strongly impairing iPSC generation^18–19^.

The shared bell set also included *Mt1* (metallothionein), consistent with a transient stress or redox-related response during reprogramming^20^; the imprinted gene *Peg3*; and chromatin- and transcription-associated genes (*H3f3b*, *Hmgn1*, *Sub1*), suggesting active transcriptional and chromatin reorganization. Importantly, these genes are *bell*, not ascending. They peak at the gateway and then decline again in iPSCs. This transient profile is consistent with a biological requirement for crossing the barrier: the epithelial-assembly machinery, stress-response module, and chromatin remodeling program are maximally engaged at the interface but are not maintained in either stable endpoint. Gateway cells therefore do not behave as simple interpolations of MET and iPSC populations.

The two culture conditions shared this core program but differed in its breadth. The 2i signature was larger (199 bell genes versus 60 in serum; Fig. 5a) and included additional cell-cycle regulators and adhesion molecules. This likely reflects a combination of sharper concentration of successful trajectories under 2i/LIF and reduced off-target heterogeneity, rather than implying that 2i necessarily deploys a completely different gateway biology^21^. The most reproducible recurring module is therefore the 15-gene serum/2i shared core, whereas the condition-specific genes extend that core in different reprogramming contexts.

Gateway cells at non-MET interfaces were rare and limited to the serum condition: Trophoblast→iPSC (*n* = 7) and Epithelial→iPSC (*n* = 15); all 85 gateway cells in 2i were of MET origin (Extended Data Fig. 4). These populations were too small for a well-powered comparison, although the non-overlapping Trophoblast bell set (OR = 0.0) suggests that alternative entry routes may carry distinct transition programs. Overall, the reprogramming gateway is defined by a conserved, transient epithelial-remodeling program that is absent from both flanking populations and peaks at the MET–iPSC boundary.

#### 2.3.3 Independent validation of gateway cell identity

The rarity of the identified gateway cells—83 in serum and 85 in 2i, about 0.05% of cells in either condition—naturally raises the question of whether they represent a genuine biological state or a technical artifact of the scoring procedure. We addressed this through several complementary tests. These included quality-control confounds (Extended Data Fig. 5), orthogonal WOT fate-probability enrichment (Fig. 6a–c; Extended Data Fig. 6), transcriptomic distinctness and cross-condition transfer (Fig. 6d–h; Extended Data Fig. 7), threshold robustness across hard and soft formulations (Extended Data Fig. 8 and Extended Data Fig. 9), matched-control stress tests that ask whether the same cells simply reflect failed or stressed intermediates (Extended Data Fig. 10), a scVI baseline that asks whether BMI creates the state or instead sharpens it (Extended Data Fig. 11), and direct comparison to CellRank and Palantir, which ask whether the same corridor is recovered by methods that target global terminal-outcome ambiguity rather than the local MET–iPSC boundary (Extended Data Fig. 12).

**Fig. 6.**
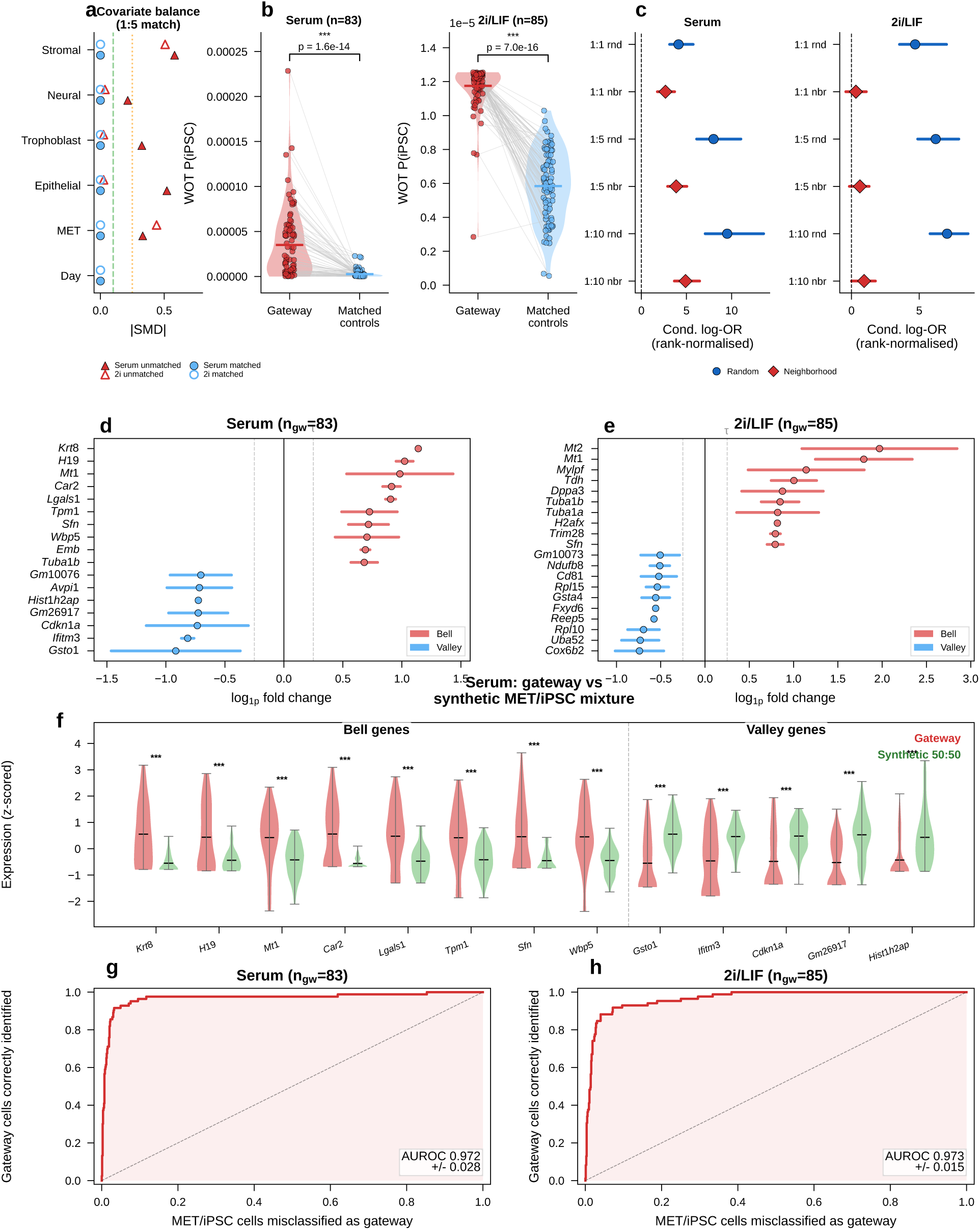
Gateway cells carry a distinct, validated transcriptomic state. **(a)** Love plot of covariate balance before (triangles) and after (circles) 1:5 day+lineage matching for serum and 2i/LIF. All covariates reach |SMD| < 0.1 after matching. **(b)** WOT backward-transport *P*(iPSC) for gateway cells versus their 1:5 day+lineage-matched MET controls (Wilcoxon signed-rank, one-sided; serum *n* = 83, 2i *n* = 85). **(c)** Conditional log-odds ratio across six matching strategies (1:1, 1:5, 1:10; random vs. neighborhood); all significantly positive (bootstrap 95% CI, 2,000 iterations). **(d,e)** Log-fold-change amplitude for the top 10 bell and top 10 valley signature genes (mitochondrial genes excluded) in serum (d) and 2i/LIF (e). Dot: amplitude = (|LFC_*S*_ | + |LFC_*T*_ |)/2; bar spans the two individual LFCs. Dashed line: detection threshold *τ* = 0.25. **(f)** Synthetic mixture control: z-scored expression of top bell and valley genes in real gateway cells (red) vs. synthetic 50:50 MET/iPSC mixtures (green). All tested genes differ significantly (Mann–Whitney *U*, *P* < 0.05; serum 17/17, 2i 20/20). **(g,h)** ROC curves for a 20-gene classifier (top 10 bell + top 10 valley; 5-fold CV with majority-class undersampling) in serum (g; AUROC = 0.968) and 2i/LIF (h; 0.974).

**Fig. 7.**
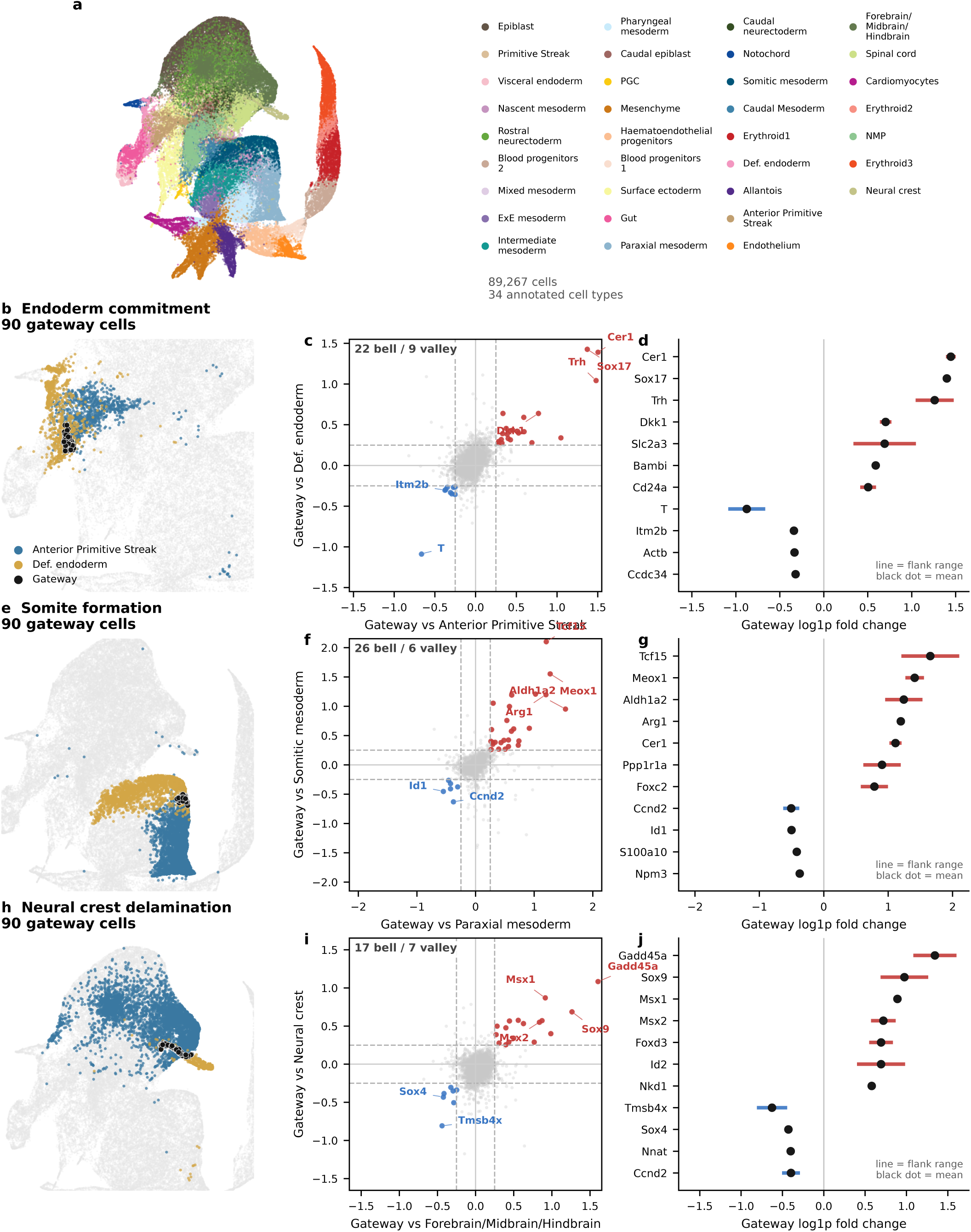
Gateway cells at gastrulation fate boundaries. **(a)** UMAP of the full gastrulation atlas (Pĳuan-Sala et al.; E6.5–E8.5, 89,267 cells, 34 cell types), colored by cell type. **(b,e,h)** Per-transition UMAP projections. Flanking cell types in red and blue; gateway cells (black circles, *n* = 90 per boundary) localize at each fate boundary. **(c,f,i)** Four-quadrant scatter plots. Each gene is plotted by LFC versus fate A (*x*) and fate B (*y*). Bell genes (red, upper-right) are elevated at the gateway relative to both flanks. Valley genes (blue, lower-left) are depressed at the gateway. **(d,g,j)** LFC amplitude forest plots. Bell genes (red bars, positive) are listed above valley genes (blue bars, negative). Black dots: mean of the two flanking LFCs; horizontal lines: range. Columns: endoderm commitment (22 bell, 9 valley; **b,c,d**), somite formation (26 bell, 6 valley; **e,f,g**), neural crest delamination (17 bell, 7 valley; **h,i,j**).

**Fig. 8.**
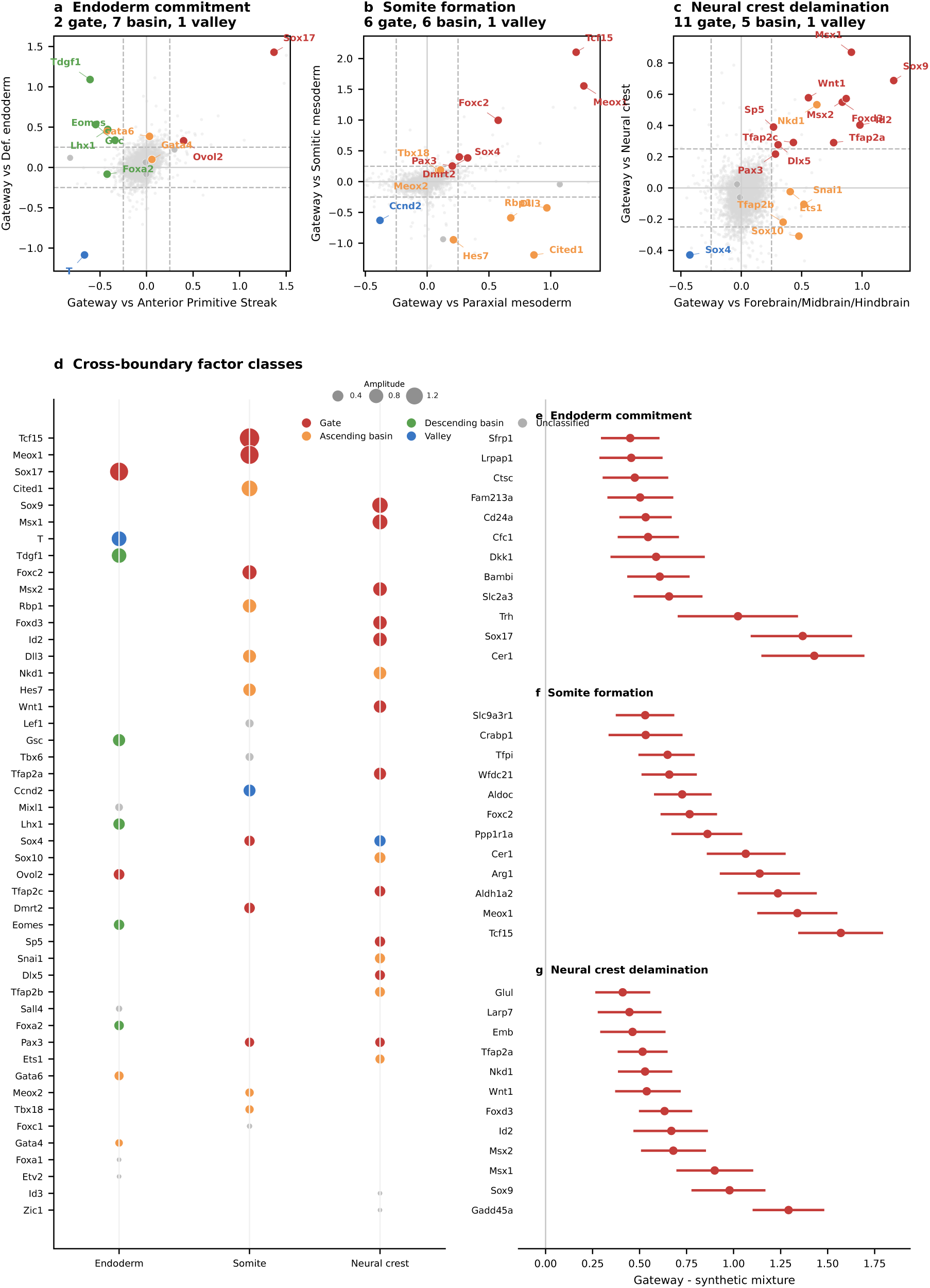
Known gastrulation regulators separate into gate-position and basin-position classes. **(a–c)** Four-quadrant scatter plots showing literature-curated regulators among all genes at each boundary. Colors indicate gate-position factors (red), ascending basin-position factors (orange), descending basin-position factors (green), valley factors (blue), or unclassified factors (gray). **(a)** Endoderm commitment: 2 gate-position (*Sox17*^31^, *Ovol2*), 7 basin-position, and 1 valley (*T*). **(b)** Somite formation: 6 gate-position (*Tcf15*^33^, *Meox1*^34^, *Foxc2*^35^, *Dmrt2*^36^, *Sox4*, *Pax3*^45^), 6 basin-position, and 1 valley. **(c)** Neural crest: 11 gate-position (*Sox9*^39^, *Msx1/2*^41^, *Foxd3*^40^, *Tfap2a/c*^38^, *Wnt1*^42^, *Dlx5*^43^, *Sp5*^44^, *Pax3*^45^, *Id2*^46^), 5 basin-position, and 1 valley (*Sox4*). **(d)** Cross-boundary dot plot. Each row: one curated regulator present in at least one focal boundary context. Dot color: classification. Dot size: LFC amplitude. Gray crosses: factor not detected or not classified at that boundary. *Sox4* switches from gate-position (somite) to valley (neural crest), whereas *Pax3* remains gate-position across the somite and neural-crest boundaries. **(e–g)** Synthetic mixture controls. Mean expression difference (gateway cells − 50:50 mixture of flanking populations) with 95% bootstrap CIs (*n* = 2,000 resamples) for the top 12 bell genes at each boundary. All CIs exclude zero, confirming that gateway cells are distinct transcriptional states rather than simple blends of the flanking populations.

**Fig. 9.**
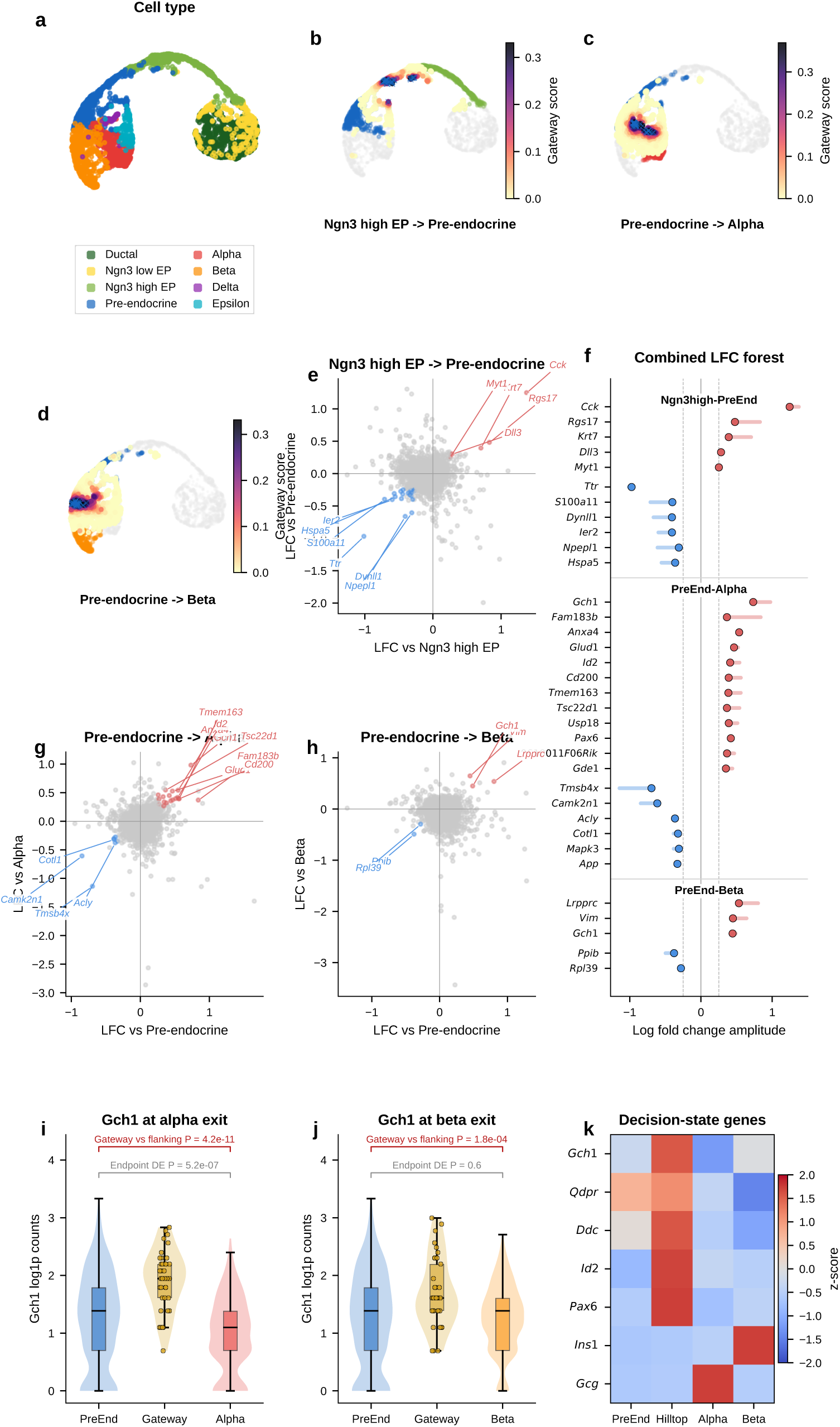
Gateway analysis reveals hub entry and a candidate shared alpha/beta pre-commitment interval. Gateway analysis of 3,696 E15.5 mouse pancreas cells from Bastidas-Ponce et al. 2019^4^. These cells span the endocrinogenesis trajectory. **(a)** UMAP of the BMI-regularized latent space colored by cell type. The trajectory runs from Ductal cells through Ngn3-expressing endocrine progenitors (Ngn3 low EP, Ngn3 high EP) to Pre-endocrine cells and four terminal hormone-producing fates (Alpha, Beta, Delta, Epsilon). **(b)** Gateway score overlay for the Ngn3 high EP–Pre-endocrine boundary (37 gateway cells at the 99th percentile, blue crosses). **(c)** Gateway score overlay for the Pre-endocrine–Alpha boundary (37 gateway cells). **(d)** Gateway score overlay for the Pre-endocrine–Beta boundary (37 gateway cells). **(e)** Four-quadrant LFC scatter for Ngn3 high EP–Pre-endocrine: 5 bell genes and 19 valley genes. The valley-dominated pattern indicates that entry into the pre-endocrine hub is driven mainly by coordinated silencing, with a narrow bell-gene burst led by *Cck*. **(f)** Combined LFC amplitude forest plot for all three boundaries (Ngn3 high EP–Pre-endocrine: all 5 bell + 6 of 19 valley; Pre-endocrine–Alpha: top 12 of 19 bell + 6 of 7 valley; Pre-endocrine–Beta: all 3 bell + 2 valley). Shared bell genes are shown in bold, highlighting recurrence of *Gch1* across the alpha and beta exits. **(g)** Four-quadrant LFC scatter for Pre-endocrine–Alpha: 19 bell genes and 7 valley genes. The alpha-exit gateway carries the richest commitment program and is led by *Gch1* and *Pax6*. **(h)** Four-quadrant scatter for Pre-endocrine–Beta: 3 bell genes and 2 valley genes. *Gch1* recurs as a bell gene, showing that the candidate BH4-associated hilltop program is shared across both endocrine exits. **(i,j)** *Gch1* expression (log_1*p*_ counts) in Pre-endocrine, gateway, and endpoint cells at the Alpha (i) and Beta (j) boundaries. At the Alpha boundary, endpoint DE classifies *Gch1* as decreased (*P* = 5.2 × 10^−7^), not increased. At the Beta boundary, endpoint DE detects no change (*P* = 0.60). Gateway-versus-flanking tests recover the transient peak at both boundaries (*P* = 4.2 × 10^−11^ for Alpha; *P* = 1.8 × 10^−4^ for Beta), showing that the shared candidate BH4-associated pre-commitment signal is missed by conventional endpoint-based DE. **(k)** Shared hilltop genes: z-scored mean expression across Pre-endocrine, Hilltop, Alpha, and Beta populations. The BH4-associated genes *Gch1* and *Qdpr*, together with *Ddc*, peak in the shared transition-state compartment, whereas lineage hormones (*Ins1*, *Gcg*) peak only after commitment.

**Fig. 10.**
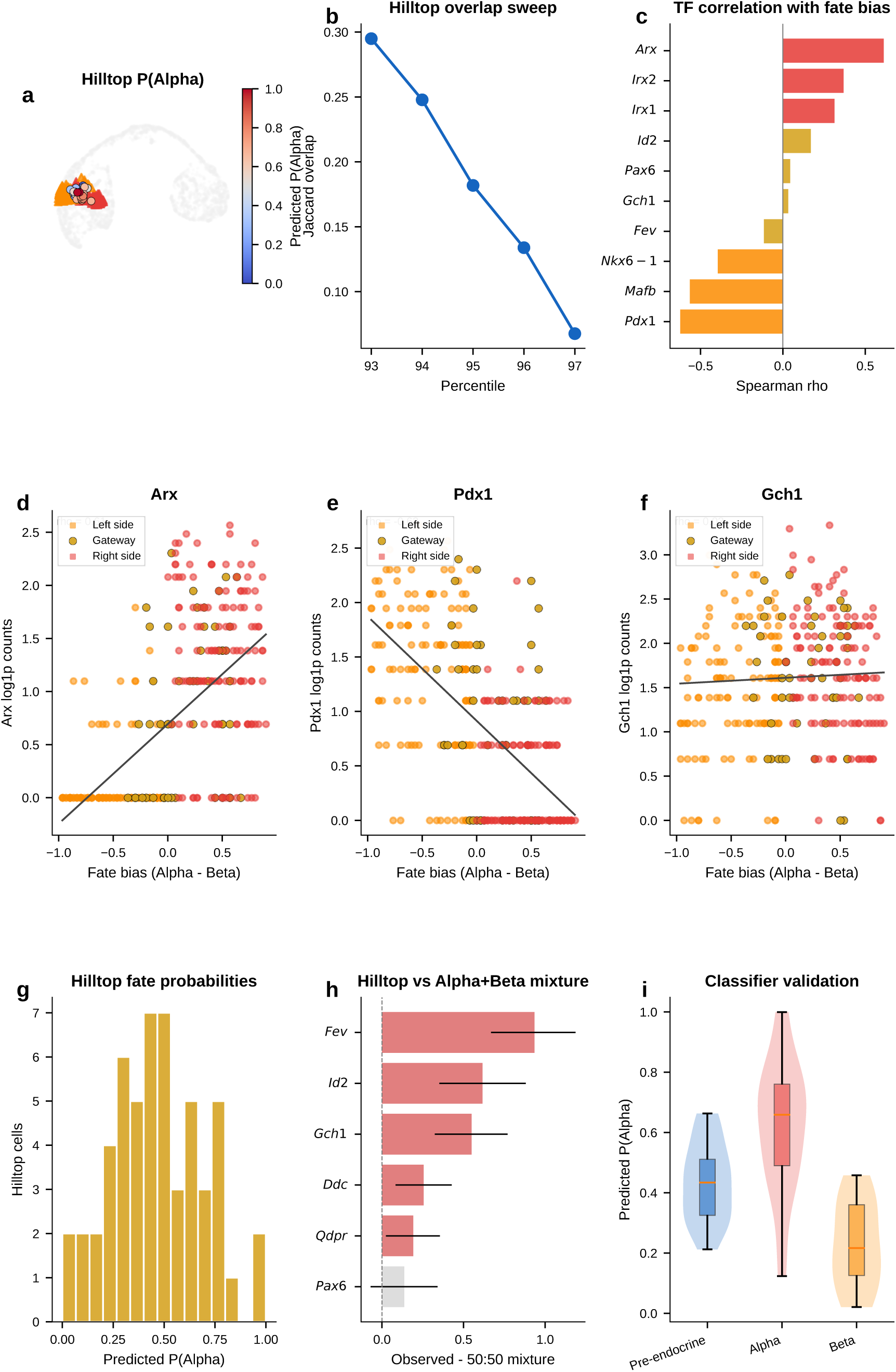
Gateway overlap reveals a shared endocrine hilltop and separates shared-interval genes from lineage-bias gradients. **(a)** Hilltop cells (*n* = 54, 95th-percentile overlap of the alpha-exit and beta-exit gateways) colored by classifier-predicted *P*(Alpha). Red/orange triangles mark alpha-exit and beta-exit gateway cells, respectively. **(b)** Jaccard overlap between the two gateway populations across percentile thresholds. Overlap emerges progressively as the threshold is relaxed, revealing the shared precursor zone that direct Alpha–Beta comparison does not isolate. **(c)** Spearman correlation of key regulators with fate bias (pull toward Alpha minus pull toward Beta). Alpha determinants (*Arx*, *Irx2*, *Irx1*) are positively correlated; beta-skewed regulators (*Pdx1*, *Nkx6-1*) are negatively correlated; *Mafb* also trends toward that side in the current E15.5 analysis; *Gch1* and *Pax6* remain close to the neutral zone, whereas *Id2* is mildly alpha-biased and *Fev* is weakly beta-leaning in the current analysis. **(d–f)** Expression of *Arx* (d), *Pdx1* (e), and *Gch1* (f) versus fate bias for all cells near the hilltop. Left-flank cells, hilltop cells, and right-flank cells are colored separately, and a fitted trend line is overlaid. *Arx* and *Pdx1* form opposing gradients, whereas *Gch1* remains largely uncorrelated. Spearman *ρ* is shown. **(g)** Predicted *P*(Alpha) for hilltop cells. A classifier trained only on exit-cell labels with no fate-bias input reaches 82% mean cross-validated accuracy and assigns most hilltop cells toward one fate while retaining an intermediate band. **(h)** Mixture control: hilltop mean minus synthetic 50:50 Alpha+Beta mixture mean for selected shared hilltop genes (2,000 bootstraps; 95% CIs). *Gch1*, *Qdpr*, *Ddc*, *Id2*, and *Fev* are enriched above the mixture expectation, arguing against simple averaging as an explanation for hilltop expression. **(i)** Classifier validation restricted to hilltop cells and stratified by actual cell-type label. Alpha-labeled hilltop cells receive higher predicted *P*(Alpha) (mean 0.62) than beta-labeled hilltop cells (mean 0.23), whereas Pre-endocrine hilltop cells remain intermediate (mean 0.44), consistent with graded lineage bias within the shared pre-commitment interval.

**Fig. 11.**
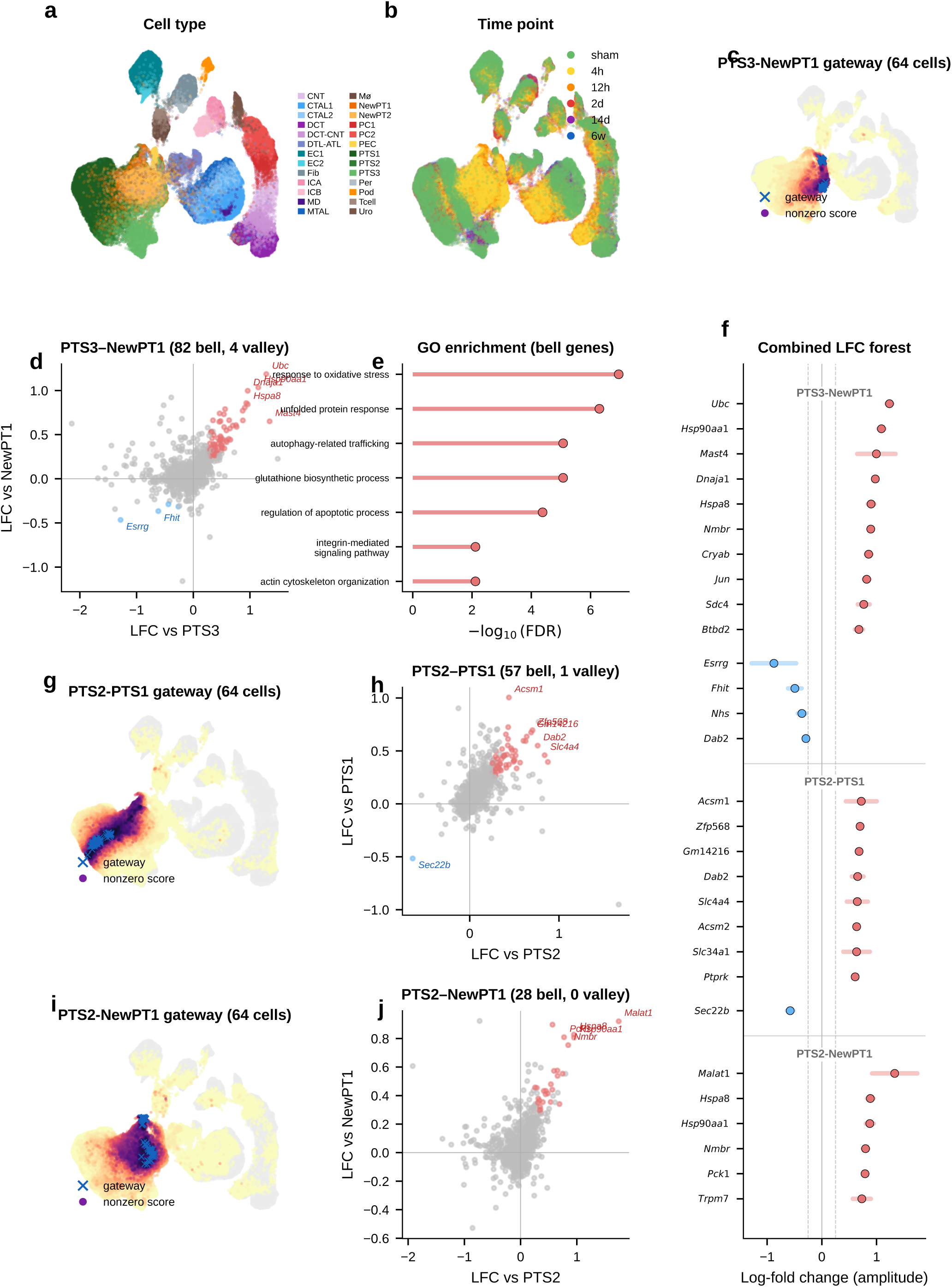
Gateway cells at injury-related and neighboring proximal-tubule boundaries in mouse kidney. Gateway analysis of 126,578 nuclei from mouse kidney after ischemia–reperfusion injury (IRI; 26 cell types; 6 time points; Kirita et al. 2020^11^). **(a)** UMAP colored by cell type (26 types). **(b)** UMAP colored by time point (sham, 4 h, 12 h, 2 d, 14 d, 6 w). **(c)** Gateway score overlay for the PTS3–NewPT1 boundary (64 gateway cells at the 99.95th percentile, blue crosses; color scale: magma_r). Gateway cells localize at the interface between homeostatic (PTS3) and injury-activated (NewPT1) proximal-tubule states. **(d)** Four-quadrant LFC scatter for PTS3–NewPT1: 82 bell genes (red) and 4 valley genes (blue). Top bell genes include established stress/remodeling genes such as *Ubc*, *Hsp90aa1*, *Dnaja1*, *Hspa8*, and *Jun*, together with additional candidates such as *Nmbr*. **(e)** GO enrichment of PTS3–NewPT1 bell genes (top 8 terms). The dominant terms highlight unfolded-protein response, protein ubiquitination, apoptotic regulation, autophagy-related trafficking, and glutathione biosynthesis. **(f)** Combined LFC amplitude forest plot across the three main kidney boundaries (PTS3–NewPT1: top 10 bell + 4 valley; PTS2–PTS1: top 8 bell + 1 valley; PTS2–NewPT1: top 6 bell). **(g)** Gateway score overlay for the PTS2–PTS1 boundary (64 gateway cells). **(h)** Four-quadrant LFC scatter for PTS2–PTS1: 57 bell genes and 1 valley gene define a transport-focused maturation program at the healthy PT boundary. **(i)** Gateway score overlay for the PTS2–NewPT1 boundary (64 gateway cells). **(j)** Four-quadrant LFC scatter for PTS2–NewPT1: 28 bell genes at the transition from healthy to injured PT, led by *Malat1* and *Pck1*.

Technical controls did not support an artifact explanation. Gateway cells showed similar quality metrics to non-gateway cells (Extended Data Fig. 5a–h): sequencing depth was unchanged in serum and only modestly higher in 2i, ribosomal-gene fraction was unchanged in serum, and mitochondrial fraction was modestly higher in both conditions. Scrublet^22^ predicted-doublet rates were very low (1.2% in serum and 0.0% in 2i; Extended Data Fig. 5i,j), arguing against doublets as the source of the signal. Gateway cells were enriched in G1 phase, consistent with the known link between mitochondrial regulation and reprogramming^23^.

The clearest orthogonal benchmark in this dataset is WOT itself. Waddington-OT estimates each cell’s probability of reaching iPSC by solving a temporal optimal-transport problem across densely sampled time points. The gateway score, by contrast, knows nothing about time; it asks only whether a cell sits in a neighborhood connected to both fates in the learned latent space. Despite these different inputs, both approaches converged on the same rare population (Fig. 4e–h). Gateway cells had higher *P*(iPSC) than surrounding cells, and this advantage persisted after matched-control analysis in which each gateway cell was paired with five non-gateway MET controls from the same day and source lineage (Fig. 6a). The enrichment remained significant after matching (*P* < 10^−6^, Wilcoxon signed-rank; Fig. 6b), was consistent across six matching strategies (Fig. 6c), and was detectable at each tested time point (Extended Data Fig. 6). Matched-control comparisons also argue against a simple stress-only or failed-transition explanation. Relative to 1:5 latent-neighborhood matched controls, gateway cells show only modest generic stress and are not enriched for apoptosis/senescence signatures. They also remain distinct from weighted synthetic MET:iPSC mixtures across the tested mixing ratios and alternative lineage controls. The overlap between serum and 2i bell genes is well above permutation null expectations (Extended Data Fig. 10; the curated stress and apoptosis/senescence signatures used for these controls are listed in Methods).

The expression effect sizes were also large and reproducible (Fig. 6d,e). Using the average absolute log-fold change against both flanking populations as an amplitude measure (Methods), the top bell genes included *Krt8*, *H19*, *Mt1*, and *Sfn* in serum, and *Mt1*, *Mt2*, and *Krt8* in 2i. To test whether this profile could arise from a simple arithmetic mixture of MET and iPSC cells, we generated synthetic 50:50 MET+iPSC profiles (Fig. 6f). All tested non-mitochondrial signature genes differed significantly between real gateway cells and the synthetic mixture (Mann–Whitney *U*, *P* < 0.05; 17/17 in serum and 20/20 in 2i). Bell genes were expressed higher in real gateway cells than in the mixture, and valley genes lower, showing that gateway cells overshoot and undershoot the midpoint rather than averaging it.

If gateway cells define a coherent transcriptional state, expression alone should identify them. A logistic-regression classifier trained on 20 genes (top 10 bell + top 10 valley) achieved AUROC = 0.97 in both serum and 2i by 5-fold cross-validation (Fig. 6g,h), separating ∼80 gateway cells from tens of thousands of flanking MET and iPSC cells. In cross-condition transfer, the strongest performance came from a conserved 10-gene shared tier (7 bell + 3 valley), which reached AUROC = 0.76 for serum→2i and 0.74 for 2i→serum while retaining 94–95% of in-domain performance (Extended Data Fig. 7). Adding source-specific genes improved within-condition fits but reduced transfer. Shared-tier model coefficients remained positively correlated between conditions (*r* = 0.69, *P* = 2.8 × 10^−2^; 8 of 10 shared genes with the same sign), indicating a conserved core program.

Threshold sweeps reached the same conclusion. Across stringent cutoffs, WOT enrichment rose sharply and median bell-gene amplitudes remained broadly stable, while bell-gene yield varied with stringency, especially in 2i/LIF. Shared bell overlap persisted across this high-stringency regime, with *Sfn* retained throughout and *Cldn6*/*Cldn7* remaining shared through the operating range (Extended Data Fig. 8).

Any hard threshold raises a basic question: if we avoid the binary cutoff, can we still detect bell and valley genes that distinguish the gateway from its flanking populations? If not, the gateway cells might just be a threshold artifact. To check this, we repeated the analysis with a complementary soft-threshold scheme (Extended Data Fig. 9). We defined a restricted positive-score soft core in each condition. Within that soft core, higher-scoring cells contributed more to the gateway-side expression mean and lower-scoring cells less. We then compared this weighted gateway profile with the flanking source and target populations.

The result confirms the existence of gateway populations with distinct gene programs even without binary membership. In serum, the soft-core recovered 55 bell and 61 valley genes. Forty bell genes were shared with the hard-threshold list, with 60% overlap at both the top 10 and top 20, and shared amplitudes remained correlated (Spearman *ρ* = 0.61). In 2i/LIF, the soft-core recovered 133 bell and 22 valley genes. It shared 104 bell genes with the hard-threshold list, with 50% overlap at the top 10, 55% at the top 20, and amplitude correlation *ρ* = 0.51. The shared serum core remained led by *Krt8*, *H19*, *Mt1*, *Car2*, and *Sfn*.

Thus, even without relying on the hard cutoff, the analysis still detects gateway populations with distinct gene programs in both conditions. The main differences lie in the broader low-score halo, especially in 2i/LIF. We therefore retain the manual hard-threshold definition for the rest of the paper because it provides a discrete operating set of gateway cells for gene calling, figure assembly, and cross-dataset comparisons.

A final question is whether gateway cells depend on the BMI-regularized latent space or instead mark a genuine transition corridor that other methods can also detect. The answer is the latter, with BMI contributing resolution rather than inventing the biology.

Plain scVI still highlighted a broad MET–iPSC transition region, indicating that the underlying corridor is also visible without BMI regularization. Extended Data Fig. 11a–d shows the same broad corridor on the gateway-scored embeddings themselves. However, without BMI regularization the local gateway population was absorbed into a much broader positive-score zone. This loss of local resolution was reflected directly in bell-gene yield. In serum, scVI recovered only 8 hard bell genes versus 60 with scAttnVI; in 2i/LIF, it recovered 57 versus 199. Using the full positive-score transition zone as a score-weighted soft comparison, scVI recovered 2 versus 51 bell genes in serum and 1 versus 65 in 2i/LIF (Extended Data Fig. 11e,f). This full-zone weighted analysis is distinct from the restricted soft-core analysis in Extended Data Fig. 9. Thus, BMI regularization does not create the transition state de novo. It preserves the rare local geometry needed to separate a compact gateway population and recover the genes that peak there.

CellRank^9^ and Palantir^24^ reach a related conclusion from a different direction. These methods identify high-entropy cells with unresolved terminal outcomes. In WOT, both recover the broader transition corridor surrounding the MET–iPSC bottleneck. This confirms that gateway cells lie within a bona fide region of fate uncertainty. Extended Data Fig. 12 shows both the shared serum corridor and the stronger divergence in 2i/LIF. Gateway analysis, however, targets a narrower object: the rare cells that sit at the local interface between annotated flanking states. This added local resolution is clearest in 2i/LIF. There, CellRank and Palantir broaden toward a larger MET-versus-stromal uncertainty field, whereas gateway analysis retains a compact MET–iPSC bottleneck and the richer bell-gene program associated with it. Gateway analysis also shows substantially greater cross-condition coherence than the entropy-based selections. Local pairwise interface detection therefore yields a more stable transient program than global branch entropy alone (Extended Data Fig. 12).

Together, these analyses indicate that the WOT benchmark contains a compact barrier-crossing population with a reproducible bell/valley gene program at the MET–iPSC interface. Gateway analysis therefore detects a real transition population rather than a scoring artifact or simple average of the flanking states.

### 2.4 Gateway analysis of gastrulation supports two classes of fate-boundary regulators

We next turned to mouse gastrulation—the developmental process that first inspired Waddington’s epigenetic landscape^1^ and the setting in which fate-boundary regulators have been studied extensively^25–28^. Decades of genetics have identified key regulators at major germ-layer boundaries. What they do not resolve is a finer question: does a TF act at the boundary itself, or does it accumulate only after commitment? Gene regulatory network theory predicts both classes: transient regulators that help cells cross the boundary and downstream regulators that stabilize the committed fate^28,52–54^. Because gateway analysis assigns genes to the boundary or the basin, gastrulation provides the clearest test in this study of whether known developmental regulators separate into boundary-enriched and post-commitment classes.

We applied the framework to the Pĳuan-Sala et al.^10^ atlas (E6.5–E8.5; 89,267 cells, 34 cell types) and scored six boundaries spanning all three germ layers. Gateway analysis identified cells across these developmental interfaces, and we focus here on the three transitions with the richest gateway programs: endoderm commitment (Anterior Primitive Streak to Definitive Endoderm; 90 gateway cells, 22 bell and 9 valley genes), somite formation (Paraxial Mesoderm to Somitic Mesoderm; 90 gateway cells, 26 bell and 6 valley genes), and neural crest specification (Forebrain/Midbrain/Hindbrain to Neural Crest; 90 gateway cells, 17 bell and 7 valley genes). We use the literature-defined landscape geometries as biological context; the method itself asks where the boundary cells and boundary programs lie. The purpose of the gastrulation analysis is to test gate-versus-basin regulator logic across boundaries of mixed geometry. Before drawing that biological distinction, however, the method first has to show that it can isolate bona fide developmental interfaces.

Gateway cells localized sharply to each developmental interface. To rule out technical artifacts, we compared gateway and non-gateway cells across total UMI counts, mitochondrial read fraction, cell-cycle phase composition, ribosomal gene fraction, and doublet-density proxy scores. Library size, mitochondrial fraction, cell-cycle phase composition, and ribosomal fraction were all similar between gateway and non-gateway cells, whereas the local doublet proxy was *lower*, not higher, in gateway cells. Synthetic mixture controls, discussed below, further confirmed that gateway cells are not arithmetic averages of the flanking populations.

The three boundaries also carried distinct molecular programs. The endoderm gate was dominated by secreted Wnt/BMP/Nodal antagonists, consistent with a signaling-attenuation gate. The somitic gate combined TF activation with retinoic-acid biosynthesis, marked by *Aldh1a2*^29^. The neural crest gate recovered core neural plate border and early neural crest regulators from the neural crest gene regulatory network^26–27^. These distinct programs argue against a generic gastrulation gateway and instead point to boundary-specific gate architecture. Together, these analyses make the technical point first: gateway analysis identifies real, non-generic developmental interfaces with boundary-specific programs.

With that interface structure in place, we next asked a more focused question: where do known lineage regulators sit relative to the gate and the basin? We compiled 49 curated gastrulation regulators with strong developmental support across the three focal transitions—14 for endoderm^31–32^, 16 for somitogenesis^25,33–37^, and 19 for neural crest^26–27,38^—drawing primarily on knockout, lineage-tracing, and canonical GRN studies. Each factor was classified as a **gate-position regulator** if it is a bell gene (elevated at the gateway relative to both flanking fates) or a **basin-position regulator** if it is ascending or descending (high in one flanking population but not at the gate). The 49 curated factors split into 19 gate-position (bell), 18 basin-position (ascending or descending), 3 valley, and 9 unclassified factors. Regulators long known to control gastrulation do not all act at the same position in the landscape. That two-class separation is the main gastrulation result: known regulators partition into factors that peak at the boundary itself and factors that accumulate only after commitment.

The clearest example is the neural crest boundary. Of 19 known NC-GRN regulators, 11 are gate-position regulators, spanning neural plate border and early neural crest specifiers^26–27^: *Sox9*, *Foxd3*, *Msx1/2*, *Tfap2a/c*, *Wnt1*, *Dlx5*, *Sp5*, *Pax3*, and *Id2*^38–46^. Later-acting regulators *Sox10*, *Snai1*, *Ets1*, *Tfap2b*, and *Nkd1* are basin-position regulators, consistent with stronger expression after the initial specification peak rather than at the boundary crest itself^26–27,47^. Gateway analysis is consistent with the known NC-GRN hierarchy while placing the classified factors at either the boundary or the flanking basin.

The endoderm and somitic boundaries reveal two additional boundary implementations of the same gate-versus-basin logic. At the endoderm boundary, only *Sox17*^31^ and *Ovol2* are gate-position regulators, while *Eomes* and *Lhx1*, both enriched on the APS side, decline toward the endoderm basin and *Foxa2* and *Gata6* rise toward the endoderm basin^48–50^. The gate is instead characterized by the same secreted Wnt/BMP/Nodal antagonists noted above, making this interface a signaling-attenuation gate rather than a TF-dominated one. At the somitic boundary, the gate combines TF activation (*Tcf15*, *Meox1*, *Foxc2*, *Dmrt2*, *Sox4*, *Pax3*) with retinoic-acid biosynthesis marked by *Aldh1a2*, consistent with the known role of RA signaling in somitogenesis^29–30,33–36,45^. The Notch-clock components *Hes7* and *Dll3*, required for maintaining segmentation periodicity^25,37^, are basin-position regulators. We therefore interpret gate-position and basin-position as boundary-localized expression classes in this atlas, not as universal labels for regulator function across every developmental context.

The gate–basin placement is not a fixed property of a protein but depends on the boundary. *Msx1*^41^ is a gate-position regulator at the neural crest boundary but a valley gene at the Intermediate Mesoderm–Somitic boundary; *Sox4* switches from gate-position (somite) to valley (neural crest); *Pax3* is gate-position at both boundaries, consistent with its appearance in both the somitic and neural-crest regulator lists; and *Ccnd2* ^51^ appears as a valley gene at the somitic boundary, consistent with cell-cycle exit as a shared prerequisite for commitment. This context-dependence is itself consistent with GRN theory^52–54^. These classifications rest on genuine expression differences at the boundary: in the current analysis, each top bell gene was expressed more strongly in real gateway cells than in synthetic 50:50 blends of the two flanking populations.

The gastrulation result is also robust to moderate threshold relaxation. Across the three focal boundaries, relaxing the operating cutoff preserved much of the core bell program, whereas stronger relaxation mainly expanded the gateway pool and added lower-amplitude genes. Only two curated regulators, *Tfap2a* and *Pax3* at the neural-crest boundary, lie close to the bell-gene decision boundary. Thus, the central gate-position versus basin-position conclusion does not depend on a large set of borderline regulator classifications.

Taken together, the gastrulation analysis is consistent with the idea that fate-boundary GRNs may use at least two recurrent regulatory architectures^28^. Some regulators peak at the gate and are consistent with switch-associated roles, whereas others accumulate in the basin and are consistent with stabilization after commitment. More broadly, the two gate logics suggest different experimental strategies: signaling-attenuation gates such as the endoderm interface may be more accessible to soluble ligands or small-molecule pathway modulation, whereas TF-cascade gates such as the somitic and neural crest boundaries may require pathway-level activation or genetic perturbation to redirect fate.

### 2.5 Gateway analysis resolves a shared pre-commitment hilltop in pancreatic endocrinogenesis

During pancreatic endocrinogenesis, multipotent pre-endocrine progenitors pass through a brief hub state and then exit toward four hormone-producing fates—alpha, beta, delta, and epsilon. Developmental genetics and single-cell atlases have mapped this topology and many mature lineage regulators^4^. What has remained difficult to see directly is the short interval in which endocrine competence is still shared. Those cells are rare and transient.

We applied gateway analysis to E15.5 mouse pancreas^4^ (3,696 cells, 8 types), where pre-endocrine cells sit at the center of this fate hub with exits toward each endocrine lineage. We identified compact gateway cores at each boundary. Three features emerged. First, hub entry is marked by a sharp transient *Cck* peak against a broader background of coordinated silencing. Second, the alpha and beta exits point back to a shared pre-commitment hilltop where candidate BH4-associated genes peak before lineage divergence. Third, alpha-versus-beta bias then emerges as opposing TF gradients away from that shared interval.

#### Entry into the pre-endocrine hub is marked by coordinated silencing and a transient Cck-enriched peak

We first asked how the endocrine hub is entered. The upstream Ngn3 high EP to Pre-endocrine boundary captures the transition from Ngn3-expressing progenitor to multipotent pre-endocrine cell. This is the most valley-dominated boundary in the study. Hub entry is marked mainly by coordinated shutdown rather than by broad new gene activation. Chaperones (*Hspa5*, *Hspe1*), progenitor markers (*Vim*, *S100a11*), and the transthyretin *Ttr* are all transiently silenced, consistent with proteome remodeling at cell-fate transitions^16^. Against that repressive backdrop, one gene stands out. *Cck* (cholecystokinin; amplitude 1.31) is the highest-amplitude bell gene at any pancreatic boundary. *Cck* expression in endocrine progenitors has been reported previously^4^. Gateway analysis localizes that Cck-enriched peak to the narrow interval between progenitor specification and pre-endocrine multipotency.

Because endocrine commitment is coupled to cell-cycle withdrawal, we next asked whether this entry gateway simply captured cells exiting the cycle. It did not. The Ngn3 high EP/Pre-endocrine corridor was already predominantly G1, and entry gateway and hilltop cells are not phase outliers within the endocrine trajectory. The leading entry bell genes also remain gateway-enriched after adjustment for continuous S and G2/M scores and after comparison only to other G1 cells from the same corridor. Cell-cycle exit is therefore part of the biological context of hub entry, but it does not explain away the localized gateway state.

#### Gateway analysis reveals a candidate shared alpha/beta pre-commitment interval that endpoint differential expression misses

We next asked whether the alpha and beta exits contain a shared transient program rather than only lineage-specific endpoint markers. The alpha exit carries the richest gateway program and recovers known and plausible commitment-associated genes including *Pax6*, *Glud1*, *Id2*, and *Hspa8* without prior marker input. The strongest signal is the top-ranked bell gene *Gch1*, which encodes the rate-limiting enzyme for tetrahydrobiopterin (BH4) biosynthesis. BH4 is a cofactor for aromatic amino-acid hydroxylases and nitric-oxide synthases. *Gch1* peaks at the gateway, higher than in both progenitors and committed alpha cells. A standard endpoint comparison would classify it as *decreased* rather than increased and therefore miss this transient peak. At the parallel beta exit, *Gch1* recurs together with *Lrpprc*. Endpoint differential expression again misses the transition-state signal. The key point is simple: this is not an alpha-specific endpoint feature. Its recurrence at both exits is consistent with a shared pre-commitment program that appears before alpha and beta cells fully diverge.

#### The alpha and beta exit gateways overlap at a shared hilltop within the pre-endocrine hub

If that shared program is real, the next question is where it sits. The recurrence of *Gch1* at both exits suggested that the two gateway populations might share cells upstream of commitment. At the hard gateway cutoff, the alpha-exit and beta-exit gateway sets are disjoint. As the cutoff is relaxed, overlap emerges continuously. We therefore defined a compact shared overlap core of 54 cells at the 95th-percentile overlap threshold. Under a matched-size null within the combined Pre-endocrine, Alpha, and Beta compartment, the null yields 18.5 cells on average. The observed 54-cell hilltop is a 2.9-fold enrichment (*P* = 1.9 × 10^−15^, exact hypergeometric). This supports a concentrated shared pre-commitment interval rather than a broad visual overlap on the same manifold.

#### Opposing TF gradients emerge across the hilltop, whereas the BH4-associated genes remain largely fate-neutral

Using a continuous fate-bias score based on kNN pull toward Alpha versus Beta neighbors, the analysis found clear opposing gradients of lineage determinants. *Arx*, *Irx2*, and *Irx1* increase toward the alpha exit, whereas *Pdx1* and *Nkx6-1* increase toward the beta exit; *Mafb* also trends toward that side in the current E15.5 analysis. By contrast, *Gch1* is uncorrelated with fate bias, and *Pax6* and *Fev* lie near the neutral zone. The shared hilltop program therefore only partially overlaps with the genes that bias lineage choice. The lineage-biased TFs define the graded axis toward alpha or beta identity, whereas the BH4-associated genes mark a candidate shared transition program that is present before that choice is resolved.

#### Independent controls support a candidate shared hilltop program

Two orthogonal controls support this interpretation. First, a logistic-regression classifier trained only on exit cells using all expressed genes and no fate-bias input provides supportive evidence for graded lineage bias. The exit-only classifier reaches 82% mean cross-validated accuracy; alpha-labeled hilltop cells receive higher predicted *P*(Alpha) than beta-labeled hilltop cells, while pre-endocrine hilltop cells remain intermediate. Second, the BH4-associated genes *Gch1* and *Qdpr*, together with the monoamine-synthesis enzyme *Ddc*, are enriched at the hilltop relative to both pre-endocrine cells alone and a synthetic 50:50 Alpha plus Beta mixture. This is inconsistent with the idea that hilltop enrichment is explained simply by immaturity or by averaging the two fates.

#### A staged model for endocrine commitment

Taken together, the pancreas analysis provides molecular detail consistent with the staged model of endocrine commitment^4^ rather than a smooth one-step drift into mature hormone identity. Entry into the hub is marked by coordinated silencing with a transient *Cck* peak. Within the hub, the alpha and beta exit gateways share a compact hilltop population where *Gch1* and *Qdpr*, together with *Ddc*, are transiently elevated. Lineage choice then emerges through opposing alpha-versus-beta TF gradients.

#### Temporal replication at E14.5

An analysis of 5,385 E14.5 endocrine-lineage cells from the same study^4^ recovers the same overall topology and retains *Cck* as the top-ranked entry bell gene (amplitude 1.26 versus 1.31 at E15.5; Extended Data Fig. 21). The candidate BH4-associated program is also retained, although less sharply than at E15.5: at the alpha exit, *Gch1* remains a strong bell gene but ranks second behind *Entpd3* (amplitude 0.663 versus 0.854 at E15.5), and at the beta exit it recurs again as a weaker bell gene (rank 7; amplitude 0.498 versus 0.465 at E15.5). The ranking is therefore not identical across timepoints, but the broader conclusion is preserved: the endocrine hub still contains a shared candidate BH4-associated pre-commitment signal before alpha and beta fates fully diverge. Scavuzzo et al.^63^ detected *Gch1* expression in Ngn3^+^ endocrine progenitors by fluorescence in situ hybridization, supporting a progenitor-stage interpretation of this BH4-associated signal.

### 2.6 A proteostasis-centered program marks the kidney injury boundary

We next tested gateway analysis in a disease-related setting. Acute kidney injury provides a natural transition between homeostatic and injury-activated proximal-tubule states. Homeostatic proximal-tubule cells dedifferentiate, proliferate, and either redifferentiate into functional epithelium or stall in a pro-fibrotic state associated with chronic kidney disease^11^. The first step out of the healthy state is therefore a candidate bottleneck. The cells occupying that interval are rare and intermediate between established annotations.

We applied the BMI framework to a mouse ischemia–reperfusion injury (IRI) dataset^11^ comprising 126,578 nuclei across 26 cell types and 6 time points—two orders of magnitude larger than the pancreas dataset. As an initial sanity check, we asked whether the method would recover *Havcr1* (KIM-1), the FDA-qualified urinary biomarker that is undetectable in healthy kidney but rapidly induced upon injury. Scoring the interface between homeostatic (PTS3) and injury-activated (NewPT1) proximal tubule cells, the framework correctly recovered *Havcr1* without any prior marker input, supporting the biological plausibility of the selected injury-entry boundary.

#### An 82-gene proteostasis-centered injury-entry program peaks at the transition

Gateway analysis uncovered an extensive transition program at the PTS3–NewPT1 boundary: 82 bell and 4 valley genes—the largest gateway program in the current analysis. The top bell genes include established proteostasis and stress-response factors *Ubc*, *Hsp90aa1*, *Dnaja1*, *Hspa8*, and *Jun*, together with less-characterized candidates such as *Nmbr* and *Mast4*. Taken together, they define a proteostasis-enriched module with additional stress and remodeling factors. These genes are compatible with an acute stress response, but they peak specifically at the gateway and are lower in the established injury state than at the boundary itself. If this were merely a diffuse injury signature, the signal should be strongest in fully injured cells. Instead, the proteostasis response is enriched at the transcriptional boundary between healthy and injured states.

Several genes in this program overlap with broadly stress-responsive genes that can also appear in technical artifact settings. However, the Kirita et al. atlas was generated by single-nucleus RNA-seq from frozen tissue^11^, so the signal is less likely to reflect enzymatic dissociation alone. The stronger point, however, is its spatial restriction: a purely technical stress signal would be expected to appear more broadly across boundaries, whereas this program concentrates at the PTS3–NewPT1 transition. That pattern is more consistent with a genuine injury-entry program.

This proteostasis burst is coupled to broader injury remodeling. The immediate-early transcription factor *Jun* indicates AP-1 pathway activation. Glutathione-synthesis enzymes *Gclm* and *Gclc* mount a coordinated antioxidant defense consistent with the glutathione/Nrf2 arm of AKI responses. *Myo5b* points to apical-trafficking and brush-border remodeling at the same boundary. Gene Ontology reinforces the same picture: unfolded-protein response, protein ubiquitination, apoptotic regulation, autophagy-related trafficking, and glutathione biosynthesis all score prominently. Gateway analysis resolves an injury-entry-associated module in which proteostasis, redox defense, and remodeling programs are co-enriched at the transition boundary. A synthetic-mixture control (Extended Data Fig. 22d) argues that this is an active state and not a simple blending artifact.

#### Boundary-specific programs argue against a purely generic stress artifact

A likely alternative explanation is that these 82 genes simply mark stressed or dying cells. A cross-boundary comparison argues against a purely generic version of that interpretation. At the homeostatic boundary between healthy S2 and S1 segments (PTS2–PTS1), the 57 bell genes are dominated by transport markers and lack the proteostasis signature. At the transition from healthy S2 to injured PT (PTS2–NewPT1), the 28-gene program is led instead by the lncRNA *Malat1* and the gluconeogenic enzyme *Pck1*, indicating a distinct metabolic transition. Only 6 of the 167 boundary–gene pairs are shared pairwise, and no gene appears at all three boundaries, indicating that the 82-gene program has strong atlas-specific distinctiveness at the PTS3–NewPT1 entry bottleneck.

To ask whether the PTS3–NewPT1 gateway simply reflects early injured PTS3 cells, we matched each IRI gateway cell to five non-gateway PTS3 controls from the same sample and time point whenever possible. Under this stricter control, *Havcr1* itself was no longer distinctive, indicating that its recovery should be treated as expected behavior rather than as a standalone validation. By contrast, *Ubc*, *Hsp90aa1*, *Hsp90ab1*, *Dnaja1*, *Hspa8*, *Jun*, *Gclm*, and *Gclc* remained elevated relative to time-matched PTS3 controls, whereas classical apoptosis markers such as *Casp3*, *Bax*, and *Bcl2* remained weak. This pattern argues against canonical caspase-dependent apoptosis as the main explanation for the gateway signal, although it does not exclude other injury-associated death programs. These controls sharpen the interpretation: the gateway marks a temporally localized injury-entry interval within early IRI PTS3 rather than simple endpoint NewPT1 identity or a gross dying-cell compartment.

#### Narrowing the transition interval

The advance over prior injury atlases is not just a longer list of injury genes. It is resolution of the commitment interval itself. By isolating the window in which proteostasis, antioxidant defense, and remodeling genes are maximally co-expressed, this gateway signature provides a candidate molecular profile of the injury-entry interval. More broadly, the resulting injury-entry signature may help distinguish cells actively entering the injury program from cells already in the injured state.

## 3 Discussion

Single-cell atlases excel at naming stable cell states, but the intervals between those states are still difficult to resolve. The cells that occupy those intervals are rare, short-lived, and easily absorbed by much larger flanking populations during embedding and endpoint-based differential expression^5–6^. Gateway analysis was developed to recover those pairwise intervals directly: which cells occupy a boundary, and which genes peak specifically within it. Across four systems spanning reprogramming, embryonic development, endocrine differentiation, and injury, the same framework recovered compact, boundary-specific populations with distinct bell/valley gene programs. In WOT, those cells align with the part of the trajectory carrying the highest iPSC fate probability and lie within the broader transition corridor identified by CellRank and Palantir. In gastrulation, pancreas, and kidney, matched controls, synthetic mixtures, threshold-sensitivity analyses, and cross-boundary comparisons support the same interpretation. The main contribution is therefore local resolution: gateway analysis isolates the compact interface itself and the genes that distinguish it, rather than averaging both into neighboring basins.

### Fate boundaries contain active transient programs

Across all four systems, the boundary is not an empty gap between endpoints. It contains a short-lived transcriptional program that rises locally and then subsides. In reprogramming, that program is marked by an epithelial remodeling module (*Cldn6*, *Cldn7*, *Sfn*) at the late MET-to-iPSC interface. In gastrulation, curated regulators separate into boundary-enriched and basin-enriched classes across boundaries of mixed geometry. In pancreas, gateway analysis resolves a transient *Cck* peak at hub entry and a shared candidate BH4-associated pre-commitment hilltop before alpha-versus-beta bias becomes strong. In kidney, it resolves a proteostasis-centered injury-entry interval that is distinct from both homeostatic PT states and the NewPT1 endpoint. These signals are not generic transition noise and are not well summarized by endpoint markers alone. Kidney injury-entry, maturation, and metabolic gateway programs are largely distinct, and the injury-entry set overlaps only minimally with endpoint NewPT1 markers. In pancreas and gastrulation, synthetic-mixture controls and context-dependent regulator behavior support the same conclusion. Pairwise boundary analysis therefore compresses broad transition responses into shorter, more interpretable bell/valley gene sets, such as BH4-associated endocrine-entry genes in pancreas and injury-entry markers in kidney. The same analysis consistently reveals real boundary populations and the molecular programs that mark them.

### Pattern-based similarity makes rare boundaries computationally visible

BMI succeeds because it prioritizes coordinated on/off gene patterns instead of raw expression magnitude. That choice makes rare interface cells remain similar to one another even when they sit between much larger mature populations. When BMI is used as a structural prior in scAttnVI (Equation 4), the graph acts like a set of local springs that preserves compact transition neighborhoods in latent space. Gateway detection then operates on neighborhoods shaped by shared program structure rather than by cell-count dominance. Because the required inputs are already available for many published atlases, this framework can also be used to re-read existing datasets as maps of decision intervals rather than only catalogs of endpoints.

### Limitations

Gateway cells are intrinsically rare, typically 0.05–1% of a dataset, so sensitivity remains limited by sampling depth. The framework recovers these populations across datasets ranging from ∼3,700 to ∼127,000 cells, and larger atlases should expose additional boundaries and rarer transition intervals. Increasing temporal resolution—for example through denser time-course sampling or metabolic pulse-chase labeling—should further improve capture of these fleeting states. The gateway score is also agnostic to direction of transit; here, directionality was resolved by integrating WOT backward transport and developmental timing, and future combination with RNA velocity^8^ or CellRank^9^ should extend the framework to datasets lacking lineage information. In addition, the current implementation relies on pre-existing cell-type annotations to define flanking populations, so it is strongest when those flanking states are already biologically credible and the goal is to refine the interval between them. Extending the method to unsupervised or annotation-free boundary discovery would broaden applicability. Finally, bell and valley genes are transcriptomic associations that nominate candidates for functional validation, and valley calls remain less powered than bell calls because of the expression floor and zero-inflation (Supplementary Note 4). The candidates reported here should therefore be viewed as focused entry points for perturbation rather than as complete inventories of every molecule operating at a boundary.

### Outlook

Gateway analysis opens an experimentally tractable window onto a class of biology that is already present in existing single-cell data but has remained difficult to resolve. By pinpointing where a decision occurs and which genes rise or fall only within that interval, the framework turns vague transition zones into perturbable states. Extending this approach to multimodal measurements, spatial context, perturbation screens, and cross-species atlases will test how general these organizing principles are. Across reprogramming, gastrulation, pancreas, and kidney, the data support a common picture: Waddington’s boundaries can contain active transient states with distinct molecular programs, not merely empty gaps between attractors. Mapping those states provides a candidate entry point for testing how transition-local programs may influence cell-fate progression in regenerative medicine and for studying how repair-associated trajectories diverge in disease.

## 4 Methods

### Resource availability

### Data and code availability

- All single-cell RNA-seq datasets analyzed in this study are publicly available from their original publications and source repositories. The benchmark and case-study atlases used here are cited in the main reference list and described under **Datasets** below.
- A public gateway-analysis code repository is available on GitHub as hucang0/gateway_scattnvi. The current public release is a notebook-first WOT gateway demo centered on `GatwaWay_scAttnVI.ipynb` and companion Python helpers for downloading the public WOT tutorial bundle, preparing per-condition inputs, training scAttnVI embeddings, calling MET →iPSC gateway cells, and plotting bell and valley genes.
- The scAttnVI/BMI training path used for full-scale reruns was executed on CUDA-capable GPUs and should be treated as a GPU-backed workflow for full-scale analysis. Small smoke tests and plotting utilities can be run without the full training footprint, but the main model-training path is not intended as a CPU fallback.

### Experimental model and study participant details

This study is entirely computational and does not involve new experimental models, animal experiments, or human participants. All datasets were obtained from previously published studies.

### Method details

#### Datasets

Five publicly available single-cell RNA-seq datasets were used in this study, comprising one benchmark dataset and four biological case studies:

1. **PBMC (peripheral blood mononuclear cells).** The PBMC dataset^12^ was obtained from the 10X Genomics public datasets repository. It contains ∼12,000 cells profiled by Chromium Single Cell 3^′^ chemistry with well-characterized immune populations including CD8 T cells, CD4 T cells, B cells, NK cells, and monocytes.
2. **WOT (Waddington–OT reprogramming).** Schiebinger et al.^3^ profiled mouse embryonic fibroblasts undergoing reprogramming to induced pluripotent stem cells (iPSCs) at half-day intervals across days 0–18 under serum and 2i/LIF conditions. The rebuild used 165,892 serum cells and 168,631 2i/LIF cells. The trajectory training sets excluded spike-in anchor iPSC cells, while the corresponding full matrices were retained for full-gene BMI and downstream validation summaries.
3. **Gastrulation.** Pĳuan-Sala et al.^10^ profiled 89,267 cells spanning 34 annotated cell types from mouse embryos at E6.5–E8.5.
4. **Pancreas (endocrinogenesis).** Bastidas-Ponce et al.^4^ profiled 3,696 E15.5 mouse pancreatic cells across ductal, endocrine-progenitor, and terminal endocrine populations. An E14.5 replication cohort of 5,385 endocrine-lineage cells from the same study was processed separately for temporal validation.
5. **Kidney (acute kidney injury).** Kirita et al.^11^ profiled 126,578 nuclei from mouse kidneys after ischemia–reperfusion injury across sham, 4 h, 12 h, 2 d, 14 d, and 6 w time points.

#### Preprocessing

Raw UMI count matrices were used as input throughout. Standard quality-control filters followed the source studies when available. Highly variable genes (HVGs) were selected with `scanpy.pp.highly_variable_genes`^55^ using the `seurat_v3` flavor and `span = 0.3:` 4,000 HVGs for gastrulation and kidney, 2,000 for pancreas and the E14.5 pancreas replication, and the 1,479 HVGs distributed with the WOT consortium release. Batch-aware HVG selection (`batch_key = sample`) was used for gastrulation and kidney. No log1p normalization was applied before BMI computation or scAttnVI/scVI training; models were trained directly from the integer counts layer.

#### BMI computation

Binary mutual information (BMI) between cell pairs was computed as described in §2.1. Gene expression was binarized at a count threshold *τ* = 1 (detected/not detected). Genes with detection frequency below 1% or above 75% were excluded before pairwise BMI calculation. For each cell pair (*i*, *j*), the joint detected/not-detected frequencies across the retained genes were used to compute BMI according to Equation 3. BMI was computed on the full gene set rather than on the HVG subset. A null distribution was estimated from 50 column permutations of the binarized matrix, edges below the 25th percentile of the null were zeroed, and each cell retained its top *k* = 32 BMI neighbors to form a sparse graph.

#### scAttnVI and scVI model architecture and training

The scAttnVI model extends scVI^6^ by adding a BMI-derived latent-distance penalty. Unless noted otherwise, all models used *n*_latent_ = 10, learning rate 10^−3^, random seed 42, and a maximum of 400 epochs with early stopping. For the large atlases (WOT, gastrulation, kidney), the reruns used *n*_hidden_ = 256, *n*_layers_ = 4, batch size = 4,096, and early-stopping patience = 30. For pancreas and the E14.5 pancreas replication, the reruns used *n*_hidden_ = 128, *n*_layers_ = 2, batch size = 512, and early-stopping patience = 45. For the PBMC penalty sweep, a smaller benchmark model was used (*n*_hidden_ = 128, *n*_layers_ = 1, batch size = 512, 300 epochs).

BMI-derived coefficients served as fixed attraction weights in latent space. We retain the name scAttnVI for continuity, but these weights are static BMI-derived attention-like coefficients rather than learned Transformer-style attention. The shared softmax temperature was *T* = 0.25 and the mixing coefficient was *⍺* = 1.0. The penalty weight *λ*_BMI_ was selected per dataset by dataset-specific penalty sweeps, with the benchmarking and operating-point rationale described in Supplementary Note 1.

The scAttnVI reruns were executed on GPU and intentionally did not fall back to CPU. For the WOT baseline comparison, plain scVI was retrained on the same localized WOT inputs and then passed through the same downstream gateway pipeline on the resulting X_scvi latent space.

UMAP embeddings were computed from the learned latent space for visualization only. The default setting used scanpy.tl.umap^55^ with *n*_neighbors_ = 30 and min_dist = 0.3. The kidney analysis used a scanpy/UMAP configuration with *n*_neighbors_ = 15. All quantitative claims rest on latent-space neighborhoods, differential expression, matched comparisons, synthetic-mixture controls, and orthogonal validations rather than on the visual embedding itself.

#### Gateway score computation

Gateway scoring followed §2.3.1. For each cell, a fate pull toward each annotated state was computed as the fraction of its latent-space nearest neighbors annotated as that state, and the gateway score was defined as the minimum of the pulls toward the two flanking populations. Personalized PageRank^14–15^ diffusion was then applied on a separate inverse-distance-weighted latent-space graph with *k* = 30, *⍺* = 0.5, and three iterations.

The fate-pull neighborhood size used in the analyses was *k* = 30 for WOT, pancreas, and kidney, and *k* = 50 for the finalized gastrulation main figure and its matching robustness analyses. Gateway cells were defined by the diffused-score percentile threshold used for each dataset: 99.95th percentile for WOT and kidney, 99.9th for gastrulation, and 99^th^ for pancreas. For the pancreas hilltop analysis, overlap between the Pre-endocrine→Alpha and Pre-endocrine→Beta gateway sets was evaluated across percentile thresholds, with hilltop cells defined as the 95th-percentile overlap within the endocrine corridor.

These thresholds were chosen per dataset to balance sensitivity against contamination by nearby non-boundary cells. Smaller atlases, such as pancreas, required a looser cutoff to retain enough gateway cells for stable gene calling, whereas larger atlases, such as WOT and kidney, supported more stringent thresholds while still retaining compact gateway sets.

#### Bell and valley gene classification

Genes with fewer than 5% of cells expressing them in the combined gateway plus reference set, or with log1p-normalized variance below 0.01, were excluded from testing. Gateway cells were compared with both flanking populations using two-sided Mann–Whitney *U* tests, and *P* values were corrected by the Benjamini–Hochberg procedure. Genes were classified as ascending, descending, bell, or valley according to the sign of the two flanking log-fold changes and an amplitude threshold *τ* = 0.25 (log1p scale). For the general classification workflow, amplitude was defined as *A* = (|LFC_*S*_ | + |LFC_*T*_ |)/2.

#### Soft-threshold and hilltop analyses

For the WOT soft-threshold analysis, we defined a soft core as the top 5% of cells within the positive diffused-score zone in each condition and retained continuous gateway-score weights within that restricted zone. Weighted differential expression was then compared with the hard-threshold gateway result. This is an alternative soft-threshold analysis, not a continuous relaxation over all positive-score cells.

For the pancreas hilltop analysis, hilltop overlap significance was evaluated against matched-size null draws restricted to the endocrine corridor using both exact hypergeometric calculations and 100,000 matched-size random draws at each threshold.

#### Stratified matching and WOT validation

To control for temporal and lineage composition in WOT, each gateway cell was matched to controls from the same experimental day and source-lineage stratum; when exact day-plus-lineage matches were insufficient, controls were drawn from the same day regardless of lineage. Multiple matching strategies were explored, and the 1:5 matched-control strategy is reported in the main text. WOT backward-transport *P*(iPSC) values were compared with one-sided Wilcoxon signed-rank tests and summarized with conditional logistic regression on rank-normalized values.

For the kidney time-matched control analysis, each IRI gateway cell at the PTS3→NewPT1 boundary was paired with five non-gateway PTS3 controls from the same IRI sample and time point whenever possible; if sample-and-timepoint matches were insufficient, controls were drawn from the same IRI time point.

#### Synthetic mixture controls

To test whether gateway cells represented distinct states rather than computational averages of their flanking populations, we constructed synthetic 50:50 mixtures by randomly pairing one cell from each flank and averaging their log1p-normalized expression profiles. The synthetic set was matched in number to the observed gateway population and assessed with 2,000 bootstrap resamples. Genes whose 95% bootstrap confidence intervals for (real gateway − synthetic mixture) excluded zero were considered distinct from the synthetic mixture.

#### Logistic regression classifiers

For WOT, a balanced logistic-regression classifier was trained on the top 10 bell plus top 10 valley genes to distinguish gateway cells from their flanking MET and iPSC populations. Non-gateway cells were subsampled to reduce class imbalance, and performance was evaluated by 5-fold stratified cross-validation. Cross-condition transfer was assessed by training in one culture condition and testing in the other without retraining.

For the pancreas hilltop analysis, a separate logistic-regression classifier was trained only on alpha-exit and beta-exit cells using all expressed genes and no fate-bias input. Performance was evaluated by 5-fold cross-validation, and predicted P(Alpha) values were projected onto hilltop, alpha, beta, and Pre-endocrine populations.

#### Comparison with scVI, CellRank, and Palantir in WOT

For the WOT scVI baseline, plain scVI was retrained on the same localized serum and 2i/LIF counts/HVG inputs used for scAttnVI, and the same gateway-scoring, diffusion, and bell/valley classification pipeline was then rerun on `X_scvi`.

For the CellRank/Palantir comparison, each condition was restricted to the MET, epithelial, iPSC, and stromal reprogramming states plus a capped set of unassigned early cells, then normalized, log-transformed, and projected by PCA. CellRank^9^ used a PseudotimeKernel + ConnectivityKernel GPCCA estimator with manually selected root and terminal medoid cells. Palantir used PCA, diffusion maps, multiscale space construction, and the same root/terminal logic. For each method, the highest-entropy cells were selected with the same cell count as the hard gateway set in the corresponding condition before comparing overlap, bell-gene recovery, and WOT fate enrichment.

#### Regulator classification in gastrulation

For gastrulation, 49 curated regulators with established or strongly supported developmental roles at the three focal boundaries were compiled from the literature. Each factor was classified as gate-position if it met bell criteria, basin-position if it was ascending or descending, valley if it met valley criteria, or unclassified otherwise. Context dependence was assessed by comparing classifications across boundaries.

#### Technical confound analysis

Gateway cells were compared with non-gateway cells on library size, number of detected genes, mitochondrial fraction, ribosomal fraction, and cell-cycle state. Where source-study S/G2M scores were provided (for example, the pancreas atlases), those annotations were used directly; otherwise, cell-cycle states were assigned from mouse orthologues of the Tirosh et al.^61^ signatures. Doublet analysis used Scrublet^22^ or the source study’s precomputed doublet-density proxy when the corresponding analysis relied on that published annotation. Continuous confound comparisons used Mann–Whitney *U* tests and cell-cycle proportions were compared with *χ*^2^ tests.

#### Curated stress and apoptosis/senescence gene sets

For matched-control module analyses, stress- and apoptosis-related signatures were fixed before testing rather than optimized on the target datasets. The WOT matched-control comparison used a compact epithelial gateway signature, a generic stress signature, and an apoptosis/senescence signature. The kidney time-matched control analysis used separate stress/proteostasis and apoptosis/senescence modules. The exact gene sets were:

| Context | Signature | Genes |
| --- | --- | --- |
| WOT | MET-gate | *Cldn6, Cldn7, Sfn, Krt8, Krt18, Epcam* |
| WOT | Generic stress | *Mt1, Mt2, Hspa8, Hsp90aa1, Hsp90ab1, Dnaja1* |
| WOT | Apoptosis/senescence | *Peg3, Cdkn1a, Bax, Bbc3, Trp53inp1, Gadd45a, Igfbp7* |
| Kidney | Stress/proteostasis | *Ubc, Ubb, Hsp90aa1, Hsp90ab1, Dnaja1, Hspa8, Jun, Gclm, Gclc* |
| Kidney | Apoptosis/senescence | *Casp3, Bax, Bcl2, Bbc3, Cdkn1a, Trp53inp1, Gadd45a* |

Representative literature support for these curated choices was grouped at the gene or gene-pair level: *Cldn6*, *Cldn7*, *Krt8*, *Krt18*, *Epcam*, and *Sfn* as epithelial/MET-associated genes during reprogramming^18–19^; *Mt1*/*Mt2* as stress-inducible metallothioneins^64^; *Hspa8*/*Dnaja1* and *Hsp90aa1*/*Hsp90ab1* as chaperone-based stress-response genes^65–66^; *Peg3*^67^, *Cdkn1a*^68^, *Bax*/*Bcl2*^69^, *Bbc3* ^70^, *Trp53inp1*^71^, *Gadd45a*^72^, and *Igfbp7* ^73^ as apoptosis/senescence-linked genes; *Ubc*/*Ubb*^74^, *Jun* ^75^, *Gclm*^76^, *Gclc*^77^, and *Casp3* ^78^ as kidney stress/redox or executioner-apoptosis genes.

In the processed WOT feature space, *Krt18*, *Hspa8*, *Hsp90aa1*, and *Hsp90ab1* were not retained in the final gene matrix, so the matched-control module scores were computed on the present subset of each signature. By contrast, all genes listed for the kidney stress and apoptosis/senescence modules were present in the final kidney matrix and were scored as specified above.

#### Pathway and Gene Ontology enrichment

Enrichment analyses were performed on bell-gene sets using all expressed genes as background. For WOT, pathway-style summaries were derived from the local rerun and reported as the top non-disease pathway enrichments. For gastrulation, pancreas, and kidney, GO/pathway enrichment summaries followed a g:Profiler-style workflow^58^ (GO:BP, GO:MF, KEGG, Reactome; FDR < 0.05), and the top reported terms were rendered for each focal boundary.

#### Quantification and statistical analysis

All statistical tests and their parameters are reported in the text and figure legends where they appear. In brief:

- **Differential expression**: two-sided Mann–Whitney *U* tests with Benjamini–Hochberg FDR correction (*⍺* = 0.05) and minimum amplitude threshold > 0.25.
- **Matched comparisons**: one-sided Wilcoxon signed-rank tests on matched pairs, with conditional logistic regression and bootstrap summaries where reported.
- **PBMC thinning benchmark**: repeated retraining across five random seeds with paired benchmark summaries.
- **Bootstrap confidence intervals**: 2,000 resamples for synthetic-mixture controls.
- **Cross-validation**: 5-fold stratified cross-validation for classifier performance.
- **Confound comparisons**: Mann–Whitney *U* tests for continuous metrics and *χ*^2^ tests for categorical phase distributions.

False-discovery correction was applied within each boundary analysis. Cross-boundary comparisons are descriptive unless a separate cross-boundary test is reported explicitly.

All analyses were performed in Python using `scanpy, scvi-tools, scipy, statsmodels, scikit-learn, matplotlib, CellRank`, and `Palantir`. Training and analysis runs for the reruns used CUDA-capable GPU execution for full-scale scAttnVI/BMI retraining. The public `gateway_scattnvi` repository uses a lightweight conda environment with `scvi-tools, torch`, and related analysis dependencies, but full-scale scAttnVI/BMI retraining still assumes CUDA-capable GPU execution.

## Acknowledgements

This work was supported by NIH R01 grant 5R01HG014004.

## 5 Supplementary Notes

### Supplementary Note 1: *λ*_BMI_ selection

The BMI penalty weight *λ*_BMI_ controls the trade-off between reconstruction fidelity and coexpression-based regularization in the scAttnVI loss (Equation 4). At *λ*_BMI_ = 0 the model reduces to standard scVI; as *λ*_BMI_ increases, the latent space is increasingly shaped by BMI-derived neighborhood structure rather than reconstruction alone. We therefore choose *λ*_BMI_ by balancing data fit against regularization.

We select *λ*_BMI_ for each dataset using *L-curve analysis*^56–57^. The idea is simple: plot the data-fit term (here, reconstruction loss) against the penalty term (normalized BMI penalty) as *λ* varies. On a log–log scale this curve is typically L-shaped: for small *λ*, the penalty term decreases rapidly with little reconstruction cost (the vertical arm), whereas for large *λ* the reconstruction loss grows steeply while the penalty term barely improves (the horizontal arm). The corner, or *elbow*, marks the regime where additional regularization begins to cost more in reconstruction fidelity than it returns in penalty reduction.

We trained scAttnVI on the PBMC dataset (11,990 cells) across 16 values of *λ*_BMI_: 0 (scVI baseline), 100, 250, 500, 750, 1,000, 1,500, 2,000, 3,000, 4,000, 6,000, 8,000, 12,000, 16,000, 32,000, and 64,000. Each configuration was trained with three independent random seeds (300 epochs, batch size 512, *n*_latent_ = 10, *n*_hidden_ = 128, 1 layer). For each model we recorded the reconstruction loss (ELBO reconstruction component) and the normalized BMI penalty, and evaluated two downstream metrics at full CD8 retention (1,448 cells) and 5% thinning (72 cells): CD8 *k*NN retention (*k* = 30, fraction of nearest neighbors that are true CD8 cells) and CD8 cluster purity (best Leiden cluster, *k* = 12 neighbors, resolution 1.0).

The L-curve (Extended Data Fig. 1a) exhibits the characteristic L-shape described above^56^. At *λ*_BMI_ = 4,000 the reconstruction loss increases by 4.5% relative to scVI (*λ* = 0), while the BMI penalty has dropped to 0.0035—well into the diminishing-returns regime where further increases in *λ* yield little additional penalty reduction. The dual-axis elbow plot (Extended Data Fig. 1b) confirms that *λ* = 4,000 sits within the elbow range (*λ* ∈ [2,000, 8,000], shaded green), where the BMI prior is effective without substantially degrading reconstruction quality. This is the expected L-curve behavior: values below the elbow leave the BMI term underweighted, whereas values above it trade reconstruction fidelity for only small additional regularization gains^57^.

At full CD8 retention, *λ* = 4,000 improves CD8 *k*NN retention by +0.045 relative to scVI (from 0.870 to 0.916; Extended Data Fig. 1c). Under extreme thinning (5% retention, 72 CD8 cells), the improvement is +0.135 (from 0.291 to 0.426), confirming that BMI regularization specifically benefits rare populations. CD8 cluster purity follows a similar pattern: at 5% thinning, *λ* = 4,000 improves purity by +0.172 relative to the scVI baseline (Extended Data Fig. 1d).

Both metrics remain stable across the elbow range (*λ* ∈ [2,000, 8,000]): at full retention, *k*NN retention varies by < 0.013 and cluster purity by < 0.06 across these four *λ* values. Under thinning, *k*NN retention varies by < 0.10 and purity by < 0.08. The biological conclusions of the PBMC analysis (§2.2) are therefore not sensitive to the specific choice of *λ*_BMI_ within this range.

We recommend selecting *λ*_BMI_ at the L-curve elbow for each new dataset, using the reconstruction loss and BMI penalty as the two axes. The elbow identifies the regime where the prior is strong enough to anchor rare-population geometry without collapsing the latent space into a BMI-dominated representation. For the PBMC benchmark, values in the range [2,000, 8,000] all produce qualitatively similar embeddings and downstream results. We chose *λ*_BMI_ = 4,000 as the representative value for the analyses in §2.2. Gateway scoring is then performed on latent-space neighborhoods after training, so the chosen *λ*_BMI_ affects the learned geometry on which the score is evaluated.

### Supplementary Note 2: Gateway score diffusion

The raw gateway score can contain isolated high-scoring cells generated by dropout noise or stochastic gene detection. To suppress such outliers, we optionally apply personalized PageRank (PPR) diffusion^14–15^ on a *k*-nearest-neighbor graph (*k* = 30) in the scAttnVI latent space (Equation 7; *⍺* = 0.5, *T* = 3 iterations). At each iteration, a cell’s score is replaced by a convex combination of its own raw score (weight 1 − *⍺*) and the weighted average of its neighbors’ scores (weight *⍺*). Isolated spikes are therefore attenuated, whereas spatially coherent boundary signal is reinforced. Diffusion is not essential for the analyses reported here; the main-text conclusions remain the same with raw scores.

Extended Data Fig. 2 compares raw and diffused results. Diffusion changes the bell-gene count, but a shared cross-condition bell core remains preserved. WOT concordance is also retained: the odds ratio for enrichment of gateway cells in the top decile of WOT *P*(iPSC) remains high in both conditions. In UMAP space, most gateway cells are shared between the raw and diffused sets, and cells unique to one set lie in the immediate neighborhood of the shared cells along the MET–iPSC boundary (Extended Data Fig. 2d,e). Across the tested sweep, serum remains nearly flat whereas 2i/LIF becomes more permissive at stronger diffusion; the chosen setting (*⍺* = 0.5, *T* = 3) lies on the lower-*⍺* plateau before that high-diffusion expansion (Extended Data Fig. 2a,b).

### Supplementary Note 3: Gateway score threshold selection

The gateway score *S*_gateway_ is a continuous quantity: every cell receives a value reflecting how strongly it sits at the interface between the iPSC lineage and the dominant non-iPSC fate. Converting this continuous score into a discrete set of gateway cells requires choosing a threshold. Higher thresholds retain fewer cells but sharpen the contrast between the gateway population and its flanking states; lower thresholds include more cells but dilute the boundary signal. We screened a range of thresholds to find the operating point that best balances statistical power with biological specificity, and to confirm that the core findings are robust across this range (Extended Data Fig. 8).

The raw gateway score is extremely sparse: only 0.6% (serum) and 2.2% (2i) of cells receive a non-zero value. After PageRank diffusion with *⍺* = 0.5 and depth = 3, the score becomes much denser, making the upper tail of the score distribution the relevant operating regime for defining gateway cells. The local rebuild therefore evaluated WOT enrichment and gateway-cell count across 10 thresholds (25th, 50th, 75th, 90th, 95th, 99th, 99.5th, 99.9th, 99.95th, and 99.99th percentiles of *S*_gateway_), and recomputed bell-gene summaries for the high-stringency subset (90th, 95th, 99th, 99.5th, 99.9th, 99.95th, and 99.99th percentiles). At each threshold, cells scoring above the cutoff were designated gateway cells.

For each threshold, we recomputed differential expression between the resulting gateway population and both the upstream (MET) and iPSC reference populations (Mann–Whitney *U* test, Benjamini–Hochberg FDR < 0.05, |log1p LFC| > 0.25). Genes significantly upregulated in gateway cells relative to *both* populations were classified as **bell genes** (peaked expression at the transition); genes downregulated relative to both were classified as **valley genes**. The relationship between threshold stringency and bell-gene detection reflects a trade-off between biological specificity and statistical power (Extended Data Fig. 8c,d). Bell-gene yield varies with the cutoff, especially in 2i/LIF, but median per-gene effect sizes remain broadly stable across the 99th–99.95th percentile range. At the 99.95th percentile, the local rebuild recovers 83 serum cells and 85 2i/LIF cells, retaining enough power for downstream statistics while still focusing tightly on the boundary-enriched population.

Independent validation by the WOT enrichment odds ratio—measuring enrichment of gateway cells in the top decile of WOT backward-transport *P*(iPSC)—shows the same pattern. OR is near baseline below the 99th percentile, then rises sharply at high stringency (Extended Data Fig. 8a,b). At the 99.95th-percentile operating point, the OR is 29.2 in serum and 251.2 in 2i/LIF. Tightening further to the 99.99th percentile leaves only 17 cells per condition; serum OR increases further and 2i/LIF remains strongly enriched, but the very small sample limits robustness.

Beyond the 99.95th percentile, the number of cells drops to 17 per condition. Although the resulting cells remain strongly enriched for WOT iPSC fate, the small sample limits the stability of downstream gene-level calls. The 99.95th percentile therefore provides the most practical balance between strong orthogonal enrichment and adequate sample size.

Although the total number of bell genes varies with the threshold, the cross-condition overlap remains substantial throughout the high-stringency regime (Extended Data Fig. 8e). At the selected operating point (99.95th percentile), the serum and 2i/LIF analyses share 15 bell genes, and the overlap is larger at looser high-stringency cutoffs. Within this shared set, the epithelial gatekeeper *Sfn* remains present throughout, while tight-junction genes *Cldn6* and *Cldn7* persist through the selected operating point. *Mt1* and *Tuba1b* also remain part of the shared program at the 99.95th percentile, reinforcing the conserved epithelial/stress gateway signature.

The results above show that the 99.95th percentile provides a practical balance between specificity and sample size, and that the gateway framework is robust in the sense that the same core program persists throughout the 99th–99.99th percentile range even though the total bell-gene count changes with threshold. The choice can therefore be adapted to the analytical goal:

| Percentile | Cells | Bell genes | Use case |
| --- | --- | --- | --- |
| 99th–99.5th | 1,687–830 | 66–86 | Exploratory; larger cell set for subclustering or trajectory analysis |
| 99.9th | 166–169 | 44–149 | Moderate stringency; retains statistical power for most DE tests |
| **99.95th** | **83–85** | **60–199** | **Default; best balance of specificity and signal strength** |
| 99.99th | 17 | 67–234 | Ultra-stringent; very large per-gene effect sizes but small sample |

Importantly, the identity of the top-ranked gateway cells is consistent across thresholds: cells selected at the 99.99th percentile are a strict subset of those at the 99.95th, which are in turn a subset of those at the 99.9th. This nested structure means that tightening the threshold does not select a different population, but progressively focuses on the highest-scoring cells within the same boundary region of the latent space.

### Supplementary Note 4: Statistical power asymmetry between bell and valley gene detection

Bell genes (transiently upregulated at the gateway) and valley genes (transiently downregulated) are identified by symmetric statistical criteria: both require |LFC| > *τ* and FDR < *⍺* in both comparisons (gateway vs. source and gateway vs. target). Despite this symmetry in the test, the two classes face fundamentally asymmetric detection regimes. Here we derive this asymmetry from first principles, validate it empirically against the WOT reprogramming data, and conclude that bell genes provide the more statistically complete characterization of gateway biology.

#### Mathematical framework

##### Setup

Let *X*_*i*_ denote the log_2_(1+count)-transformed expression of gene *g* in cell *i*:

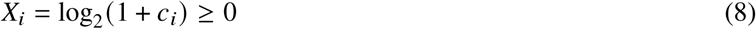

where *c*_*i*_ ≥ 0 is the raw UMI count. Expression is **bounded below** by zero and **unbounded above**.

Denote the population means in the three groups as *μ*_*G*_ (gateway, *n*_*G*_ cells), *μ*_*S*_ (source, *n*_*S*_ cells), and *μ*_*T*_ (target, *n*_*T*_ cells). The two log-fold-changes are

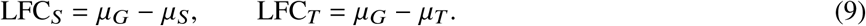

A gene is classified as *bell* if LFC_*S*_ > *τ*, LFC_*T*_ > *τ*, and both pass FDR < *⍺*. A gene is classified as *valley* if LFC_*S*_ < −*τ*, LFC_*T*_ < −*τ*, and both pass FDR < *⍺*. The *detection bottleneck* is

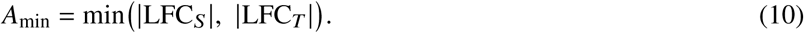

A gene is detected if and only if *A*_min_ > *τ* and both *p*-values survive FDR correction. (The reported amplitude uses the mean, *A* = (|LFC_*S*_ | + |LFC_*T*_ |)/2; the analysis below concerns the detection criterion, which depends on the minimum.)

##### Bounded downregulation, unbounded upregulation

- **Bell genes.** For a bell gene, *μ*_*G*_ > *μ*_*S*_ and *μ*_*G*_ > *μ*_*T*_. Since *μ*_*G*_can be arbitrarily large (no upper bound on expression), the amplitude *A*_min,bell_ = min(*μ*_*G*_− *μ*_*S*_, *μ*_*G*_ − *μ*_*T*_) is bounded only by the maximum achievable expression level: *A*_min,bell_ ∈ (0, ∞).
- **Valley genes.** For a valley gene, *μ*_*G*_ < *μ*_*S*_ and *μ*_*G*_ < *μ*_*T*_. Since *μ*_*G*_ ≥ 0 (Eq. 8),

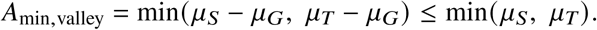

The valley detection bottleneck is **bounded above** by the expression level of the lower-expressing flanking population.

This is the core asymmetry:

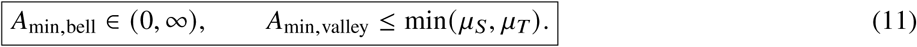

##### Consequence for the dual-comparison bottleneck

For a gene to be detected, *both* |LFC_*S*_ | > *τ* and |LFC_*T*_ | > *τ* must hold simultaneously. Define the *symmetry ratio*

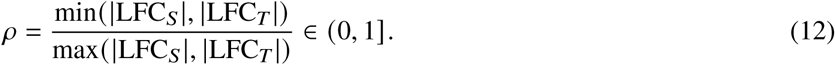

###### Proposition

For a valley gene with fixed total effect *E* = |LFC_*S*_ | + |LFC_*T*_ |, the detection bottleneck is *A*_min_ = *ρ* · *E*/(1 + *ρ*). The gene passes the threshold *τ* if and only if *ρ* > *τ*/(*E* − *τ*). As *E* decreases (due to the expression floor), the minimum required *ρ* increases—requiring more symmetric contrasts for detection.

###### Proof

Write |LFC_*S*_ | = *A*_min_ and |LFC_*T*_ | = *E* − *A*_min_ with *A*_min_ ≤ *E* − *A*_min_ (i.e., *A*_min_ is the bottleneck). Then *ρ* = *A*_min_/(*E* − *A*_min_), so *A*_min_ = *ρE*/(1 + *ρ*). The detection criterion *A*_min_ > *τ* becomes *ρ* > *τ*/(*E* − *τ*). Since valley genes have *E* ≤ *μ*_*S*_+ *μ*_*T*_ (bounded), and *E* is typically smaller for valley than bell candidates (empirically confirmed below), valley genes require higher *ρ* to pass—but empirically achieve lower *ρ*.

##### Zero-inflation amplifies the asymmetry

Single-cell expression data follows a zero-inflated distribution. Model the expression of gene *g* in population *k* as a mixture: *X* = 0 with probability *π*_*k*_; *X* = *Y*_*k*_ > 0 with probability 1 − *π*_*k*_. For a valley gene in gateway cells, *μ*_*G*_ is small, implying high dropout (*π*_*G*_ → 1). This has two consequences:

1. **Floor compression.** The maximum achievable |LFC| against a reference population with mean *μ*_ref_ is *μ*_ref_ − 0 = *μ*_ref_ (when *μ*_*G*_ = 0). This ceiling on valley LFC does not exist for bell genes.
2. **Rank-test power loss.** The Mann–Whitney *U* test operates on ranks. When *π*_*G*_ is high, many gateway cells share the value *X* = 0, creating ties. The asymptotic variance of the *U* statistic under tied ranks is

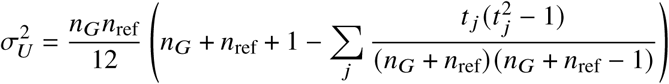

where *t* _*j*_ is the number of observations tied at rank *j*. Large tie groups from zero-inflated gateway cells reduce 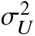 and thus the test’s ability to reject the null.

#### Empirical validation from WOT data

We test these predictions in the WOT MEF-to-iPSC reprogramming dataset, which contains 83 serum gateway cells and 85 2i/LIF gateway cells drawn from ∼165,000 total cells per condition, with ∼9,000 genes tested per condition.

##### Candidate pool asymmetry

Among genes significant in both comparisons (FDR < 0.05), the number of upregulated candidates (both LFCs > 0) far exceeds downregulated candidates (both LFCs < 0):

| Condition | Bell candidates | Valley candidates | Ratio |
| --- | --- | --- | --- |
| Serum | 407 | 117 | 3.5× |
| 2i/LIF | 1,765 | 102 | 17.3× |

##### LFC symmetry confirms the bottleneck

The symmetry ratio *ρ* (Eq. 12) is significantly lower for valley candidates than bell candidates in both conditions:

| Condition | Bell $\rho$ (median) | Valley $\rho$ (median) | $P$ (MW) |
| --- | --- | --- | --- |
| Serum | 0.907 | 0.722 | $2.1 \times 10^{-18}$ |
| 2i/LIF | 0.794 | 0.653 | $6.4 \times 10^{-7}$ |

Valley genes are strongly downregulated versus one flanking population but only modestly below the other. The weaker contrast is the bottleneck that prevents detection. For bell genes, both contrasts are nearly equal (*ρ* ≈ 0.9), meaning the gateway uniquely expresses these genes at levels exceeding both flanking populations.

##### Near-miss accumulation

Valley genes accumulate just below the threshold. Define the *near-miss ratio* as the number of genes with *A*_min_ ∈ (0.75*τ*, *τ*) divided by the number with *A*_min_ > *τ*. In serum, this ratio is 1.28 for valley candidates versus 0.68 for bell candidates (1.9×), consistent with valley genes accumulating at the detection boundary.

##### Valley genes are genuine

Despite the detection disadvantage, the detected valley genes represent robust biological signal by three criteria. First, all 93 valley genes (58 serum, 46 2i; 11 shared) achieve combined significance *P* < 10^−3^ by Fisher’s method, with median combined *P* < 10^−10^. Second, of the 58 serum valley genes, a majority show concordant downregulation (both LFCs < 0) in the 2i condition. Third, the 11 shared valley genes, including *Cox6b2*, *Rpl10*, and *Uba52*, yield Fisher’s exact OR = 57.9 (*P* < 10^−12^).

#### Conclusion

The dual-comparison framework creates an inherent statistical asymmetry: bell gene detection benefits from unbounded upregulation, symmetric LFCs (*ρ* ≈ 0.9), and a large candidate pool; valley gene detection is constrained by the expression floor (Eq. 11), produces asymmetric LFCs (*ρ* ≈ 0.65), and draws from a 3.5–17.3× smaller candidate pool. We retain identical thresholds for both classes for methodological consistency, and report all detected valley genes for completeness. However, we focus biological interpretation primarily on bell genes, which provide a more statistically powered and biologically coherent characterization of the molecular programs active at fate boundaries.

### Supplementary Note 5: Soft-threshold gateway gene analysis

The standard gateway gene analysis uses a hard percentile threshold (99.95th percentile of diffused gateway scores) to define a discrete set of gateway cells, then applies Mann–Whitney *U* tests to identify bell and valley genes. While effective (Supplementary Note 3), this approach raises a basic question: if we avoid binary gateway membership, can we still detect bell and valley genes that distinguish the gateway from its flanking populations? If not, the gateway cells could simply be an artifact of the cutoff. Here we describe a complementary soft-threshold approach. Instead of a binary gateway membership rule, we compute a weighted gateway-side expression profile from a compact soft-core of gateway-score cells and compare it with the flanking source and target populations. The answer is yes: the two methods recover the same boundary biology and largely overlapping top-ranked genes.

Rather than averaging across the full positive-score zone, we define a restricted *soft-core* from the upper tail of positive diffused gateway-score cells within each condition (here, the top 5%). This yields 228 serum cells from the 4,550-cell positive-score zone and 274 2i/LIF cells from the 5,479-cell positive-score zone. Each cell *i* in the soft-core receives a weight proportional to its diffused gateway score: *w*_*i*_ = *s*_*i*_/Σ_*j*_ *s*_*j*_, where *s*_*i*_ is the diffused gateway score. In practice, the gateway-side expression mean becomes a weighted average: higher-scoring soft-core cells contribute more to each gene’s gateway mean, while lower-scoring soft-core cells still contribute but with less influence. This retains continuity within a compact transition band while avoiding the much broader positive-score halo used in the separate scVI comparison of Supplementary Note 6.

For each gene, we compute weighted log-fold-changes between the weighted soft-core mean and the unweighted reference populations (MET source and iPSC target cells, randomly subsampled to *n* ≤ 5,000). Significance is assessed by a weighted Welch *t*-test with Bessel-corrected weighted variance and Welch–Satterthwaite degrees of freedom. The effective soft-core sample size is 117.1 in serum and 213.9 in 2i/LIF. Bell and valley genes are classified using the same criteria as the standard analysis (|LFC| > 0.25, FDR < 0.05 in both comparisons).

The restricted soft-core analysis identifies 55 bell and 61 valley genes in serum, compared with 60 bell and 58 valley genes from the hard-threshold analysis. In 2i/LIF it identifies 133 bell and 22 valley genes, compared with 199 bell and 46 valley genes from the hard-threshold analysis.

In serum, 40 bell genes are shared between hard and soft-core analyses (Jaccard index = 0.53). Concordance is stronger at the top of the rank list: 60% overlap in both the top 10 and top 20. The leading serum shared bell genes remain *Krt8*, *H19*, *Mt1*, *Car2*, and *Sfn*. For these shared serum bell genes, amplitude is correlated between methods (Spearman *ρ* = 0.61; Extended Data Fig. 9g,h).

In 2i/LIF, 104 bell genes are shared between hard and soft-core analyses (Jaccard index = 0.46), with 50% top-10 overlap and 55% top-20 overlap. Amplitude correlation for shared 2i/LIF bell genes is also retained (Spearman *ρ* = 0.51; Extended Data Fig. 9g,h).

Valley-gene concordance is weaker than bell-gene concordance under the restricted soft-core: 39 serum valley genes and 19 2i/LIF valley genes are shared between hard and soft-core analyses (Extended Data Fig. 9i).

The soft-only and hard-only genes tend to have amplitudes closer to the detection threshold, consistent with borderline genes that cross or miss the significance cutoff depending on the exact restricted weighting scheme. Overall, this analysis supports the same boundary biology while showing that the broad positive-score halo should not be conflated with the compact soft-core reported in Extended Data Fig. 9.

The soft-threshold analysis confirms that the core gateway gene program is robust to the choice of thresholding strategy. The top-ranked bell and valley genes are recovered regardless of whether a hard percentile cutoff or compact soft-core weighting is used, and effect-size estimates are correlated (Extended Data Fig. 9). This concordance supports the detection of gateway populations with distinct programs rather than populations created by one exact cutoff. We nevertheless retain the hard-threshold definition in the main text because it provides a discrete operating set of gateway cells for gene calling, figures, and cross-dataset comparisons.

### Supplementary Note 6: WOT scVI baseline comparison

To compare scVI and scAttnVI fairly on the WOT benchmark, we retrained plain scVI on the same filtered serum and 2i/LIF datasets used in the main analysis, using the same gene matrix, latent dimension, and optimization settings but setting *λ*_BMI_ = 0. Gateway scores were then computed with the same downstream pipeline used for scAttnVI: identical source and target annotations, the same *k*NN-based pull score, the same PPR diffusion parameters, and the same hard-threshold percentile within each condition. Because the two models learn different latent spaces, UMAPs were generated separately for visualization; the comparison in Extended Data Fig. 11 therefore asks whether each model isolates a compact transition zone and recovers the associated bell-gene signal, not whether the two embeddings are geometrically identical.

Plain scVI still revealed a MET–iPSC transition corridor in both serum and 2i/LIF (Extended Data Fig. 11a–d). The broad transition is therefore not created de novo by BMI regularization. The difference emerges in how sharply that transition is resolved. Under soft scoring, the positive-score zone expanded to 93,489 cells in serum and 158,608 cells in 2i/LIF for scVI, compared with 4,550 and 5,479 cells for scAttnVI. scVI thus captured the overall reprogramming corridor, but it spread that signal across a much broader population.

At the hard threshold, both models return the same number of gateway cells by construction because the percentile cutoff is matched within each condition. Bell-gene recovery is therefore the informative comparison. In serum, scVI recovered 8 hard bell genes, whereas scAttnVI recovered 60; in 2i/LIF, it recovered 57 versus 199 (Extended Data Fig. 11e). For the soft comparison in Extended Data Fig. 11f, we used the full positive-score transition zone (*s*_*i*_ > 0 after diffusion) with gateway-score weights, rather than the restricted soft-core analysis used in Extended Data Fig. 9. Under this full-zone weighted analysis, scVI recovered 2 versus 51 bell genes in serum and 1 versus 65 in 2i/LIF. These results indicate that reconstruction alone is sufficient to reveal the broad transition, whereas BMI regularization improves the local resolution needed to separate a compact gateway population and recover the transient genes that peak there.

### Supplementary Note 7: Comparison with CellRank and Palantir in WOT

CellRank and Palantir provide an important point of reference because they recover the same broad reprogramming landscape that gateway analysis interrogates. Palantir places cells on a diffusion manifold, orders them from an early root, and estimates branch probabilities and entropy with respect to terminal outcomes. CellRank builds a directed Markov chain on the cell graph and computes absorption probabilities to terminal states. These probabilities can likewise be summarized as entropy. In both methods, high entropy marks cells whose eventual fate remains unresolved. For WOT, the biologically appropriate terminal comparison is stromal versus iPSC, because this is the dominant successful-versus-failed split described in the original reprogramming study. MET is therefore treated as an intermediate state on the route to reprogramming rather than as a terminal fate. This makes the comparison informative but not identical. A high-entropy CellRank or Palantir cell is not, by construction, a MET–iPSC gateway cell. It is a cell with uncertain terminal outcome. Gateway analysis asks a more localized question: which rare cells lie specifically at the interface between two annotated flanking states, here MET and iPSC, and which genes peak only in that interval.

Extended Data Fig. 12 shows both the common ground and the added resolution. In serum (Extended Data Fig. 12a–c), Palantir partially recovers the same MET–iPSC corridor as the hard gateway, with a Jaccard overlap of 0.169 and a median backward WOT *P*(iPSC) of 1.03 × 10^−5^, indicating partial but incomplete recovery of the fate-enriched corridor. Palantir also recovers a related transient program, including *Krt8*, *Mt1*, *H19*, *Car2*, *Lgals1*, and *Tpm1*, and shares 9 of the top 20 bell genes with the gateway result. CellRank is weaker in serum, with a Jaccard overlap of 0.012, a lower median *P*(iPSC) of 7.79 × 10^−6^, only 12 bell genes, and 1 of the top 20 bell genes shared with gateway analysis. These serum results support the reality of the transition corridor and show that gateway cells lie within a bona fide region of elevated fate uncertainty.

The distinction becomes clearer in 2i/LIF (Extended Data Fig. 12d–j). Here, neither CellRank nor Palantir recovers the same localized hard gateway cell set, and both have zero cell-level overlap with the gateway under the matched-size selection used in this analysis. Palantir remains weakly fate-enriched, with a median backward WOT *P*(iPSC) of 4.99 × 10^−6^ and only 12 bell genes, while CellRank broadens into a distinct high-entropy field with a lower median *P*(iPSC) of 3.57 × 10^−6^. Most importantly, the recovered gene programs diverge. The 2i/LIF gateway retains a large bell-gene set (199 genes) headed by *Mt2*, *Mt1*, *Mylpf*, *Tdh*, *Dppa3*, and *Tuba1b*, whereas CellRank is dominated by broader stromal- and matrix-associated genes such as *Col1a1*, *Col1a2*, *Bgn*, *Sparc*, and *Timp3*, and Palantir yields only a small, distinct bell set led by *Tuba1a*, *Rangrf*, and *Atp5k*.

This added resolution is also reflected in cross-condition coherence (Extended Data Fig. 12j). The serum and 2i/LIF gateway analyses share 15 bell genes overall and 4 of their top 20 bell genes (*Mt1*, *Sfn*, *Dstn*, and *Tuba1b*). By comparison, Palantir shares 1 bell gene overall and 1 of its top 20 bell genes across conditions, and CellRank shares 0 bell genes overall and 0 of its top 20 bell genes. Gateway analysis is therefore not only more localized within each condition, but also more coherent across reprogramming contexts. Together, these comparisons show that gateway cells are not an isolated artifact of one model. They sit within the same transition corridor identified by global fate-mapping methods. Gateway analysis, however, resolves that corridor into a rarer and more coherent local interface with a sharper transient gene program.

---

**Extended Data Fig. 1.**
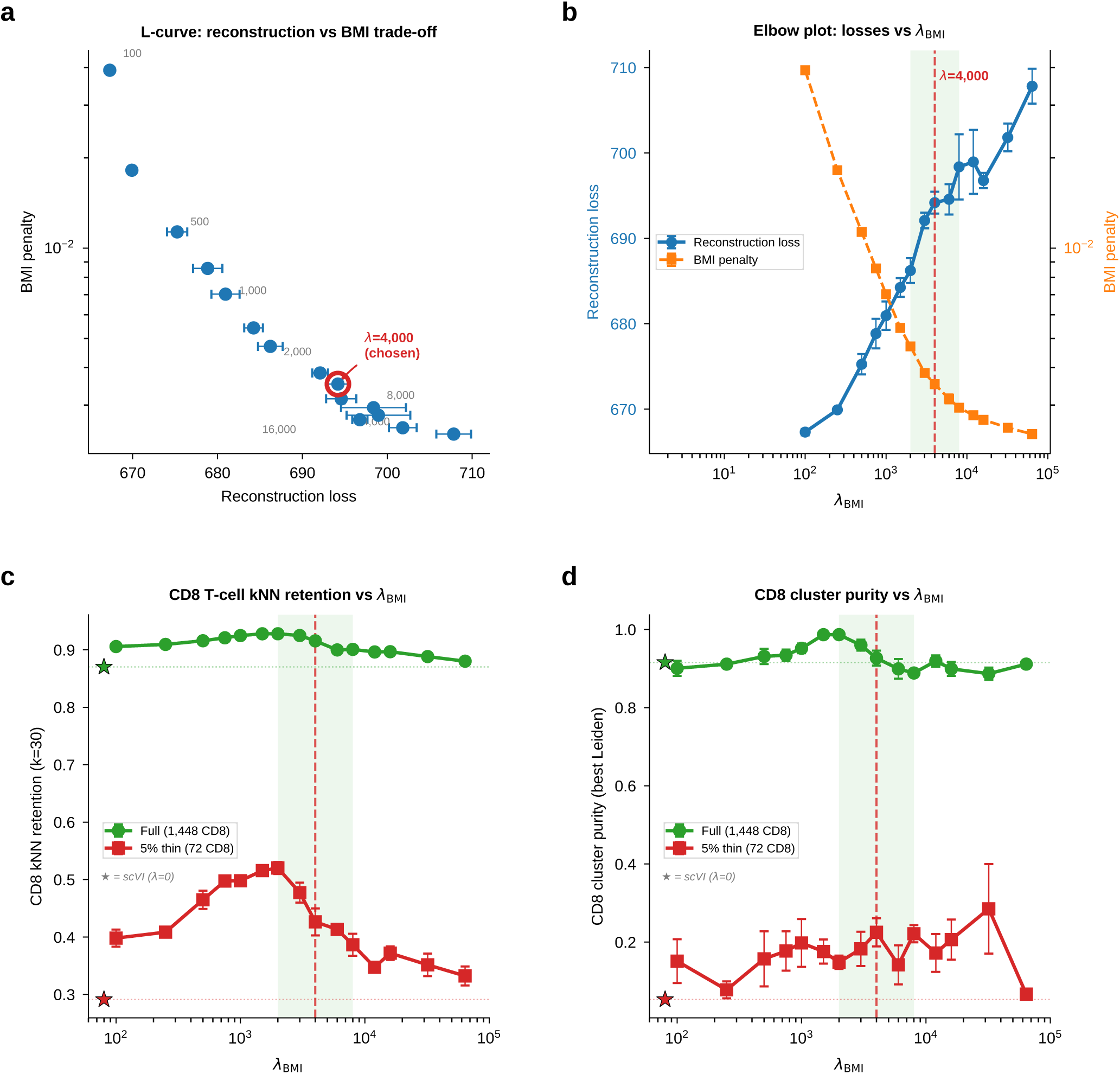
*λ*_BMI_ penalty sweep on PBMC. **(a)** L-curve showing the trade-off between reconstruction loss and BMI penalty as *λ*_BMI_ increases from 100 to 64,000. **(b)** Dual-axis elbow plot: reconstruction loss increases monotonically with *λ*, while BMI penalty decreases. **(c)** CD8 T-cell *k*NN retention across *λ* values at full retention and 5% thinning. **(d)** CD8 cluster purity across *λ* values. The chosen *λ* = 4,000 sits in the elbow range and preserves rare-cell geometry under thinning.

**Extended Data Fig. 2.**
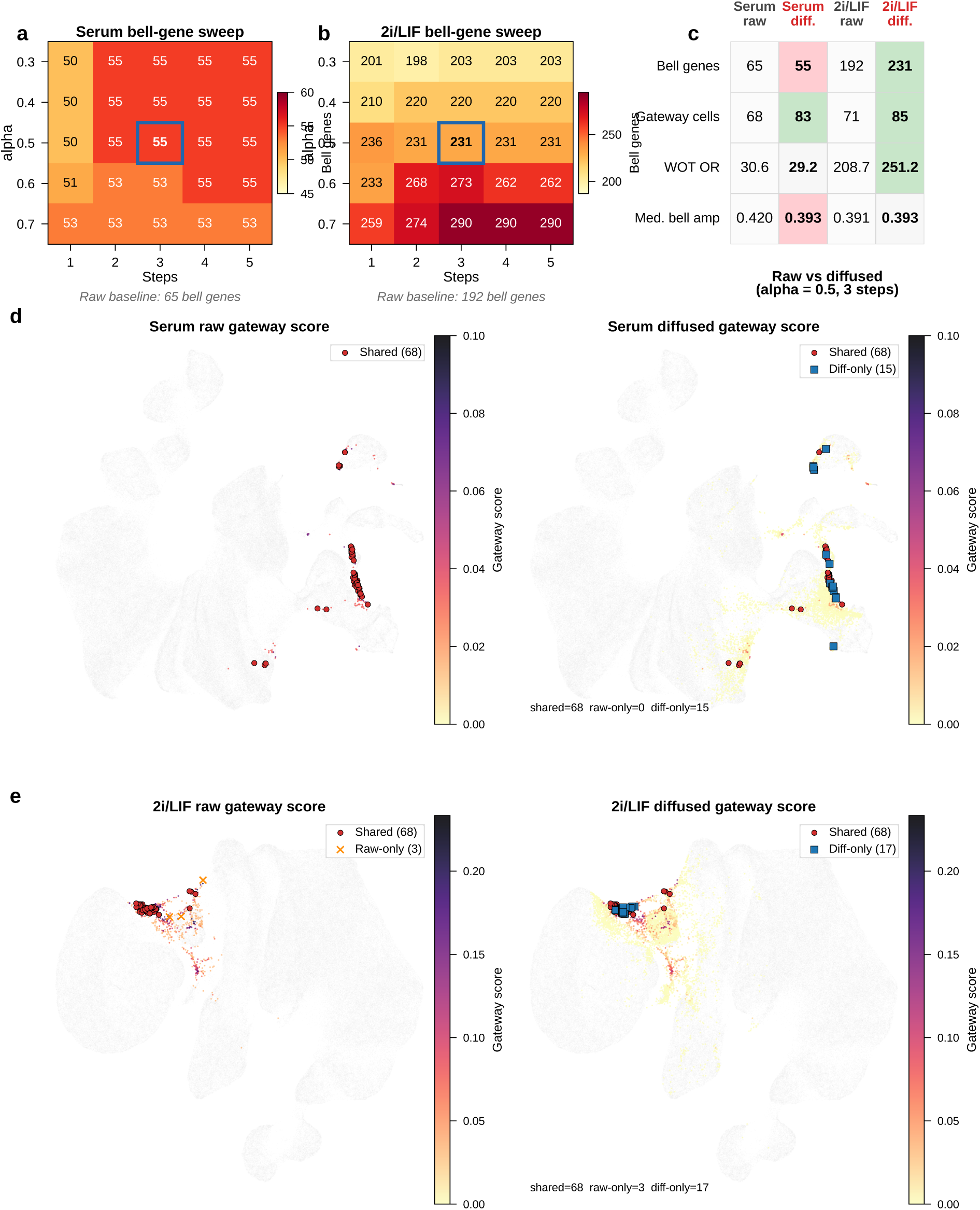
Personalized PageRank diffusion on kNN-in-latent-space gateway scores. **(a,b)** Heatmaps of statistically significant bell-gene count across diffusion parameters for serum (a) and 2i/LIF (b). Blue rectangle: chosen configuration (*⍺* = 0.5, 3 steps). Raw baseline bell-gene counts are shown beneath each heatmap. **(c)** Quantitative comparison of raw and diffused gateway scores at the selected operating point across bell-gene count, gateway-cell count, WOT top-decile enrichment odds ratio, and median bell-gene amplitude. **(d,e)** UMAP colored by gateway score before (left, raw) and after (right, diffused) for serum (d) and 2i/LIF (e). Gateway cells are defined at the 99.95th-percentile threshold for each score. Shared cells are shown in red; raw-only cells in orange; diffused-only cells in blue. Diffusion smooths the sparse raw scores along the latent *k*NN manifold while preserving spatial specificity at the MET–iPSC boundary.

**Extended Data Fig. 3.**
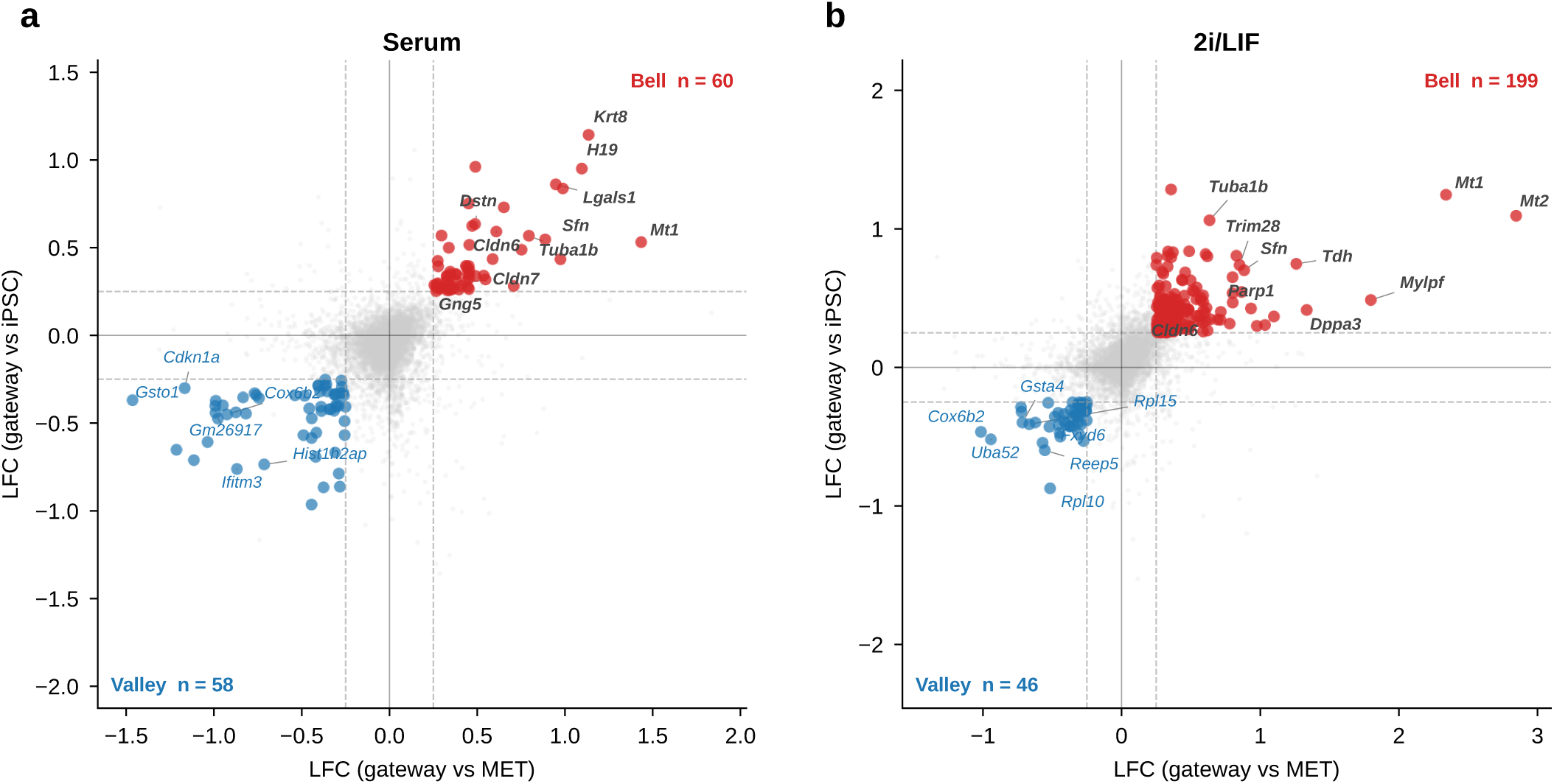
Quadrant classification of gateway-specific gene programs. **(a,b)** Scatter plots of log-fold changes for all tested genes in serum (a; 9,448 genes) and 2i/LIF (b; 8,822 genes). Each point represents a gene; the *x*-axis shows log_1*p*_ fold change of gateway cells versus MET and the *y*-axis shows log_1*p*_ fold change versus iPSC. Genes in the upper-right quadrant (red) are bell genes and genes in the lower-left quadrant (blue) are valley genes. Labeled genes are the shared bell and valley genes conserved across both conditions. Serum: 60 bell, 58 valley; 2i/LIF: 199 bell, 46 valley.

**Extended Data Fig. 4.**
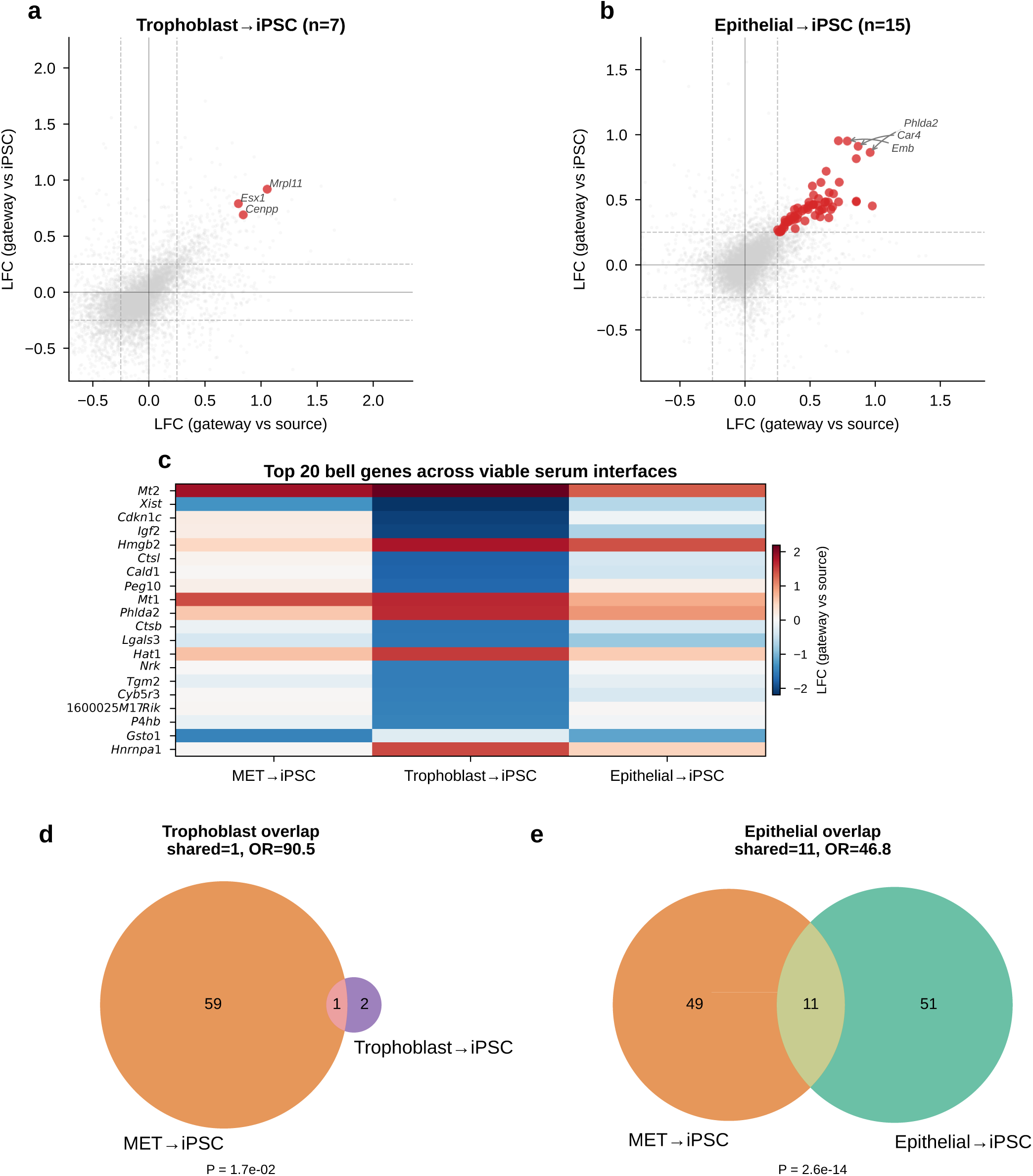
Non-MET gateway interfaces carry distinct bell programs. **(a,b)** Four-quadrant scatter plots of log-fold changes for gateway cells at non-MET interfaces in serum: Trophoblast→iPSC (**a**; *n* = 7 gateway cells; 14 bell and 2 valley genes) and Epithelial→iPSC (**b**; *n* = 15 gateway cells; 59 bell genes). No non-MET interfaces were detected in 2i/LIF. **(c)** Heatmap of bell-gene LFC across the MET→iPSC reference and viable non-MET interfaces. **(d,e)** Venn diagrams of bell-gene overlap with MET→iPSC, showing no overlap for trophoblast and limited overlap for epithelial. Gateway scores were computed using kNN fate pulls (*k* = 30) with PPR diffusion (*⍺* = 0.5, 3 steps, 99.95th-percentile threshold).

**Extended Data Fig. 5.**
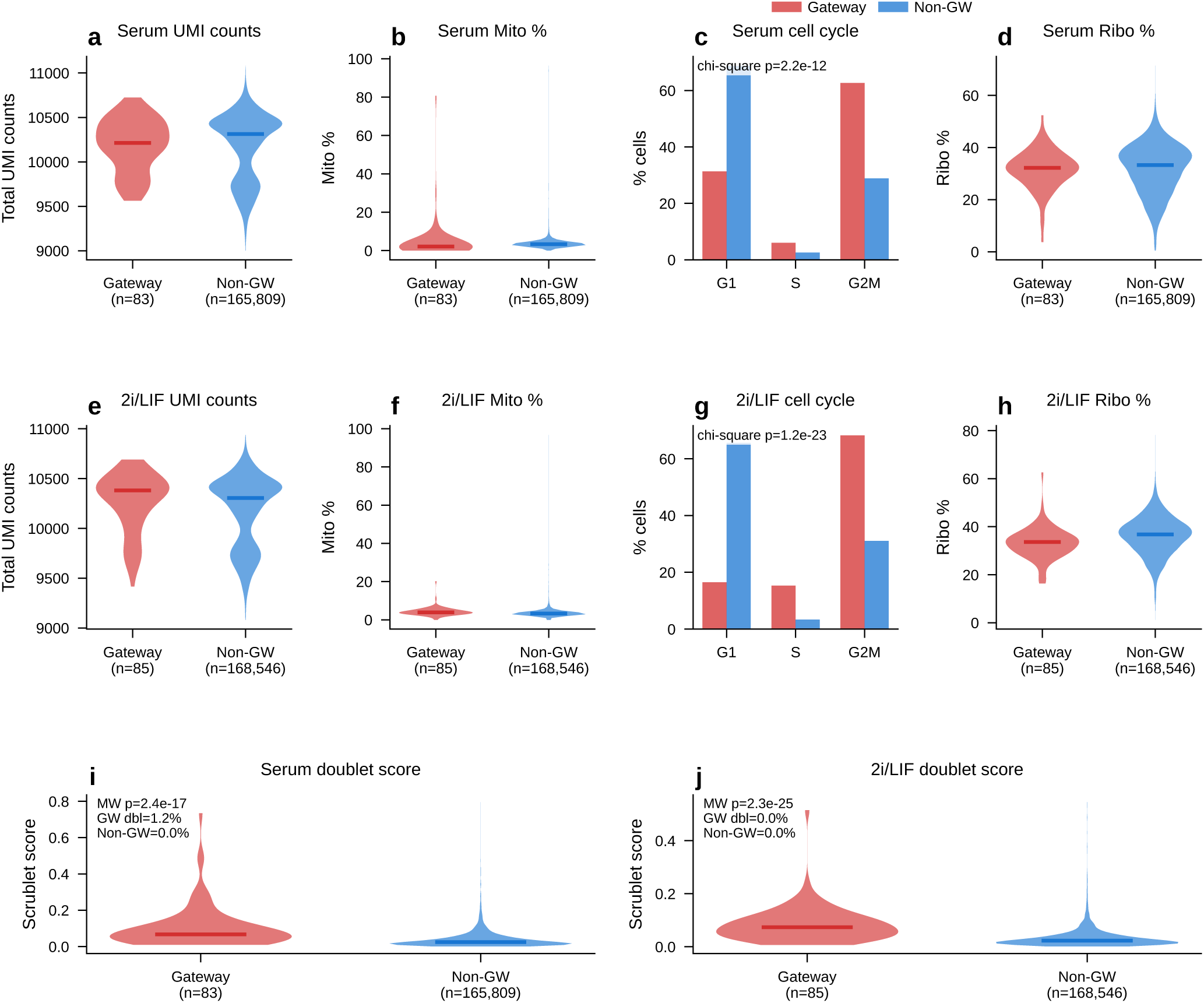
Technical confound controls and doublet analysis for gateway cells. **(a–d)** Serum confound controls comparing gateway (*n* = 83, red) versus non-gateway (*n* = 165,809, blue) cells: total UMI counts, mitochondrial read fraction, cell-cycle phase distribution, and ribosomal read fraction. **(e–h)** Equivalent analyses for 2i/LIF (*n* = 85 gateway cells, *n* = 168,546 non-gateway cells). **(i,j)** Scrublet^22^ doublet score distributions for gateway versus non-gateway cells in serum (i) and 2i/LIF (j). Predicted doublet rates among gateway cells remain very low overall (1.2% in serum; 0.0% in 2i/LIF). Mann–Whitney *U* test (two-sided) for continuous and doublet-score panels; *χ*^2^ test for cell-cycle phase distributions.

**Extended Data Fig. 6.**
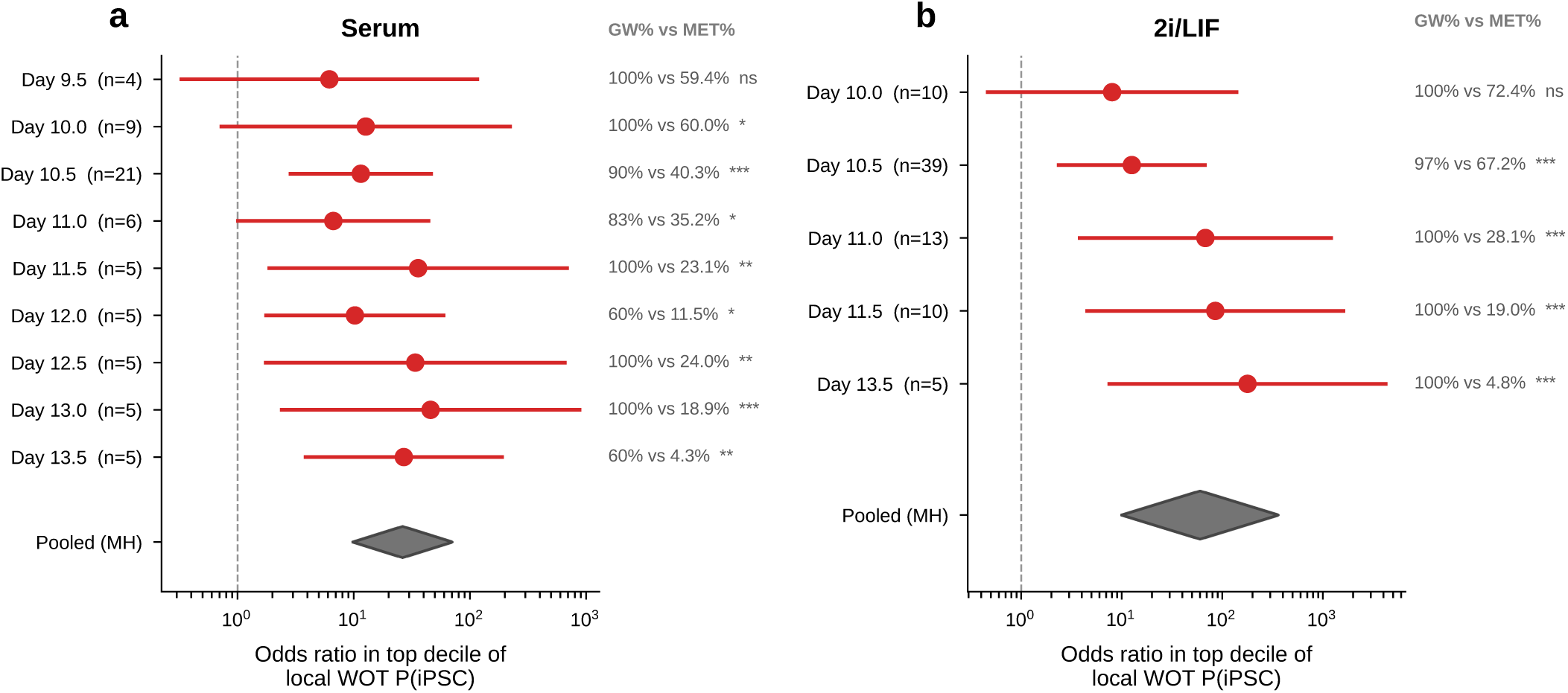
Per-day WOT enrichment of gateway cells. **(a,b)** Forest plots of odds ratios for gateway cells versus same-day MET controls falling in the top decile of WOT backward-transport *P*(iPSC), for serum (a) and 2i/LIF (b). Dots and horizontal lines show OR with 95% CIs; diamonds show the pooled Mantel–Haenszel estimate. Gateway cells are enriched at every time point tested in both conditions.

**Extended Data Fig. 7.**
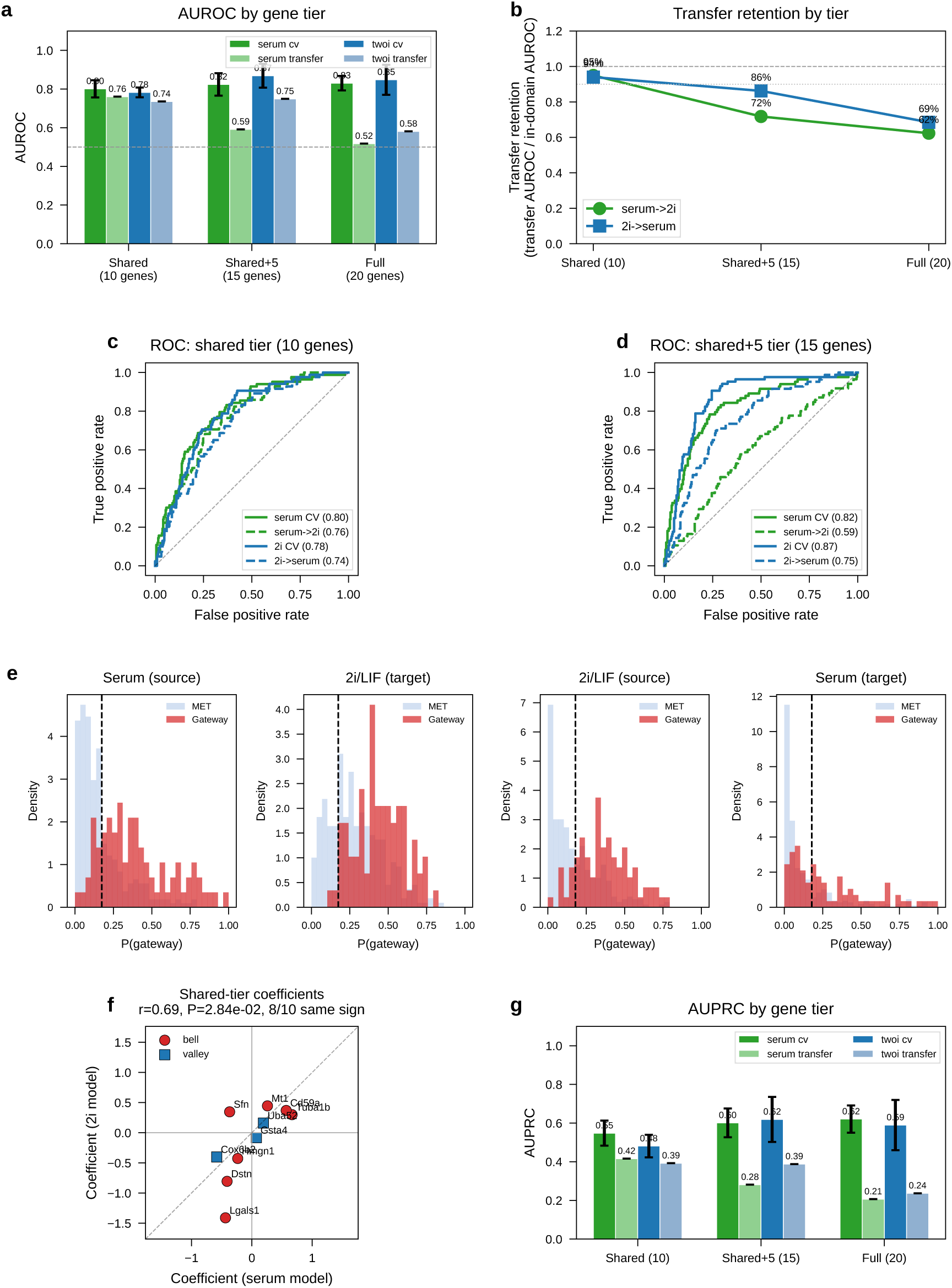
Cross-condition gateway classifier transfer. **(a)** AUROC for in-domain 5-fold CV versus cross-condition transfer across three gene tiers: shared-only (10 genes: top 7 shared bell + top 3 shared valley by cross-condition minimum amplitude), shared+5 (15 genes), and source full20 (20 genes). **(b)** Transfer retention ratio (transfer AUROC / in-domain AUROC), showing strongest retention for the shared tier and lower retention after adding source-specific genes. **(c,d)** ROC curves for shared and shared+5 tiers, comparing in-domain (solid) versus transfer (dashed). **(e)** Score distributions for gateway and day-matched MET control cells in source and target conditions for the shared tier. **(f)** Coefficient correlation between serum and 2i shared-tier models. **(g)** AUPRC by gene tier, complementing AUROC for imbalanced classification.

**Extended Data Fig. 8.**
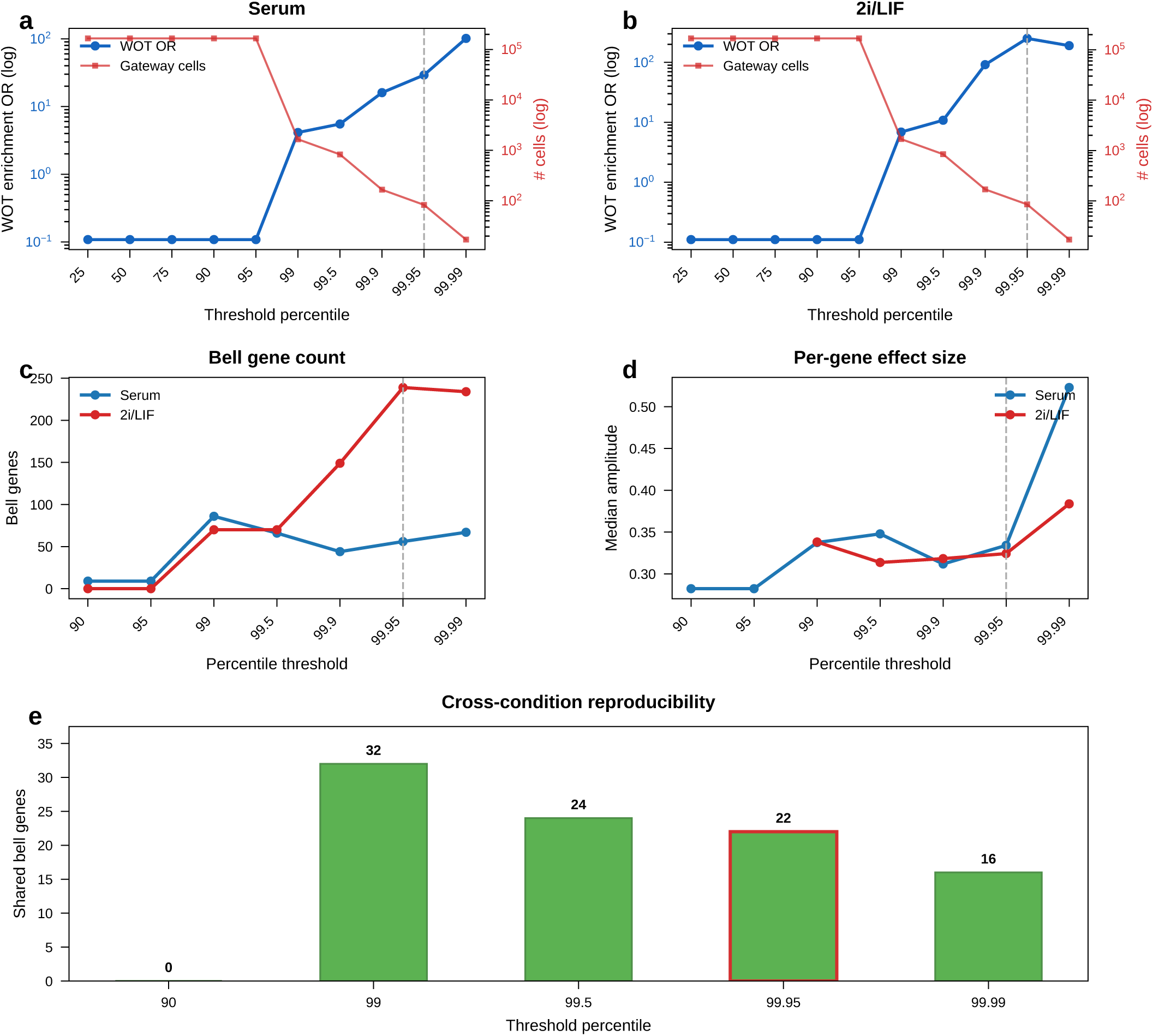
Gateway score threshold selection and robustness. **(a,b)** WOT enrichment odds ratio (OR, blue, left log axis) and gateway cell count (red, right log axis) as a function of threshold percentile for serum (a) and 2i/LIF (b). The dashed line marks the 99.95th-percentile operating point. **(c)** Bell gene count across tested thresholds for serum and 2i/LIF. Counts vary with threshold, especially in 2i/LIF, but remain substantial across the 99th–99.99th regime. **(d)** Median per-gene bell amplitude across the same thresholds. Effect sizes remain broadly stable across informative thresholds, with the sharpest changes only at the most extreme cutoff. **(e)** Number of shared bell genes between serum and 2i/LIF across tested thresholds. Shared overlap persists across the 99th–99.99th regime; *Sfn* is shared throughout, and *Cldn6*/*Cldn7* remain shared through the 99.95th-percentile operating point.

**Extended Data Fig. 9.**
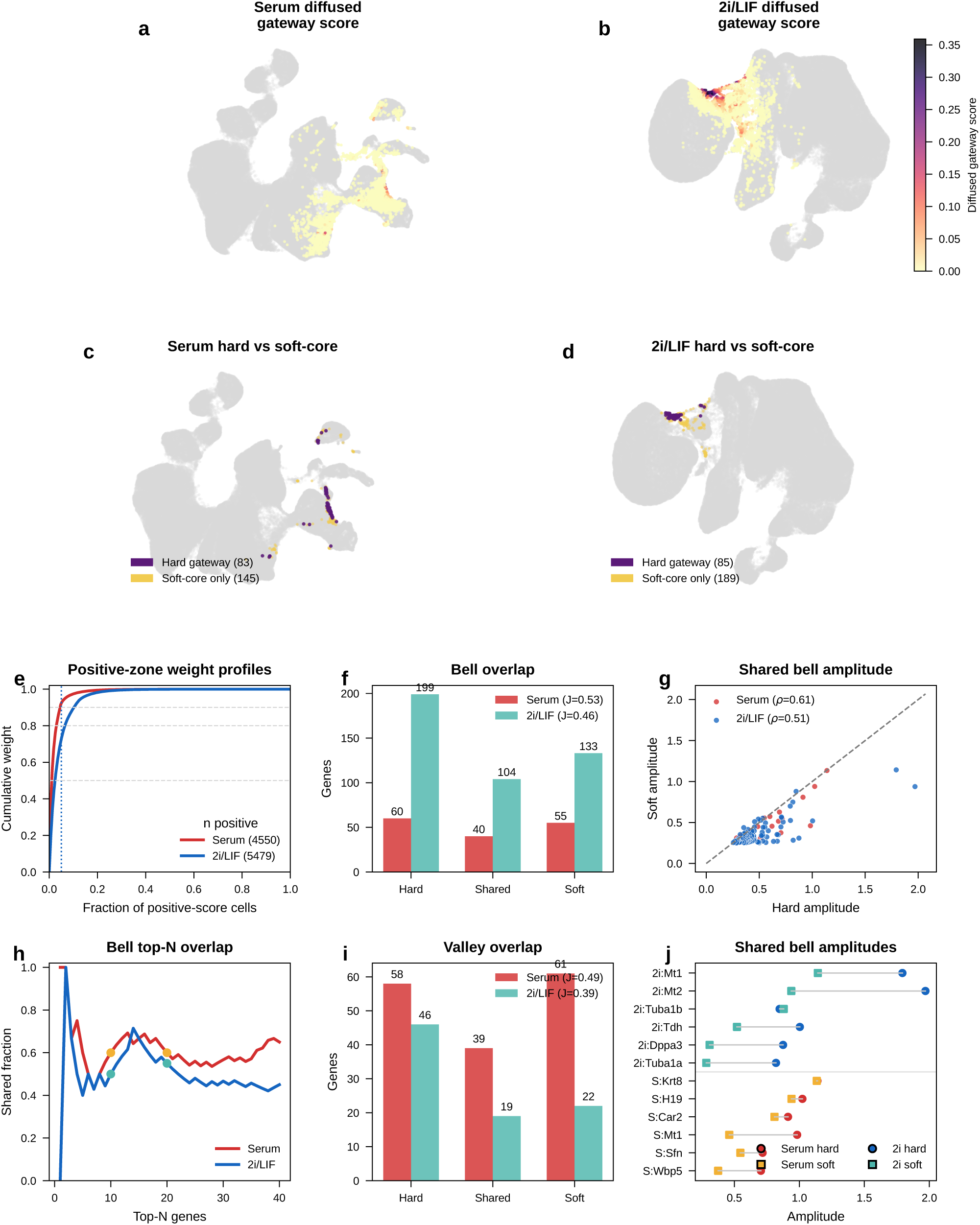
Soft-core gateway gene analysis. **(a,b)** UMAPs of the serum (a) and 2i/LIF (b) WOT BMI latent spaces colored by positive diffused gateway score. **(c,d)** Comparison of hard-threshold gateway cells versus the restricted positive-score soft-core in serum (c) and 2i/LIF (d). **(e)** Cumulative weight profiles across all positive-score cells in serum and 2i/LIF; dotted lines mark the restricted soft-core cutoff used for the continuous analysis. **(f)** Bell-gene overlap summary between hard-threshold and soft-core analyses for serum and 2i/LIF; legend reports Jaccard overlap within each condition. **(g)** Correlation of bell-gene amplitudes shared between hard and soft-core analyses in serum and 2i/LIF. **(h)** Top-*N* rank overlap curves for bell genes in serum and 2i/LIF. **(i)** Valley-gene overlap summary between hard-threshold and soft-core analyses for both conditions. **(j)** Paired hard-versus-soft-core amplitudes for representative shared bell genes in serum and 2i/LIF.

**Extended Data Fig. 10.**
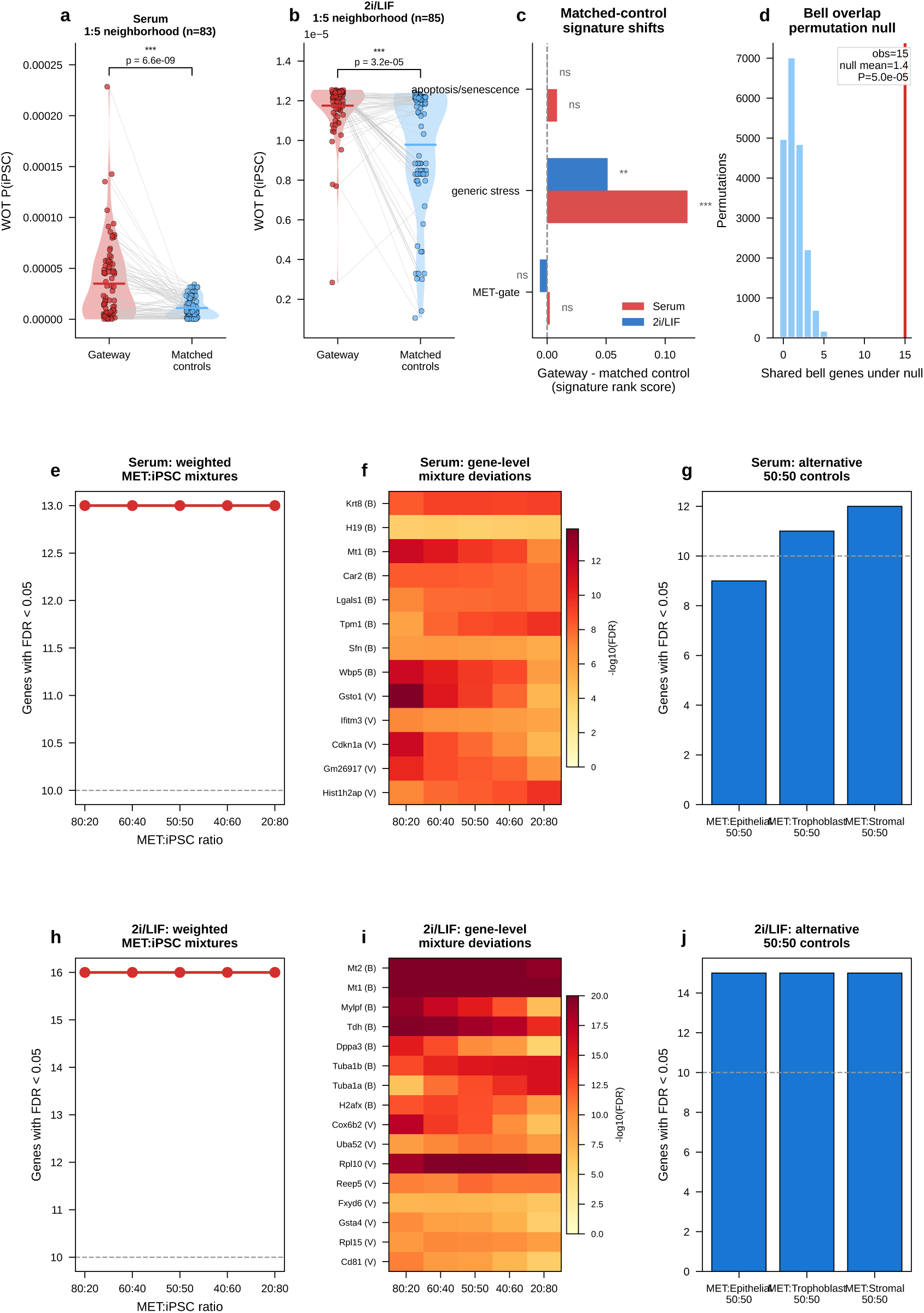
Synthetic-mixture and orthogonal WOT robustness controls. **(a,b)** Gateway cells versus 1:5 latent-neighborhood matched controls for serum (a) and 2i/LIF (b), scored by independent WOT backward-transport *P*(iPSC). **(c)** Gateway-minus-control signature shifts for MET-gate, generic stress, and apoptosis/senescence signatures in serum and 2i/LIF, using the same 1:5 latent-neighborhood matching. **(d)** Permutation test for overlap between serum and 2i/LIF bell genes. The observed overlap exceeds the null distribution. **(e–g)** Added local serum weighted-mixture controls: number of significant gateway-panel genes across MET:iPSC ratios (e), gene-level deviations across those ratios (f), and alternative 50:50 controls using MET mixed with epithelial, trophoblast, or stromal cells (g). **(h–j)** Analogous weighted-mixture controls in 2i/LIF.

**Extended Data Fig. 11.**
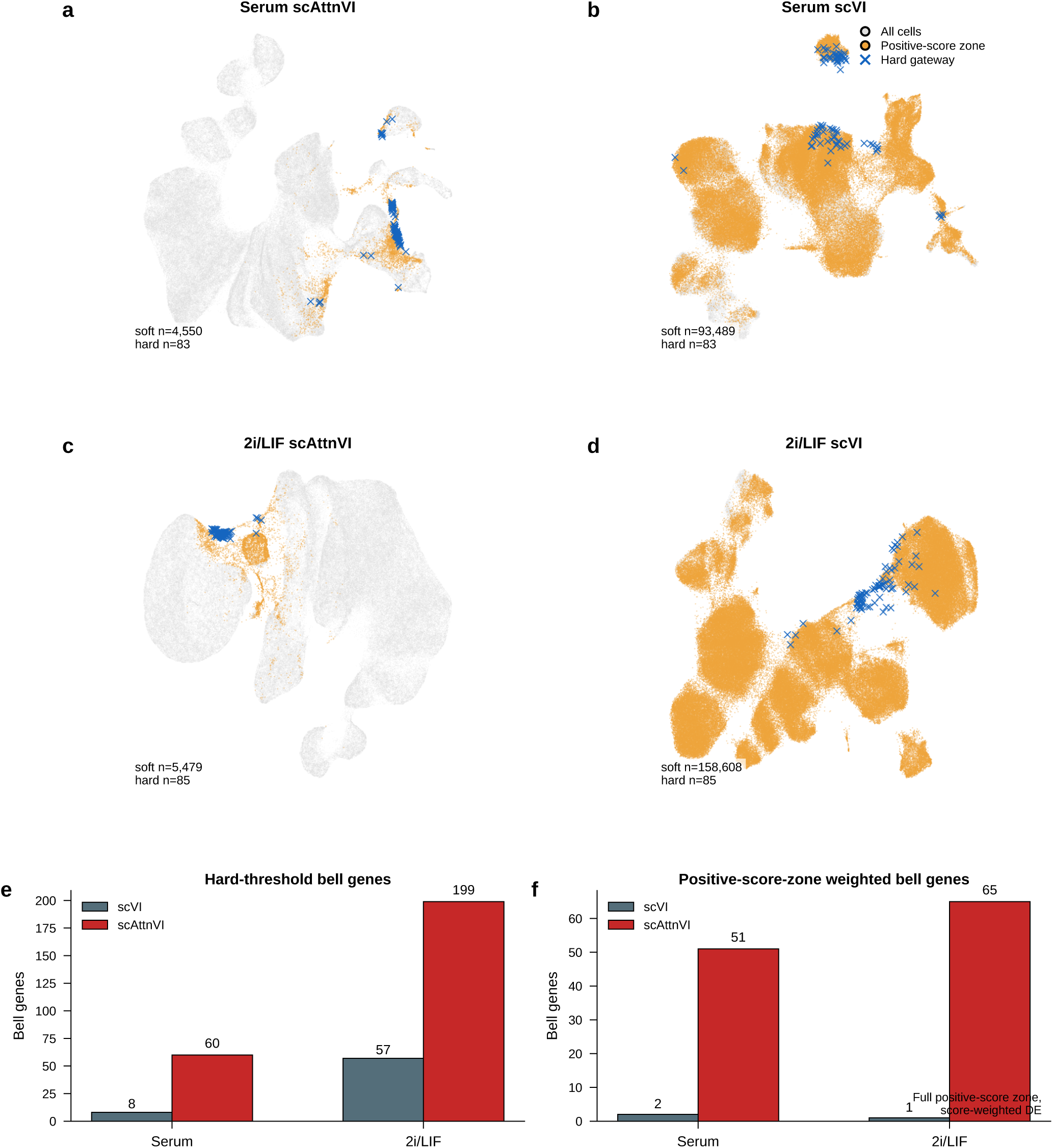
WOT scVI versus BMI-regularized scAttnVI. **(a–d)** Model-specific UMAPs for serum and 2i/LIF. Gray points denote all cells, orange points the positive-score transition zone, and blue crosses the hard gateway cells. Both models highlight a MET–iPSC transition region, but the scVI positive-score zone is markedly more diffuse. **(e)** Hard-threshold bell-gene recovery. In the current analysis, scAttnVI recovers substantially more hard bell genes than scVI in both serum (60 versus 8) and 2i/LIF (199 versus 57). **(f)** Full positive-score-zone weighted bell-gene recovery. Using the entire positive-score transition zone with gateway-score weights, scAttnVI recovers 51 versus 2 bell genes in serum and 65 versus 1 in 2i/LIF. This definition differs from the restricted soft-core analysis used in Extended Data Fig. 9.

**Extended Data Fig. 12.**
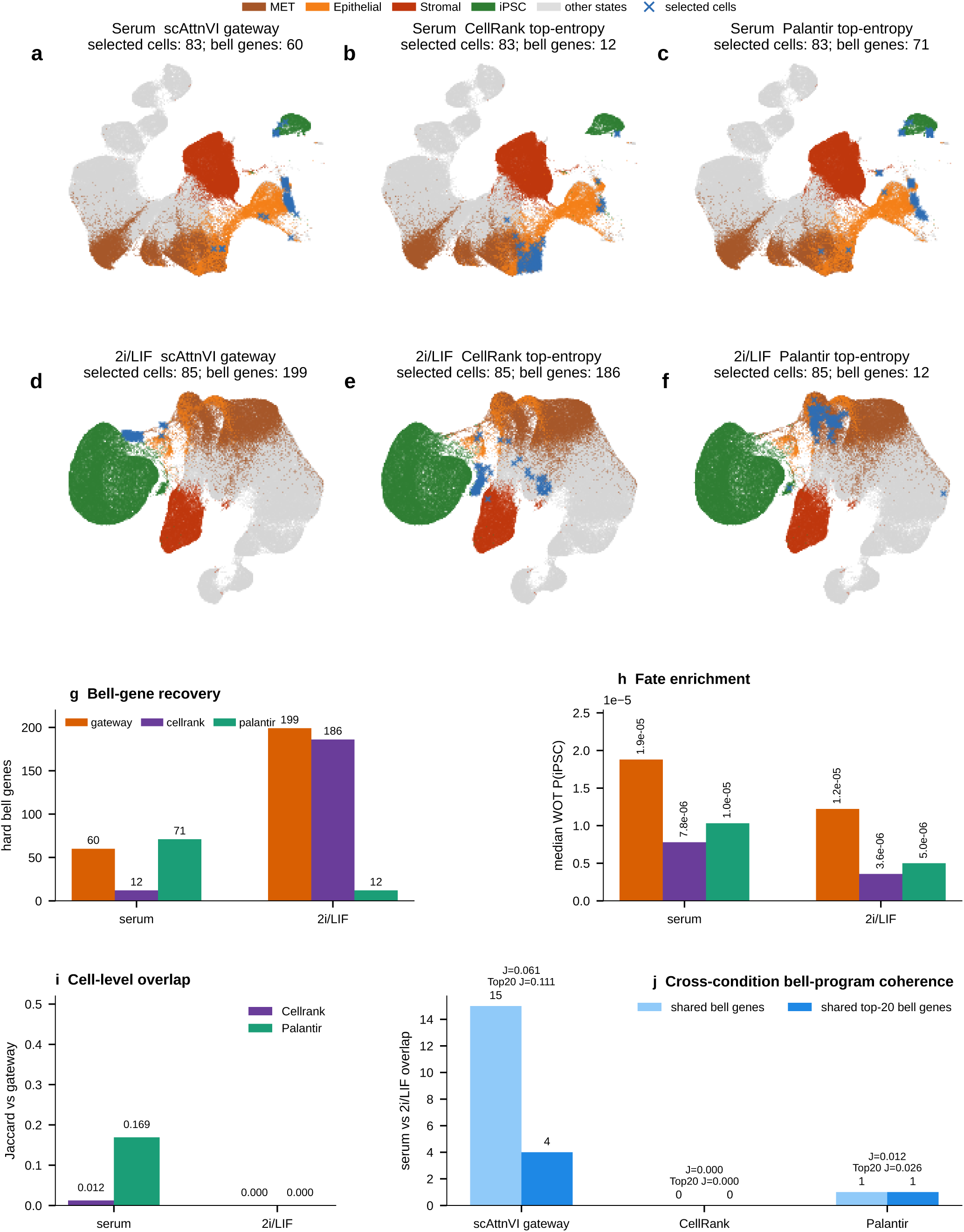
Comparison of the WOT MET–iPSC gateway with CellRank and Palantir. **(a–c)** Serum WOT UMAP showing the hard scAttnVI gateway, top-entropy CellRank cells, and top-entropy Palantir cells, each matched to the same hard cell count. Palantir partially overlaps the serum gateway corridor, whereas CellRank is more diffuse. **(d–f)** Analogous comparison in 2i/LIF. In this condition, neither CellRank nor Palantir recovers the same localized MET–iPSC interface as the hard gateway. **(g)** Hard bell-gene recovery. In serum, Palantir recovers a partially gateway-like transient program, whereas CellRank is sparse. In 2i/LIF, CellRank recovers a large but distinct stromal-ambiguity program and Palantir recovers a much smaller bell-gene set. **(h)** Median WOT backward transport probability to iPSC. Serum Palantir remains modestly fate-enriched, whereas CellRank and both 2i/LIF selections are weaker. **(i)** Cell-level overlap with the hard gateway. Palantir shows modest serum overlap, whereas both methods show little or no overlap with the 2i/LIF gateway. **(j)** Cross-condition bell-program coherence between serum and 2i/LIF. Gateway analysis shows the greatest cross-condition stability, with 15 shared bell genes overall and 4 shared genes among the top 20 bell genes, compared with 1 and 1 for Palantir and 0 and 0 for CellRank. Together, these comparisons show that established fate-mapping methods support the reality of the WOT transition corridor, whereas gateway analysis resolves the narrower MET–iPSC bottleneck and recovers the most stable transient program at that boundary.

**Extended Data Fig. 13.**
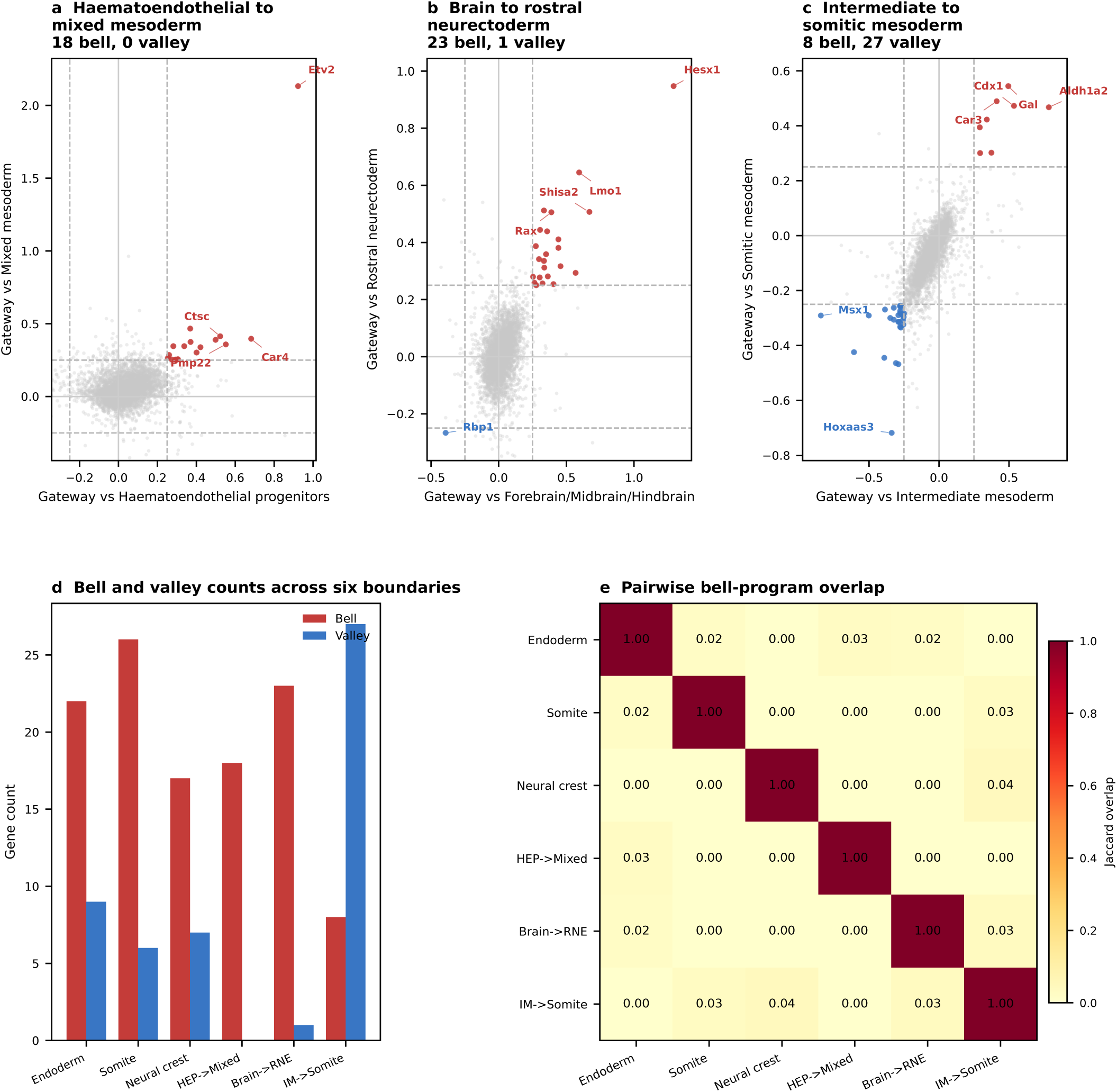
Additional gastrulation boundaries and cross-boundary analysis. **(a–c)** Four-quadrant scatter plots for three additional boundaries. **(a)** Haematoendothelial → Mixed Mesoderm, led by *Etv2*^59^. **(b)** Forebrain/Midbrain/Hindbrain → Rostral Neurectoderm, led by *Hesx1*^60^. **(c)** Intermediate Mesoderm → Somitic Mesoderm, where *Msx1* is a valley gene here but a gate gene at the neural-crest boundary, underscoring context-dependent TF roles. **(d)** Bell and valley counts across all six analyzed boundaries. **(e)** Pairwise bell-program overlap heatmap. Only 1–2 genes are shared between any pair, and none recur at more than two boundaries.

**Extended Data Fig. 14.**
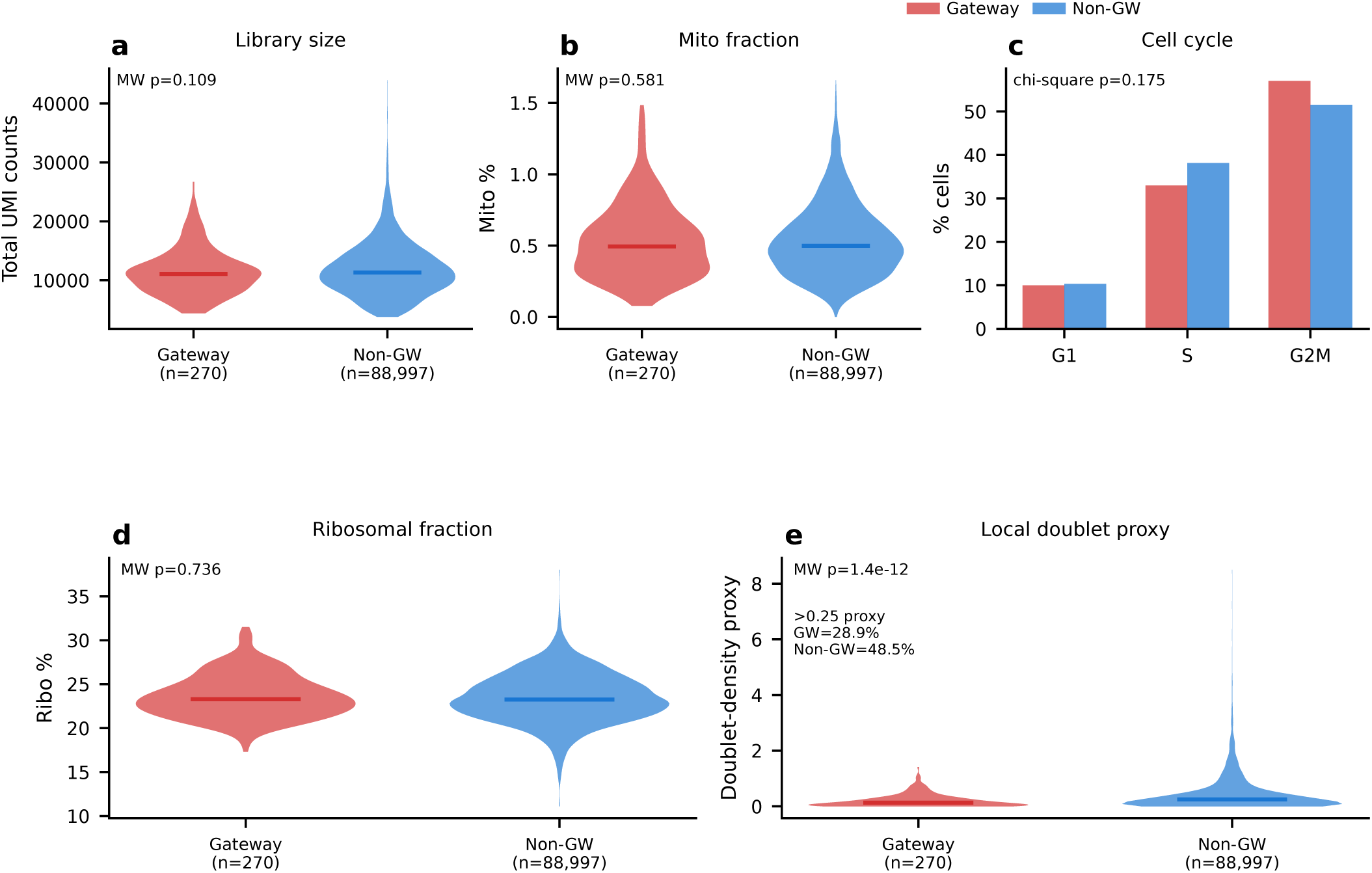
Technical confound controls for gastrulation gateway cells. Gateway cells (*n* = 270, union of 90 per focal boundary; red) versus non-gateway cells (*n* = 88,997; blue). **(a)** Total UMI counts (library size). **(b)** Mitochondrial read fraction. **(c)** Cell-cycle phase distribution (Tirosh et al.^61^). **(d)** Ribosomal read fraction. **(e)** Local doublet-density proxy derived from the doub.density field available in the localized gastrulation input. *P*-values: Mann–Whitney *U* (a,b,d,e) or chi-squared (c). Library size, mitochondrial fraction, ribosomal fraction, and cell-cycle composition show only small differences, while the local doublet proxy is lower in gateway cells than in non-gateway cells, arguing against doublet enrichment as the source of the signal.

**Extended Data Fig. 15.**
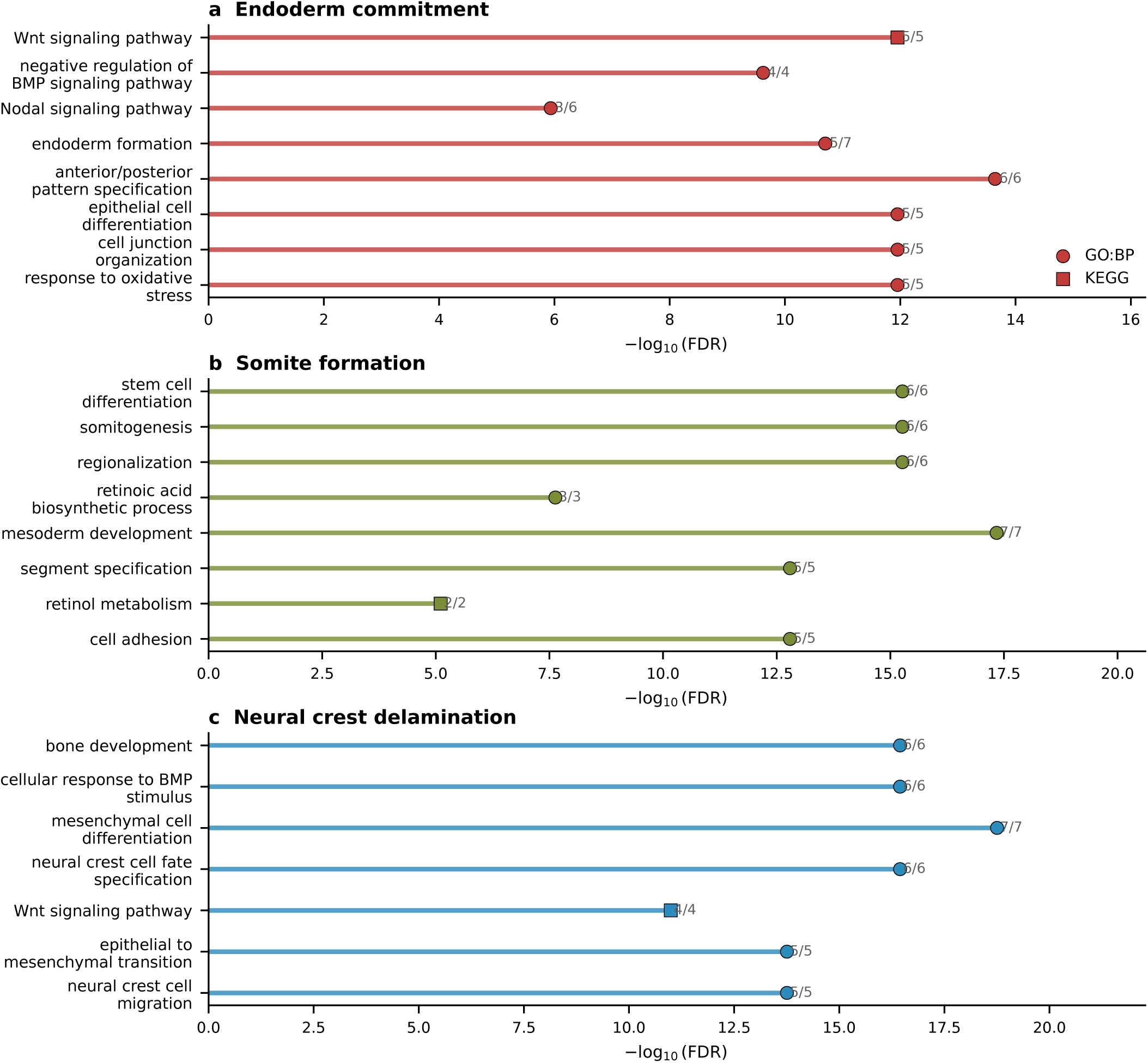
Gene Ontology enrichment of bell-gene programs. **(a–c)** Top enriched terms from the local enrichment panel. **(a)** Endoderm commitment: Wnt signaling, BMP regulation, and anterior-patterning programs. **(b)** Somite formation: stem-cell differentiation, somitogenesis, and regionalization. **(c)** Neural crest delamination: bone development, BMP response, and mesenchymal specification.

**Extended Data Fig. 16.**
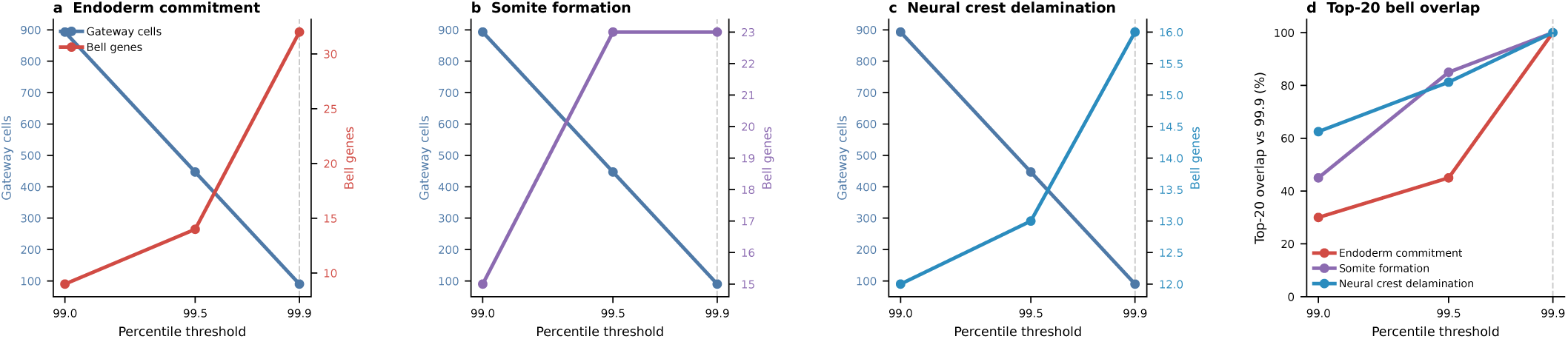
Threshold robustness of the three focal gastrulation boundaries. **(a–c)** Gateway-cell counts (blue) and bell-gene counts (colored) as the percentile threshold is relaxed from 99.9 to 99.5 and 99.0 for the APS→DE, Paraxial→Somitic, and Forebrain/Midbrain/Hindbrain→Neural Crest boundaries. Moderate relaxation expands the gateway pool while retaining much of the core bell program. **(d)** Top-20 bell-gene overlap relative to the selected threshold. The neural-crest and somitic boundaries preserve the strongest overlap under moderate relaxation, whereas the APS→DE boundary broadens more quickly. Together these analyses show that the gastrulation result is not tied to a single exact cutoff, although the most relaxed thresholds admit additional lower-amplitude genes.

**Extended Data Fig. 17.**
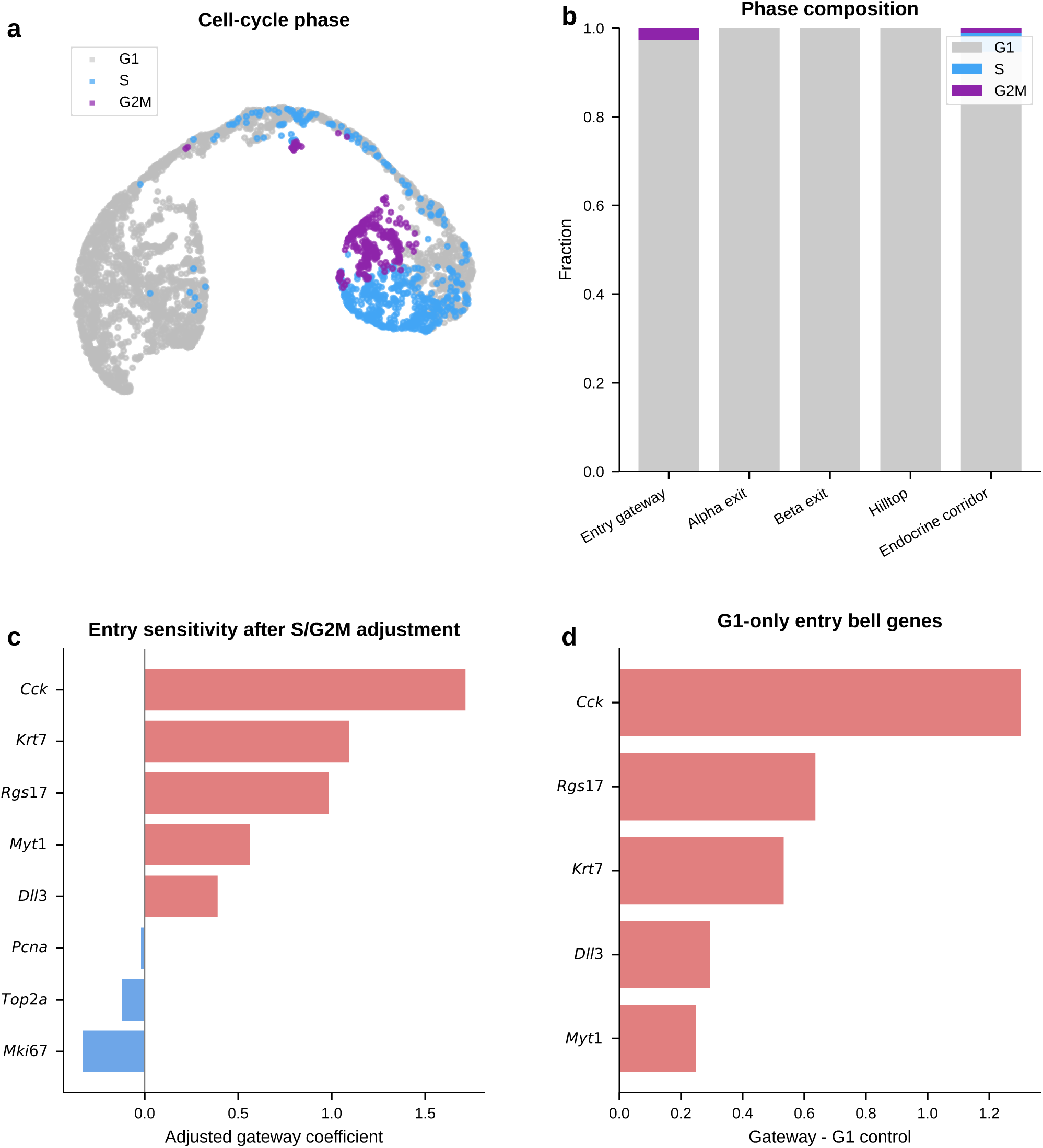
Cell-cycle control for pancreas hub entry. **(a)** UMAP of the pancreas trajectory colored by inferred cell-cycle phase using the existing S and G2/M scores from the trained embedding. The endocrine corridor already lies predominantly in G1. **(b)** Phase composition across the hub-entry and endocrine-exit populations. Entry gateway, exit gateway, and hilltop cells are not distinct cycling compartments relative to the surrounding endocrine corridor. **(c)** Entry-boundary sensitivity check after adjusting for continuous S and G2/M scores within the Ngn3 high EP/Pre-endocrine corridor. The leading bell genes (*Cck*, *Rgs17*, *Krt7*, *Dll3*, *Myt1*) retain positive gateway coefficients, whereas canonical cell-cycle controls (*Mki67*, *Pcna*, *Top2a*) do not. **(d)** Mean expression in entry gateway cells versus G1-only corridor controls. The same entry bell genes remain elevated when the comparison is restricted to other G1 cells, arguing that the entry gateway is not reducible to cell-cycle exit alone.

**Extended Data Fig. 18.**
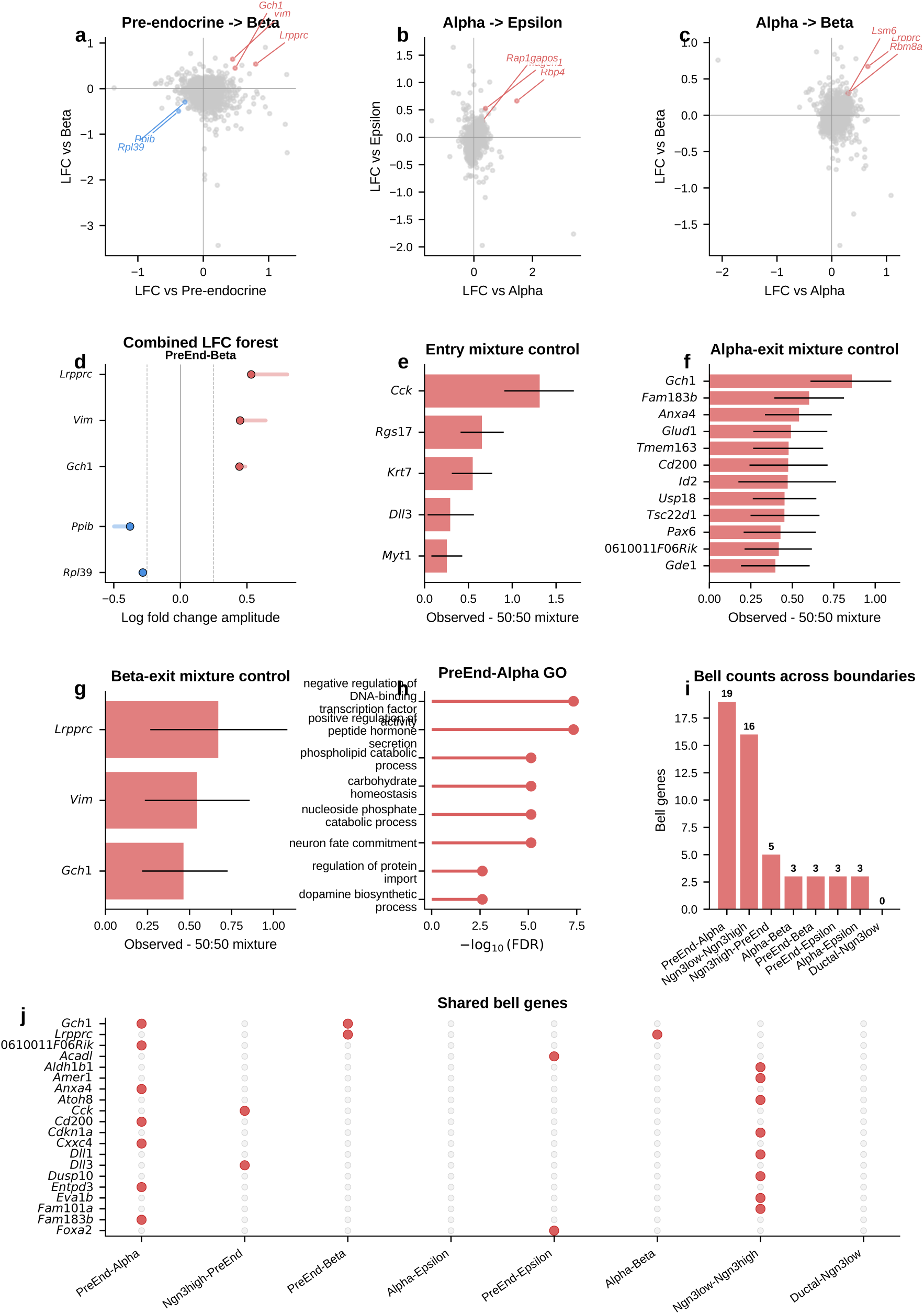
Additional pancreatic boundaries and controls supporting the shared endocrine hilltop program. **(a)** Four-quadrant LFC scatter for Pre-endocrine–Beta (3 bell, 2 valley). *Gch1* again appears as a bell gene, confirming that the candidate BH4-associated signal is not unique to the alpha exit. **(b)** Four-quadrant LFC scatter for Alpha–Epsilon (3 bell, 0 valley). Top genes include *Rbp4* and *Mageh1*, illustrating that other endocrine interfaces carry much sparser and different programs. **(c)** Four-quadrant LFC scatter for Alpha–Beta (3 bell, 0 valley), capturing a post-commitment interface between sibling hormone-producing fates and emphasizing the sparsity of the direct alpha-beta boundary relative to the shared pre-endocrine hilltop. **(d)** LFC amplitude forest plot for Pre-endocrine–Beta (3 bell + 2 valley genes), highlighting recurrence of *Gch1*. **(e)** Synthetic mixture control for Ngn3 high EP–Pre-endocrine bell genes. All 5 bell genes differ significantly from the synthetic 50:50 mixture (95% bootstrap CIs exclude zero; *n* = 2,000 resamples), supporting a bona fide gateway state rather than a simple mixture. **(f)** Synthetic mixture control for Pre-endocrine–Alpha bell genes. All 19 bell genes are significantly different from the mixture; the top 12 are shown. **(g)** Synthetic mixture control for Pre-endocrine–Beta bell genes, completing the mixture-control validation for the three main pancreatic gateway boundaries. **(h)** GO enrichment of Pre-endocrine–Alpha bell genes (top 8 terms). Top terms include neuron fate commitment, nucleoside phosphate catabolism, phospholipid catabolism, and positive regulation of peptide hormone secretion, linking the alpha-exit gateway to a transient neuroendocrine-like program. **(i)** Bell gene counts across all 8 scored pancreatic boundaries. Pre-endocrine–Alpha (19) has the richest program, whereas most other boundaries are much sparser, underscoring the special status of the alpha and beta exits from the hub. **(j)** Shared bell gene analysis (dot matrix). *Gch1* is shared between Pre-endocrine–Alpha and Pre-endocrine–Beta, identifying BH4 biosynthesis as a candidate shared endocrine transition program. *Lrpprc* is shared between Pre-endocrine–Beta and Alpha–Beta.

**Extended Data Fig. 19.**
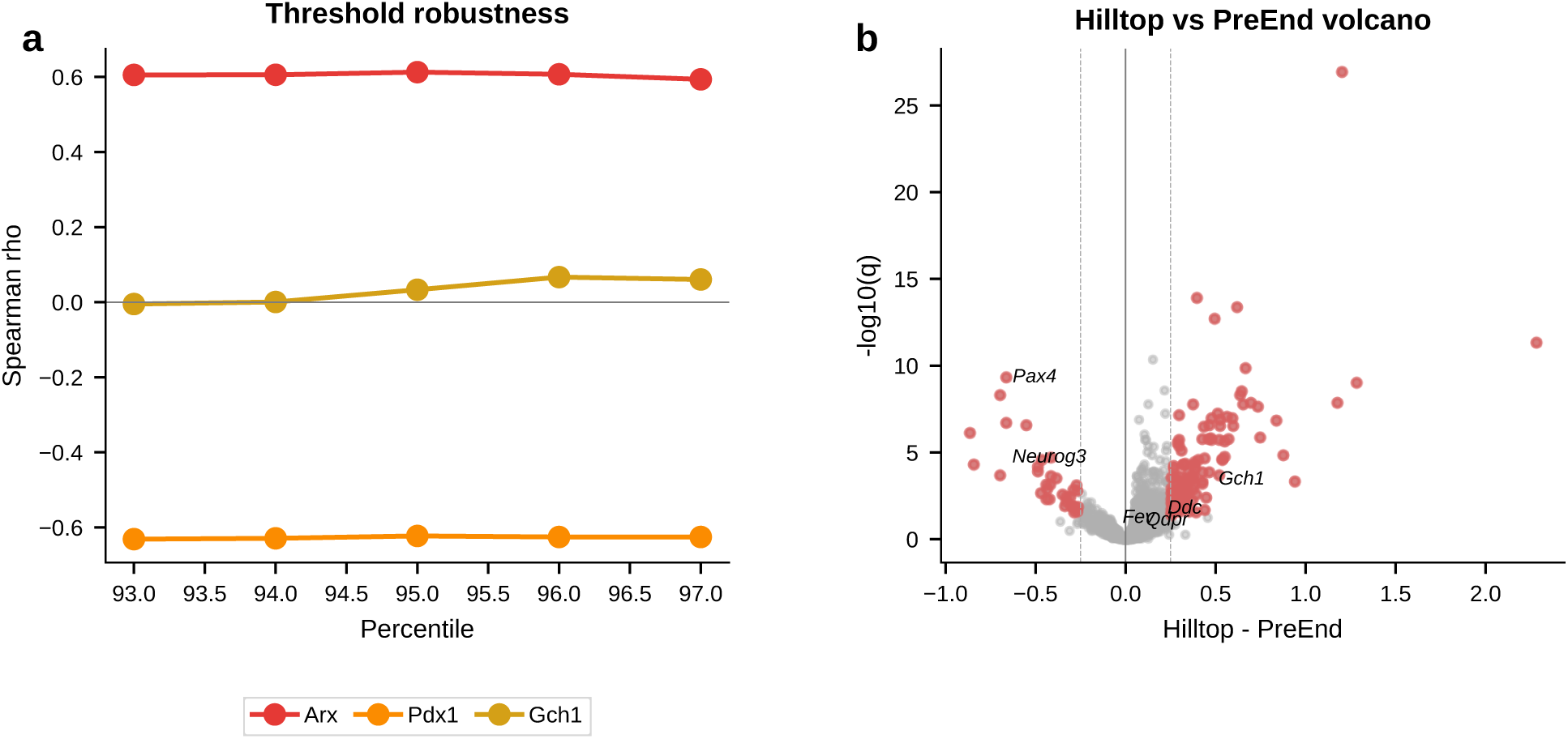
Hilltop robustness and orthogonal validation. **(a)** Threshold robustness. Spearman *ρ* of *Arx*, *Pdx1*, and *Gch1* with fate bias across 93rd–97th percentile thresholds. *Arx* (*ρ* ≈ +0.61) and *Pdx1* (*ρ* ≈ −0.62) remain stable across the tested range, whereas *Gch1* stays near the neutral zone (|*ρ*| < 0.1), showing that the separation between lineage-bias genes and shared hilltop genes is not driven by one specific cutoff. **(b)** Volcano plot: hilltop versus Pre-endocrine DE, confirming that the hilltop is a transcriptionally distinct state rather than simply a subset of immature pre-endocrine cells.

**Extended Data Fig. 20.**
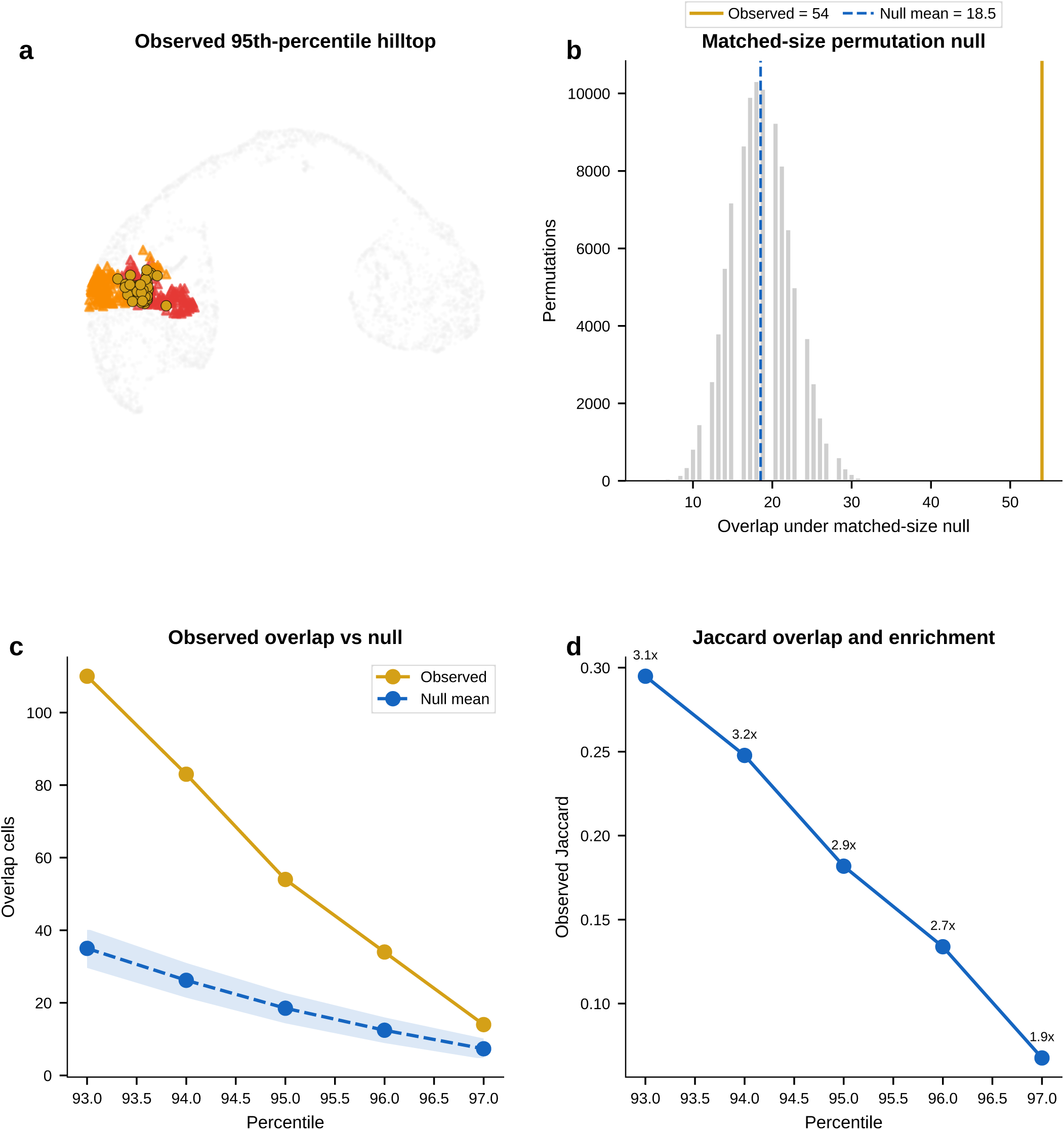
Matched-size permutation support for the pancreatic hilltop. **(a)** Observed hilltop overlap between the Pre-endocrine–Alpha and Pre-endocrine–Beta gateway sets at the 95th percentile. Hilltop cells (gold) sit at the shared precursor zone between the two exit-specific gateway cores. **(b)** Permutation null for the 95th-percentile hilltop, restricted to the endocrine corridor (Pre-endocrine, Alpha, and Beta cells) and matched to the observed alpha-exit and beta-exit gateway set sizes. The observed overlap (54 cells) exceeds the null expectation (18.5 cells; exact hypergeometric *P* = 1.9 × 10^−15^). **(c)** Observed overlap across 93rd–97th percentile thresholds compared with the matched-size null interval. Overlap remains above the null from the 93^rd^through 96th percentiles and narrows toward the null only at the ultra-stringent 97th-percentile core. **(d)** Jaccard overlap across the same threshold sweep, with enrichment relative to the matched-size null indicated above each point. The hilltop therefore represents a concentrated shared pre-commitment interval rather than the overlap expected from two independent gateway cores of the same size.

**Extended Data Fig. 21.**
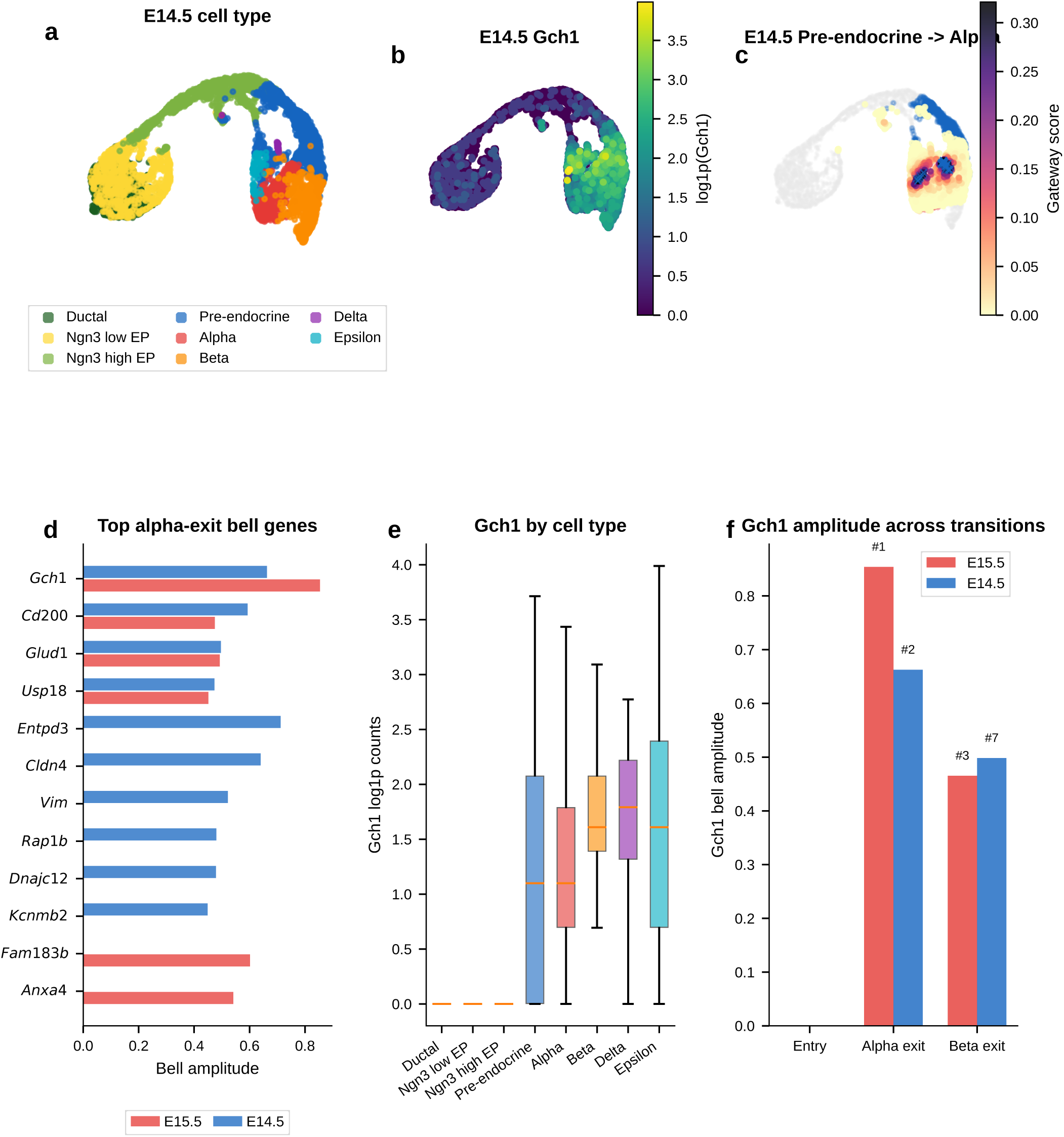
Temporal replication of the hilltop program at E14.5. Gateway analysis of 5,385 E14.5 endocrine-lineage cells from the same Bastidas-Ponce et al. 2019 study^4^, reprocessed locally from the combined E14.5/E15.5 atlas using the same scAttnVI and gateway pipeline as the main E15.5 analysis. **(a)** UMAP of the BMI-regularized latent space colored by cell type. The trajectory follows the same endocrine topology as E15.5 but contains proportionally more progenitors and fewer committed endocrine cells. **(b)** *Gch1* expression (log counts) overlaid on UMAP, showing strongest enrichment in the pre-endocrine and endocrine compartments. **(c)** Gateway cells (n = 54) at the Pre-endocrine–Alpha boundary, localized between Pre-endocrine and Alpha populations. **(d)** Top 10 bell genes at the Pre-endocrine–Alpha boundary at E14.5 (blue) compared with E15.5 (red). Four of the E15.5 top-10 bell genes (*Gch1*, *Cd200*, *Glud1*, and *Usp18*) recur within the E14.5 top 10, while *Entpd3* becomes the strongest E14.5 alpha-exit bell gene and *Gch1* remains second. **(e)** *Gch1* expression by cell type at E14.5. The local rerun preserves the hilltop-like profile: low in progenitors, highest in Pre-endocrine cells, and moderate in committed Alpha and Beta cells. **(f)** *Gch1* bell amplitude across the three pancreatic transitions at E15.5 (red) and E14.5 (blue). *Gch1* is absent at the upstream EP–Pre-endocrine boundary at both timepoints, present at both endocrine exits at both timepoints, strongest at the alpha exit at E15.5 (rank 1, amplitude 0.854), and weaker but still recovered at E14.5 (alpha rank 2, amplitude 0.663; beta rank 7, amplitude 0.498). Because both timepoints come from the same study, this result supports temporal stability rather than cross-study independence.

**Extended Data Fig. 22.**
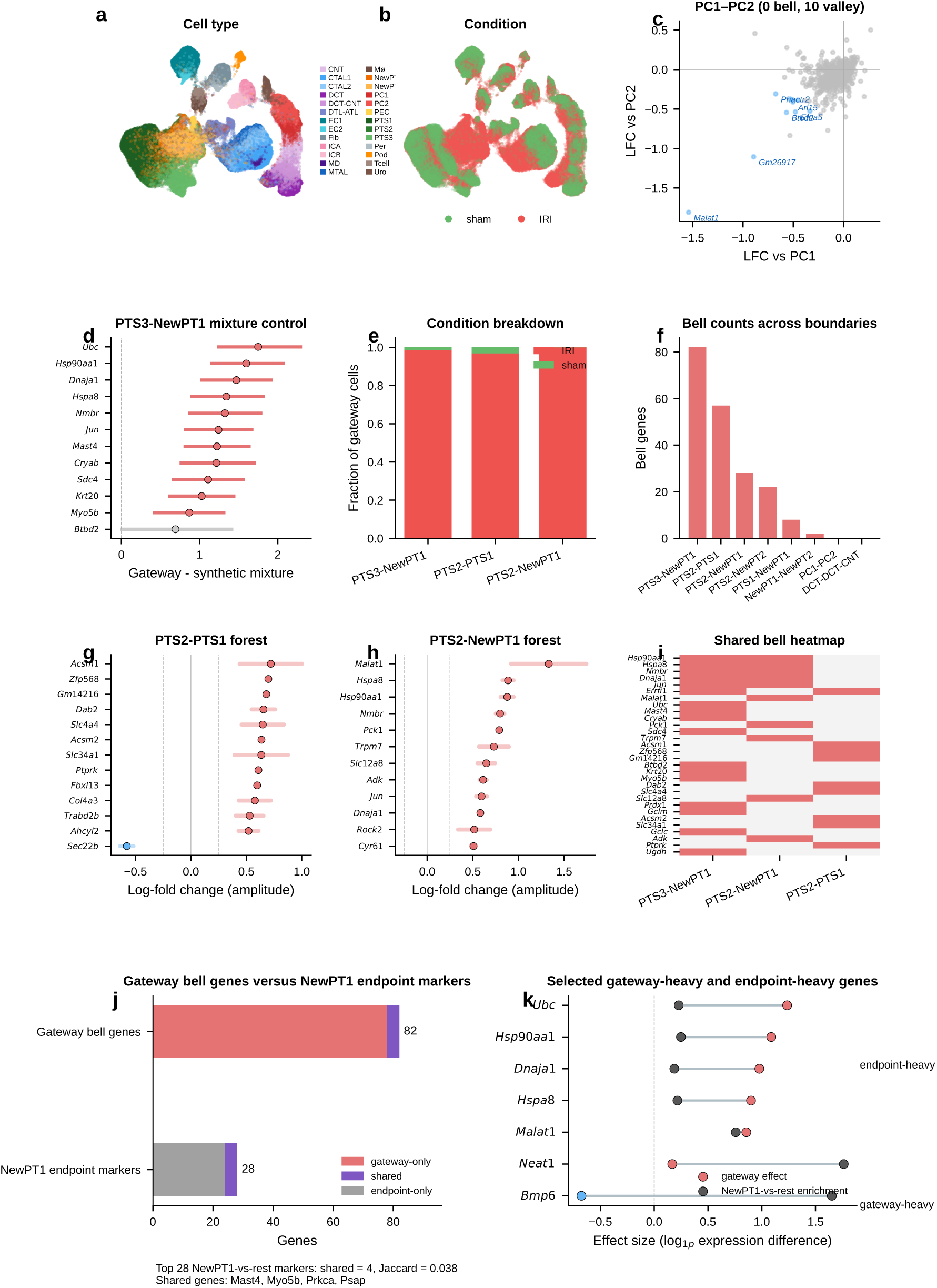
Additional kidney gateway boundaries and controls. **(a)** UMAP of the BMI-regularized latent space colored by cell type. Proximal-tubule subtypes include healthy segments (PTS1, PTS2, PTS3) and injury-associated populations (NewPT1, NewPT2). **(b)** UMAP colored by condition (sham versus IRI). NewPT1 and NewPT2 are dominated by IRI cells, while most other types contain cells from both conditions. **(c)** Four-quadrant LFC scatter for PC1–PC2 (0 bell, 10 valley genes). The absence of bell genes at this collecting-duct boundary indicates a transition dominated by program silencing rather than activation. **(d)** Synthetic mixture control for PTS3–NewPT1 bell genes. For each of the top 12 bell genes, horizontal bars show the mean expression difference (gateway − synthetic 50:50 PTS3/NewPT1 mixture) with 95% bootstrap CIs (*n* = 2,000). All 12 genes differ significantly from the mixture (CIs exclude zero), supporting a bona fide gateway state rather than a simple mixture. **(e)** Condition breakdown for the top 3 boundaries. Gateway cells are overwhelmingly IRI-derived (98.4–100.0%), consistent with injury-induced transitional states. **(f)** Bell gene counts across all 8 scored kidney boundaries, sorted by decreasing count. PTS3–NewPT1 (82) dominates; PTS2–PTS1 (57) and PTS2–NewPT1 (28) show substantial programs. **(g)** LFC amplitude forest plot for PTS2–PTS1 (top 12 of 57 bell genes plus the single valley gene), highlighting a transport- and maturation-focused program rather than the proteostasis module seen at injury entry. **(h)** LFC amplitude forest plot for PTS2–NewPT1 (top 12 of 28 bell genes). *Malat1* tops the list. **(i)** Shared bell gene heatmap across the three boundaries with the largest bell programs. Only 6 pairwise boundary–gene overlaps are detected, and no gene is shared across all three boundaries. **(j)** Overlap between the PTS3–NewPT1 gateway bell set and the top 28 NewPT1-vs-rest endpoint markers is minimal (4 shared genes out of 82 gateway bell genes; Jaccard 0.038), supporting a transient injury-entry state rather than simple relabeling of established endpoint identity. **(k)** Selected genes show that gateway-heavy markers such as *Ubc*, *Hsp90aa1*, *Dnaja1*, and *Hspa8* have large gateway effects but weaker NewPT1-vs-rest enrichment, whereas *Neat1* and *Bmp6* are strongly endpoint-skewed and *Malat1* remains elevated in both comparisons.

**Extended Data Fig. 23.**
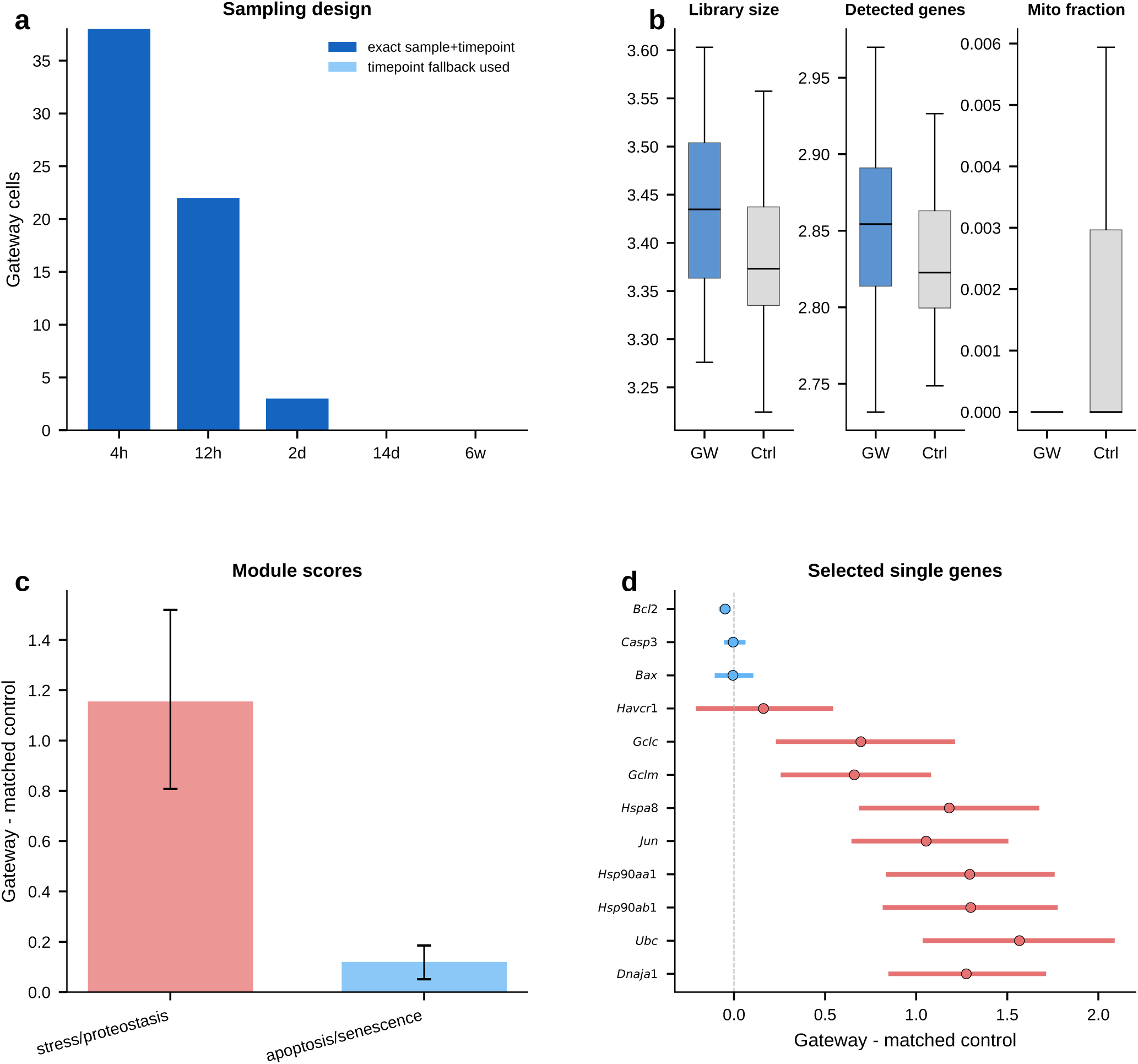
Kidney gateway cells versus time-matched PTS3 controls. **(a)** Sampling design. Each IRI gateway cell at the PTS3–NewPT1 boundary was paired with five non-gateway PTS3 controls from the same IRI sample and time point whenever possible. Gateway cells are concentrated at early injury time points (4 h and 12 h). **(b)** QC shifts for gateway cells relative to time-matched controls. Library size and detected genes are modestly higher in gateway cells, whereas mitochondrial fraction is not elevated. **(c)** Module scores. A generic stress/proteostasis module remains enriched in gateway cells, whereas the interpretation should not rely on a broad apoptosis program alone. **(d)** Selected single genes. *Ubc*, *Hsp90aa1*, *Hsp90ab1*, *Dnaja1*, *Hspa8*, *Jun*, *Gclm*, and *Gclc* remain elevated relative to time-matched PTS3 controls, whereas *Havcr1* is no longer distinctive and classical apoptosis markers (*Casp3*, *Bax*, *Bcl2*) remain weak. These results support a temporally localized injury-entry interval rather than simple endpoint identity or a gross dying-cell compartment.

## References

1. Waddington, C.H. (1957). The Strategy of the Genes (Allen & Unwin).

2. Moris, N., Pina, C., and Arias, A.M. (2016). Transition states and cell fate decisions in epigenetic landscapes. Nat. Rev. Genet. 17, 693–703.

3. Schiebinger, G., Shu, J., Tabaka, M., Cleary, B., Subramanian, V., Solomon, A., Gould, J., Liu, S., Lin, S., Berber, P., et al. (2019). Optimal-transport analysis of single-cell gene expression identifies developmental trajectories in reprogramming. Cell 176, 928–943.

4. Bastidas-Ponce, A., Tritschler, S., Dony, L., Scheber, K., Tarquis-Medina, M., Salinno, C., Schirge, S., Burtscher, I., Böttcher, A., Theis, F.J., et al. (2019). Comprehensive single cell mRNA profiling reveals a detailed roadmap for pancreatic endocrinogenesis. Development 146, dev173849.

5. Kiselev, V.Y., Andrews, T.S., and Hemberg, M. (2019). Challenges in unsupervised clustering of single-cell RNA-seq data. Nat. Rev. Genet. 20, 273–282.

6. Lopez, R., Regier, J., Cole, M.B., Jordan, M.I., and Yosef, N. (2018). Deep generative modeling for single-cell transcriptomics. Nat. Methods 15, 1053–1058.

7. Trapnell, C., Cacchiarelli, D., Grimsby, J., Pokharel, P., Li, S., Morse, M., Lennon, N.J., Livak, K.J., Mikkelsen, T.S., and Rinn, J.L. (2014). The dynamics and regulators of cell fate decisions are revealed by pseudotemporal ordering of single cells. Nat. Biotechnol. 32, 381–386.

8. La Manno, G., Soldatov, R., Zeisel, A., Braun, E., Hochgerner, H., Petukhov, V., Lidschreiber, K., Kastriti, M.E., Lönnerberg, P., Furlan, A., et al. (2018). RNA velocity of single cells. Nature 560, 494–498.

9. Lange, M., Bergen, V., Klein, M., Setty, M., Haghverdi, L., Buttner, M., Moez, M., Peidli, S., Wolf, F.A., Murray, I., et al. (2022). CellRank for directed single-cell fate mapping. Nat. Methods 19, 159–170.

10. Pĳuan-Sala, B. et al. (2019). A single-cell molecular map of mouse gastrulation and early organogenesis. Nature 566, 490–495.

11. Kirita, Y., Wu, H., Uchimura, K., Wilson, P.C., and Bhatt, H. (2020). Cell profiling of mouse acute kidney injury reveals conserved cellular responses to injury. Proc. Natl. Acad. Sci. USA 117, 15874–15883.

12. Zheng, G.X.Y., Terry, J.M., Belgrader, P., Ryvkin, P., Bent, Z.W., Wilson, R., Ziraldo, S.B., Wheeler, T.D., McDermott, G.P., Zhu, J., et al. (2017). Massively parallel digital transcriptional profiling of single cells. Nat. Commun. 8, 14049.

13. Ying, Q.-L., Wray, J., Nichols, J., Batlle-Morera, L., Doble, B., Woodgett, J., Cohen, P., and Smith, A. (2008). The ground state of embryonic stem cell self-renewal. Nature 453, 519–523.

14. Page, L., Brin, S., Motwani, R., and Winograd, T. (1999). The PageRank citation ranking: Bringing order to the web. Stanford InfoLab Technical Report.

15. Cowen, L., Ideker, T., Raphael, B.J., and Sharan, R. (2017). Network propagation: a universal amplifier of genetic associations. Nat. Rev. Genet. 18, 551–562.

16. Signer, R.A.J., Magee, J.A., Salic, A., and Morrison, S.J. (2014). Haematopoietic stem cells require a highly regulated protein synthesis rate. Nature 509, 49–54.

17. Sakurai, K., Talukdar, I., Patil, V.S., Dang, J., Li, Z., Chang, K.-Y., Lu, C.-C., Sber, N., Bhatt, D.M., and Bhatt, D.M. (2014). Kinome-wide functional analysis highlights the role of cytoskeletal remodeling in somatic cell reprogramming. Cell Rep. 7, 1085–1095.

18. Li, R., Liang, J., Ni, S., Zhou, T., Qing, X., Li, H., He, W., Chen, J., Li, F., Zhuang, Q., et al. (2010). A mesenchymal-to-epithelial transition initiates and is required for the nuclear reprogramming of mouse fibroblasts. Cell Stem Cell 7, 51–63.

19. Samavarchi -Tehrani, P., Golipour, A., David, L., Sung, H.-K., Beyer, T.A., Datti, A., Woltjen, K., Nagy, A., and Wrana, J.L. (2010). Functional genomics reveals a BMP-driven mesenchymal-to-epithelial transition in the initiation of somatic cell reprogramming. Cell Stem Cell 7, 64–77.

20. Zhou, G., Meng, S., Li, Y., Ghebre, Y.T., and Bhatt, D.M. (2016). Optimal ROS signaling is critical for nuclear reprogramming. Cell Rep. 15, 919–925.

21. Kolodziejczyk, A.A., Kim, J.K., Tsang, J.C.H., Ilicic, T., Henriksson, J., Natarajan, K.N., Tuck, A.C., Gao, X., Bühler, M., Liu, P., et al. (2015). Single cell RNA-sequencing of pluripotent states unlocks modular transcriptional variation. Cell Stem Cell 17, 471–485.

22. Wolock, S.L., Lopez, R., and Klein, A.M. (2019). Scrublet: computational identification of cell doublets in single-cell transcriptomic data. Cell Syst. 8, 281–291.

23. Armstrong, L., Tilgner, K., Saretzki, G., Atkinson, S.P., Stojkovic, M., Moreno, R., Przyborski, S., and Lako, M. (2010). Human induced pluripotent stem cell lines show stress defense mechanisms and mitochondrial regulation similar to those of human embryonic stem cells. Stem Cells 28, 661–673.

24. Setty, M., Kiseliovas, V., Levine, J., Gayoso, A., Mazutis, L., and Pe’er, D. (2019). Characterization of cell fate probabilities in single-cell data with Palantir. Nat. Biotechnol. 37, 451–460.

25. Pourquié, O. Vertebrate segmentation: from cyclic gene networks to scoliosis. Cell 145, 650–663 (2011).

26. Simoes-Costa, M. & Bronner, M. E. Establishing neural crest identity: a gene regulatory recipe. Development 142, 242–257 (2015).

27. Sauka-Spengler, T. & Bronner-Fraser, M. A gene regulatory network orchestrates neural crest formation. Nat. Rev. Mol. Cell Biol. 9, 557–568 (2008).

28. Davidson, E. H. & Erwin, D. H. Gene regulatory networks and the evolution of animal body plans. Science 311, 796–800 (2006).

29. Niederreither, K. et al. Retinoic acid synthesis and hindbrain patterning in the mouse embryo. Development 127, 75–85 (2000).

30. Duester, G. Retinoic acid synthesis and signaling during early organogenesis. Cell 134, 921–931 (2008).

31. Kanai -Azuma, M. et al. Depletion of definitive gut endoderm in *Sox17*-null mutant mice. Development 129, 2367–2379 (2002).

32. Arnold, S. J. et al. Pivotal roles for eomesodermin during axis formation, epithelium-to-mesenchyme transition and endoderm specification in the mouse. Development 135, 501–511 (2008).

33. Burgess, R., Rawls, A., Tjian, R. & Olson, E. N. *Paraxis*: a basic helix-loop-helix protein expressed in paraxial mesoderm and developing somites. Dev. Biol. 168, 296–306 (1995).

34. Mankoo, B. S. et al. The concerted action of *Meox* homeobox genes is required upstream of genetic pathways essential for the formation, patterning and differentiation of somites. Development 130, 4655–4664 (2003).

35. Kume, T., Jiang, H., Topczewska, J. M. & Hogan, B. L. M. The murine winged helix transcription factors, Foxc1 and Foxc2, are both required for cardiovascular development and somitogenesis. Genes Dev. 15, 2470–2482 (2001).

36. Seo, K. W. et al. Targeted disruption of the DM domain containing transcription factor gene *Dmrt2* reveals an essential role in somite patterning. Dev. Biol. 290, 200–210 (2006).

37. Bessho, Y. et al. Dynamic expression and essential functions of Hes7 in somite segmentation. Genes Dev. 15, 2642–2647 (2001).

38. de Crozé, N., Maczkowiak, F. & Monsoro-Burq, A. H. Reiterative AP2a activity controls sequential steps in the neural crest gene regulatory network. Proc. Natl Acad. Sci. USA 108, 155–160 (2011).

39. Cheung, M. & Briscoe, J. Neural crest development is regulated by the transcription factor Sox9. Development 130, 5681–5693 (2003).

40. Mundell, N. A. & Labosky, P. A. Neural crest stem cell multipotency requires Foxd3 to maintain neural potential and repress mesenchymal fates. Development 138, 641–652 (2011).

41. Tribulo, C. et al. Regulation of *Msx* genes by a Bmp gradient is essential for neural crest specification. Development 130, 6441–6452 (2003).

42. McMahon, A. P. & Bradley, A. The Wnt-1 (*int*-1) proto-oncogene is required for development of a large region of the mouse brain. Cell 62, 1073–1085 (1990).

43. Depew, M. J. et al. Dlx5 regulates regional development of the branchial arches and sensory capsules. Development 126, 3831–3846 (1999).

44. Dunty, W. C. et al. Wnt3a/*β*-catenin signaling controls posterior body development by coordinating mesoderm formation and segmentation. Development 135, 85–94 (2008).

45. Relaix, F. et al. A Pax3/Pax7-dependent population of skeletal muscle progenitor cells. Nature 435, 948–953 (2005).

46. Yokota, Y. Id and development. Oncogene 20, 8290–8298 (2001).

47. Southard-Smith, E. M., Kos, L. & Pavan, W. J. Sox10 mutation disrupts neural crest development in Dom Hirschsprung mouse model. Nat. Genet. 18, 60–64 (1998).

48. Ang, S. L. et al. The formation and maintenance of the definitive endoderm lineage in the mouse: involvement of HNF3/forkhead proteins. Development 119, 1301–1315 (1993).

49. Morrisey, E. E. et al. GATA6 regulates HNF4 and is required for differentiation of visceral endoderm in the mouse embryo. Genes Dev. 12, 3579–3590 (1998).

50. Lee, C. S. et al. Foxa2 is required for the differentiation of pancreatic *⍺*-cells. Dev. Biol. 278, 484–495 (2005).

51. Sicinski, P. et al. Cyclin D2 is an FSH-responsive gene involved in gonadal cell proliferation and oncogenesis. Nature 384, 470–474 (1996).

52. Spitz, F. & Furlong, E. E. M. Transcription factors: from enhancer binding to developmental control. Nat. Rev. Genet. 13, 613–626 (2012).

53. Levine, M. & Davidson, E. H. Gene regulatory networks for development. Proc. Natl Acad. Sci. USA 102, 4936–4942 (2005).

54. Stadhouders, R., Filber, G. J. & Graf, T. Transcription factors and 3D genome conformation in cell-fate decisions. Nature 569, 345–354 (2019).

55. Wolf, F.A., Angerer, P., and Theis, F.J. (2018). SCANPY: large-scale single-cell gene expression data analysis. Genome Biol. 19, 15.

56. Hansen, P.C. (1992). Analysis of discrete ill-posed problems by means of the L-curve. SIAM Rev. 34, 561–580.

57. Hansen, P.C. and O’Leary, D.P. (1993). The use of the L-curve in the regularization of discrete ill-posed problems. SIAM J. Sci. Comput. 14, 1487–1503.

58. Raudvere, U. et al. g:Profiler: a web server for functional enrichment analysis and conversions of gene lists (2019 update). Nucleic Acids Res. 47, W191–W198 (2019).

59. Koy ano-Nakagawa, N. et al. Etv2 as a master regulator of hematoendothelial lineage specification. Cell Rep. 1, 1–11 (2012).

60. Dattani, M. T. et al. Mutations in the homeobox gene HESX1/Hesx1 associated with septo-optic dysplasia in human and mouse. Nat. Genet. 19, 125–133 (1998).

61. Tirosh, I. et al. Dissecting the multicellular ecosystem of metastatic melanoma by single-cell RNA-seq. Science 352, 189–196 (2016).

62. Tikhonov, A.N. and Arsenin, V.Y. (1977). Solutions of Ill-Posed Problems (Winston & Sons).

63. Scavuzzo, M. A. et al. Endocrine lineage biases arise in temporally distinct endocrine progenitors during pancreatic morphogenesis. Nat. Commun. 9, 3356 (2018).

64. Ghoshal, K., Wang, Y., Sheridan, J. F. & Jacob, S. T. Metallothionein induction in response to restraint stress: transcriptional control, adaptation to stress, and role of glucocorticoid. J. Biol. Chem. 273, 27904–27910 (1998).

65. Deane, C. A. S. & Brown, I. R. Components of a mammalian protein disaggregation/refolding machine are targeted to nuclear speckles following thermal stress in differentiated human neuronal cells. Cell Stress Chaperones 22, 191–200 (2017).

66. Bansal, G. S., Norton, P. M. & Latchman, D. S. The 90-kDa heat shock protein protects mammalian cells from thermal stress but not from viral infection. Exp. Cell Res. 195, 303–306 (1991).

67. Relaix, F. et al. Pw1/Peg3 is a potential cell death mediator and cooperates with Siah1a in p53-mediated apoptosis. Proc. Natl Acad. Sci. USA 97, 2105–2110 (2000).

68. Jurk, D. et al. Postmitotic neurons develop a p21-dependent senescence-like phenotype driven by a DNA damage response. Aging Cell 11, 996–1004 (2012).

69. Oltvai, Z. N., Milliman, C. L. & Korsmeyer, S. J. Bcl-2 heterodimerizes in vivo with a conserved homolog, Bax, that accelerates programmed cell death. Cell 74, 609–619 (1993).

70. Yu, J., Zhang, L., Hwang, P. M., Kinzler, K. W. & Vogelstein, B. PUMA induces the rapid apoptosis of colorectal cancer cells. Mol. Cell 7, 673–682 (2001).

71. Tomasini, R. et al. TP53INP1 is a novel p73 target gene that induces cell cycle arrest and cell death by modulating p73 transcriptional activity. Oncogene 24, 8093–8104 (2005).

72. Hildesheim, J. et al. Gadd45a protects against UV irradiation-induced skin tumors, and promotes apoptosis and stress signaling via MAPK and p53. Cancer Res. 62, 7305–7315 (2002).

73. Wajapeyee, N., Serra, R. W., Zhu, X., Mahalingam, M. & Green, M. R. Oncogenic BRAF induces senescence and apoptosis through pathways mediated by the secreted protein IGFBP7. Cell 132, 363–374 (2008).

74. Bianchi, M., Giacomini, E., Crinelli, R., Radici, L., Carloni, E. & Magnani, M. Dynamic transcription of ubiquitin genes under basal and stressful conditions and new insights into the multiple UBC transcript variants. Gene 573, 100–109 (2015).

75. Ueyama, T., Umemoto, S. & Senba, E. Immobilization stress induces c-fos and c-jun immediate early genes expression in the heart. Life Sci. 59, 339–347 (1996).

76. Solis, W. A. et al. Glutamate-cysteine ligase modifier subunit: mouse Gclm gene structure and regulation by agents that cause oxidative stress. Biochem. Pharmacol. 63, 1739–1754 (2002).

77. Bea, F. et al. Homocysteine stimulates antioxidant response element-mediated expression of glutamate-cysteine ligase in mouse macrophages. Atherosclerosis 203, 105–111 (2009).

78. Slee, E. A., Adrain, C. & Martin, S. J. Executioner caspase-3, -6, and -7 perform distinct, non-redundant roles during the demolition phase of apoptosis. J. Biol. Chem. 276, 7320–7326 (2001).

